# HDAC4 controls senescence and aging by safeguarding the epigenetic identity and ensuring the genomic integrity

**DOI:** 10.1101/2020.06.04.132787

**Authors:** Eros Di Giorgio, Harikrishnareddy Paluvai, Emiliano Dalla, Liliana Ranzino, Alessandra Renzini, Viviana Moresi, Valentina Cutano, Raffaella Picco, Claudio Brancolini

**Affiliations:** Department of Medicine, Università degli Studi di Udine, p.le Kolbe 4, 33100 Udine, Italy; DAHFMO Unit of Histology and Medical Embryology, Sapienza University of Rome, via Antonio Scarpa 16, 00161, Rome, Italy

## Abstract

The epigenome of senescent cells is characterized by a deep redistribution of H3K27 acetylation. H3K27 is target of class IIa Histone Deacetylases (HDAC4, 5, 7, 9) as part of large repressive complexes. We report here that, among class IIa HDACs, HDAC4 is post-transcriptionally downregulated during senescence and aging. HDAC4 knock-out (KO) triggers premature senescence as a result of two waves of biological events: the accumulation of replication stress (RS) and the expression of inflammatory genes. The latter is achieved directly, through the activation of enhancers (TEs) and super-enhancers (SEs) that are normally monitored by HDAC4, and indirectly, through the de-repression of repetitive elements of retroviral origin (ERVs). The accumulation of DNA damage and the activation of the inflammatory signature influence each other and integrate into a synergistic response required for senescence onset. Our work discloses the key role played by HDAC4 in maintaining epigenome identity and genome integrity.

## INTRODUCTION

Cellular senescence and aging are complex responses characterized by proliferative arrest and loss of regenerative potential^1^. The senescence state is distinguished by a deep epigenetic reprogramming that sculptures the chromatin to maintain cellular survival, arrest the cell-cycle and secrete apocrine and paracrine factors in the presence of consistent DNA damage^2^. In turn, the haploinsufficiency of epigenetic and metabolic regulators increases genome fragility, affects the DNA damage response (DDR) and predisposes to senescence, cell death or malignant transformation^3–7^.

H3K27ac/H3K27me3 ratio alterations were identified as the driving forces of premature senescence and cancer^3,7–10^. The balanced action of acetyltransferases (HAT) of the SWI/SNF/p300 complex^11^ and HDACs of the Sin3, NuRD, CoREST, MiDAC and NCOR complexes^12^ controls the acetylation status of H3K27. Class IIa HDACs are catalytically inactive epigenetic readers, quickly recruited on H3K27ac loci^13,14^, where they monitor the acetylation status through the binding of class I HDACs^15^.

Here we investigated the role played by class IIa HDACs in the regulation of senescence entrance. The senescence phenotype induced by HDAC4 depletion was studied in detail to unveil the dual role of this epigenetic regulator in preserving genome integrity and in safeguarding regulative elements controlling cell fate.

## RESULTS

### HDAC4 expression is silenced during cellular senescence and aging

The epigenetic reprogramming plays key roles in establishing cellular senescence and aging. In this context the contribution of class IIa HDACs has been suggested^16,17^, but not addressed in a comprehensive manner. Important epigenetic modulators of senescence and aging are down-regulated during the progressive cell cycle arrest^3,4,18,19^. For this reason, we evaluated class IIa HDACs expression in different models of senescence and aging. HDAC4 and HDAC9, and to a lesser extent HDAC7, were progressively downregulated in human IMR90 fibroblasts undergoing replicative senescence (Fig. 1a/b). As a model of ageing, we compared the protein levels of class IIa HDACs in the dermis and in the liver of young (4 months) and old (25 months) mice. HDAC4 and HDAC9 levels decreased, both in aged dermis and liver, whereas HDAC5 decreased only in old dermis (Fig. 1c). Therefore, telomers attrition in normal cells and physiological aging similarly affected the protein levels of HDAC4 and HDAC9.

**Figure 1.**
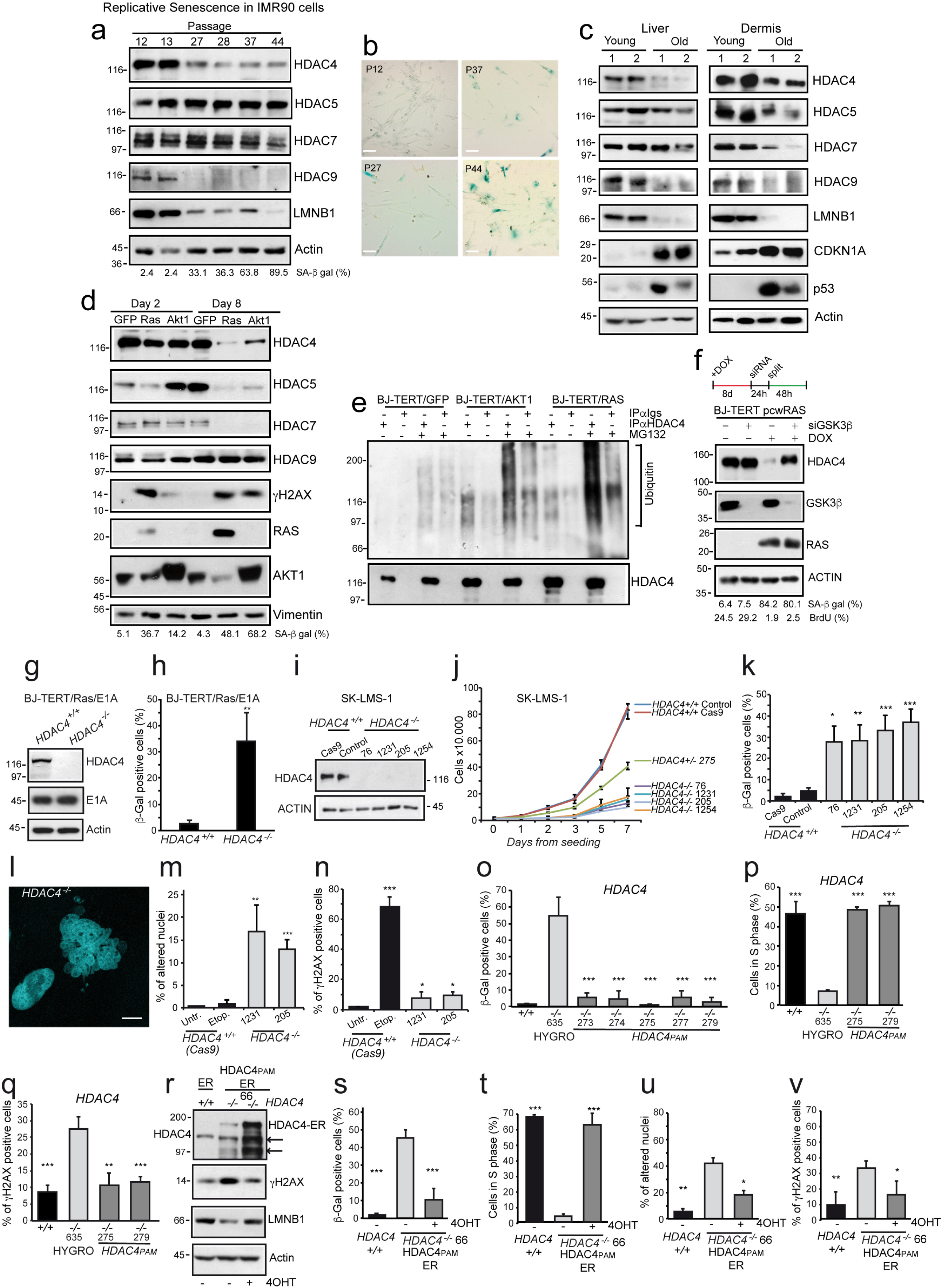
HDAC4 is dysregulated during senescence and aging and is required for senescence escape. **a.** Immunoblot analysis in IMR90 cells undergoing replicative senescence. Actin was used as loading control. **b.** Microscopic images of SA-β-gal stained IMR90 cells (scale bar 50 µm). **c.** Immunoblot analysis in tissue-derived lysates obtained from C57BL/6J female mice sacrificed at 128 (young) and 774 (old) days of age. Actin was used as loading control. **d.** Immunoblot analysis in BJ/*hTERT* expressing the indicated transgenes for the indicated time. Vimentin was the loading control. **e.** Cellular lysates obtained in BJ/*hTERT* expressing for 8 days the indicated transgenes and treated or not for 8h with MG132 were immunoprecipitated using anti-HDAC4 and immunoblotted with the indicated antibodies. **f.** Immunoblot analysis in BJ/*hTERT* cells expressing HRAS^G12V^ and silenced for GSK3β, as indicated. **g.** Immunoblot analysis in BJ/*hTERT/E1A/RAS*/*HDAC4*^+*/*+^ or ^*-/-*^ cells, as indicated. Actin was used as loading control. **h.** Analysis of the senescent cells as scored after SA-β-gal staining. Mean ± SD; n = 3. **i.** Immunoblot analysis in SK-LMS-1/*HDAC4*^+*/*+^ or ^*-/-*^ as indicated. **j.** Cell-proliferation curve of the indicated *HDAC4*^+*/*+, +*/-, -/-*^ SK-LMS-1 cells. Mean ± SD; n = 4. **k.** Analysis of the senescent cells as scored after SA-β-gal staining. Mean ± SD; n = 4. **l.** Representative image of normal and altered DAPI-stained nuclei observed in SK-LMS-1 *HDAC4*^*-/-*^ cells (scale bar 10 µm). **m-n.** Analysis of the % of cells displaying altered nuclei or γH2AX foci (>5). Etoposide was a control (2h, 20µM). Mean ± SD; n = 4. **o-p-q.** Analysis of SA-β-gal (o), BrdU (p) and γH2AX (q) positivity in wt or HDAC4 KO cells expressing HYGRO^R^ (clone 635) or HDAC4^PAM^. Mean ± SD; n = 4. The significance is relative to clone 635. **r.** Immunoblot analysis in SK-LMS-1 wt or KO (clone 66) cells re-expressing a tamoxifen inducible *HDAC4*^*PAM*^*-ER*. Arrowheads point to HDAC4 cleavage products observed in HDAC4-ER expressing cells. Actin was the loading control. **s-v.** Analysis of SA-β-gal (s), BrdU (t), nuclear alteration (u) and γH2AX (v) positivity in the indicated cells. Mean ± SD; n = 4.

The delivery in BJ/*hTERT* of the strong oncogenes HRAS^G12V^ or myrAKT1 triggers a premature cell-cycle arrest named oncogene-induced senescence (OIS)^20,21^. In both models of OIS (Fig.S1a/b), the permanent proliferation arrest is characterized by the post-transcriptional downregulation of HDAC4, HDAC5 and HDAC7, but not of HDAC9 (Fig.1d, S1c,d,h). This trend was also observed in IMR90/*RAS* cells (Fig. S1e).

Other oncogenes of viral origin, like E1A and its ΔC-fragment (1-143), are able to overcome OIS, by targeting p16/Rb pathways^22^. HDAC4/5/9, but not HDAC7, were upregulated by the expression of E1AΔC in BJ/*hTERT* (Fig.S1c). Moreover, the co-expression of E1A and RAS by-passed the senescent arrest and recovered the protein levels of HDAC4 and HDAC5, but not of HDAC7 (Fig.S1c). Therefore, OIS and OIS escape affect mainly HDAC4 and HDAC5 levels. HDAC4 and HDAC5 were decreased also during stress-induced senescence (SISP) following H_2_O_2_ treatment and cytokines-induced senescence (Fig.S1f,g,i). In all these conditions, class IIa HDACs are regulated at the post-transcriptional level, except for HDAC9, which transcription is stimulated by oncogenes (Fig. S1c and Table S1).

Our preliminary screening identified HDAC4 as the class IIa HDAC member repressed in all tested models of senescence and aging. Most of this downregulation is due to the ubiquitin-proteasome system (UPS) mediated degradation. HDAC4 levels in senescent cells were restored after UPS, but not autophagy inhibition (Fig. S2a/b). Accordingly, HDAC4 was highly poly-ubiquitylated in senescent cells (Fig. 1e). This degradation requires GSK3β^23^, as the treatment with LiCl (Fig. S2c) and the silencing of GSK3β restored HDAC4 levels in senescent cells (Fig.1f).

### The depletion of HDAC4 triggers senescence in different cell types

To investigate the role played by HDAC4 during senescence, we knocked-out HDAC4 in BJ/*hTERT/Ras/E1A* cells, a standard model of OIS escape (Fig.1g and Fig.S2d/e). HDAC4 KO caused the appearance of SA-β-gal positive cells (Fig.1h) and increased the expression of senescence markers (*CDKN1A, IL1B, IGFBP7*) (Fig.S2f). The induction of senescence in these cells is unrelated to hTERT or E1A levels (Fig. 1g). To confirm the anti-senescence effect of HDAC4, its expression was knocked-out also in low grade leiomyosarcoma cells SK-LMS-1 (Fig.1i and Fig.S2d/e). HDAC4 depletion triggered cell-cycle arrest (Fig.1j) and the appearance of SA-β-gal positive cells (Fig.1k and Fig.S2g). All KO clones failed to grow in semisolid medium, a strong indication of malignancy suppression (Fig.S2h/i). These phenotypes were reproducible in the 4 KO clones generated with 2 different sgRNAs (sg1: 76, 1231; sg2: 205, 1254), while an intermediate phenotype was obtained in the heterozygous clone 275 (Fig. 1j). SK-LMS-1*/HDAC4*^*-/-*^ cells were also characterized by a moderate induction of cell death (Fig.S2j).

*HDAC4* ^*-/-*^ cells were characterized by an altered nuclear morphology (Fig. 1l/m) and the accumulation of γH2AX foci (Fig. 1n). The re-expression of a Cas9 resistant form of HDAC4 (*HDAC4*^*PAM*^) rescued the senescent phenotype in all these clones (Fig. S2k and Fig. 1o), stimulated the entering into the cell-cycle (Fig. 1p), supported their growth in soft agar (Fig. S2l) and reduced the DNA damage levels (Fig. 1q). Similarly, the tamoxifen inducible re-expression of *HDAC4* in SK-LMS-1/*HDAC4*^*-/-*^ cells (clone 66, +4-OHT) (Fig. 1r) rescued every defect observed in the absence of HDAC4 (−4-OHT) (Fig.1r-v, S2m). As further evidence of senescence induction, SK-LMS-1/*HDAC4*^*-/-*^ were sensitive to the senolytic drug navitoclax/ABT*-*263^24^ (Fig. S3a) and lost the linker histone H1^25^ and LMNB1^26^ (Fig. S3b).

The premature senescence onset observed in *HDAC4*^*-/-*^ cells was only partially dependent on MEF2 de-repression. In fact, the expression of a super-repressive mutant^27^ of MEF2 partially restored the proliferation of KO cells (Fig. S3c/d). The appearance of DNA damage and senescence was confirmed in different cellular models (BJ-*TERT/LT*, BJ-*TERT/LT/RAS* and in the melanoma cells WM115) after RNAi-mediated silencing of *HDAC4* (Fig. S3e/f).

### The transcriptome of *HDAC4*^*-/-*^ SK-LMS-1 cells is typical of senescent cells

The transcriptomes of four LMS *HDAC4*^*-/-*^ clones were compared with two control clones (expressing Cas9 or Cas9/sgRNA1 but in which the KO was not achieved) (Fig. S3g). By applying stringent statistical criteria, we identified a minimal signature of 230 genes significantly modulated in all *HDAC4*^*-/-*^ clones (Fig. 2a,S3g and Table S7). 142 out of 230 of these genes (62%) were induced after HDAC4 deletion (Fig. 2b). An unbiased GSEA analysis identified a senescence geneset among the most enriched in *HDAC4*^*-/-*^ cells (Fig. 2c). Moreover, senescence-associated secretory phenotype (SASP), Ras-induced senescence (RIS) and NFKB1 target genes were all positively enriched in *HDAC4*^*-/-*^ (Fig. 2c). Interestingly, in the case of RIS the similarities arose not simply by the SASP but also from the activation of a common epigenetic reprogramming (Fig. S3h).

**Figure 2.**
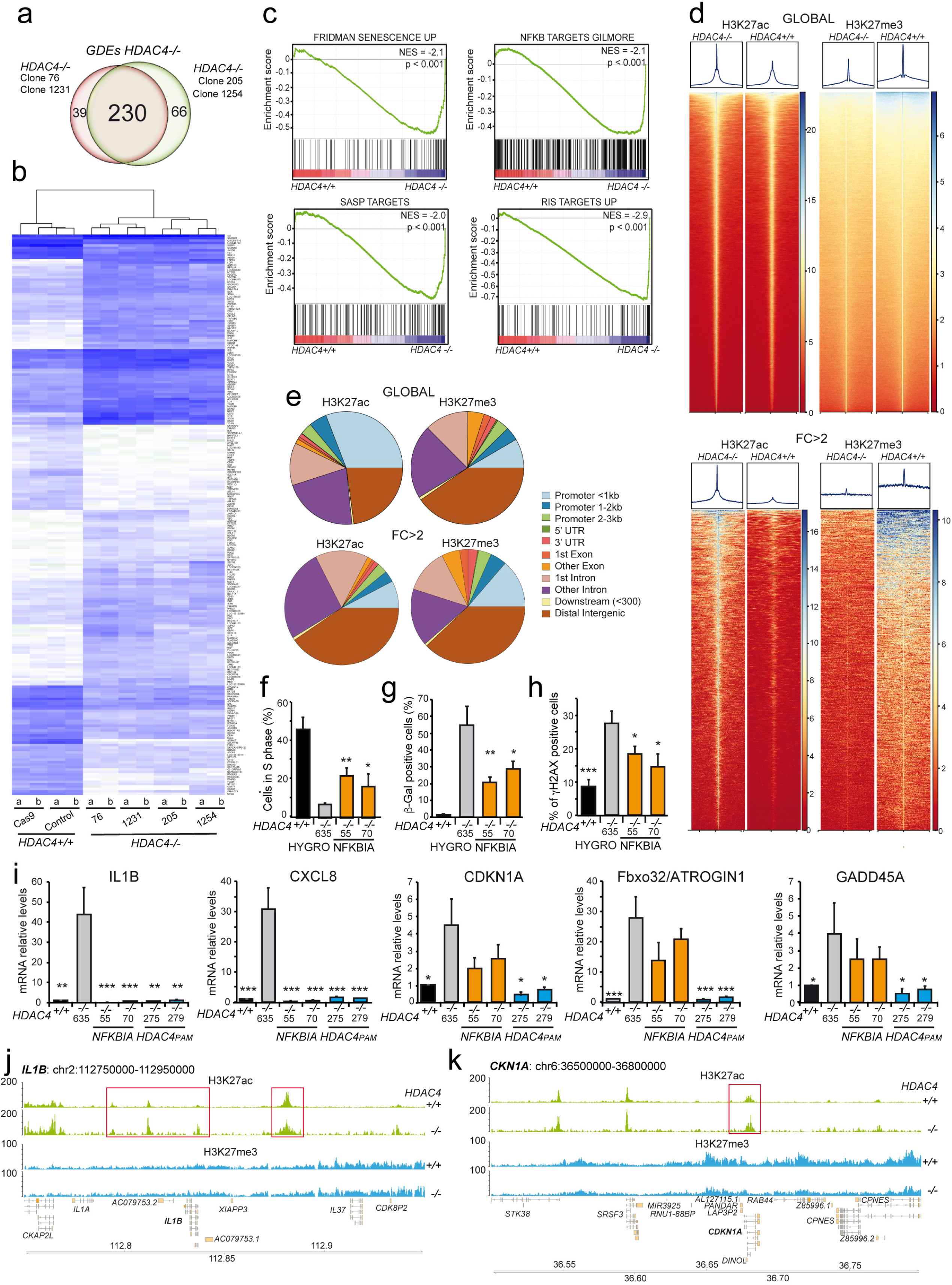
HDAC4 depletion causes a senescence-like transcriptional and epigenetic reprogramming. **a.** Venn diagrams showing the number of transcripts differently regulated in SK-LMS-1/*HDAC4*^*-/-*^ cells generated by using two different sgRNAs (green and red). Differentially expressed genes (DEGs) were selected based on |fold change| >2 and p <0.05. **b.** Heat-map of the absolute expression levels of the DEGs in the indicated clones and biological replicates hierarchically clustered accordingly to average linkage. Blue shades intensity is proportional to transcripts abundance. **c.** GSEA plots displaying the NES obtained by interrogating the transcriptome of HDAC4 wt and KO cells with the indicated gene sets. **d.** (Upper) Heat-map of the 97831 H3K27ac and 139200 H3K27me3 enriched peaks in the indicated SK-LMS-1 cells; (Lower) Heat-map of the 4441 H3K27ac peaks displaying a FPKM (KO/wt)>2 and of the 1205 H3K27me3 peaks displaying a FPKM (wt/KO)>2 (FC>2), in a region of ±15Kb around peak summit. **e.** Genomic distribution of all the H3K27ac and H3K27me3-enriched peaks (top panel), or only of those associated with a FC>2, as indicated and shown in Fig. 2d. **f-g-h.** Analysis of BrdU (f), SA-β-gal (g) and γH2AX (h) positivity in wt or HDAC4 KO cells expressing HYGRO^R^ (clone 635) or GFP-NFKBIA (clones 55 and 70). Mean ± SD; n = 4. The significance is relative to clone 635. **i.** mRNA expression levels of the indicated genes in the indicated SK-LMS-1 clones. Mean ± SD; n = 4. The significance is relative to clone 635. **j-k.** Detailed view of the H3K27ac (green) and H3K27me3 (light-blue) tracks at the *IL1B* (j) and *CDKN1A* (k) loci in SK-LMS-1/*HDAC4*^+*/*+^ and ^*-/-*^, as indicated. Red boxes highlight the H3K27ac regions significantly hyper-acetylated in *HDAC4*^*-/-*^, that correspond to a super-enhancer in the case of *IL1B* (SE_02_17300084) and to an intronic region previously associated to class IIa HDACs binding for *CDKN1A*. In the latter case a strong demethylation in *HDAC4*^*-/-*^ affects the whole locus.

Genome-wide levels of H3K27ac and H3K27me3 showed moderate and focused decreases of H3K27me3 and locus specific increases of H3K27ac, at 36h from HDAC4 depletion (Fig. 2d). Interestingly, the re-acetylation observed in *HDAC4*^*-/-*^ cells involved mainly intronic (18%) and intergenic regions (45%), as similarly observed in other studies^14,28,29^, while the demethylation occurred close to TSS (Fig.2e).

To unveil the contribution of NFKB in this senescent response, we inhibited its activity by introducing IKBα S32A/S36A^30^ (referred hereinafter as NFKBIA) in *HDAC4*^*-/-*^ cells. NFKB1 blockage recovered senescence only partially (Fig. 2f-h). In particular, while the expression of SASP genes (*IL1B, CXCL8*) strongly depends on NFKB1, the anti-proliferative response (*CDKN1A, GADD45A, Fbox32/ATROGIN*) was rescued only by HDAC4 (Fig. 2i and S3i). This was confirmed at chromatin level. 2% and 8% of the H3K27ac regions hyper-acetylated in *HDAC4*^*-/-*^ cells organize the chromatin respectively in: a) typical enhancers (TE) and b) super-enhancers (SE), found activated during senescence (TESs and SESs) (Fig.S3j/k). The latter, exemplified by the *IL1B-IL1A-IL37* SES (Fig.2j), display high preference for NFKB, JUN/AP-1 and MEF2 binding (Table S2). The activation of typical enhancers, as well as the increased acetylation and decreased methylation of H3K27 on gene promoters, can control the expression of negative regulators of cell proliferation, as exemplified by the *CDKN1A* locus (Fig. 2k).

### HDAC4 depletion triggers senescence in melanoma cells

The best-characterized *in vivo* model of OIS is the human nevus^31^. Among the 18 HDACs, only HDAC4 expression was significantly decreased in nevi and significantly increased in melanomas (Fig. S4a). Moreover, high levels of HDAC4 negatively correlated with patients’ survival (Fig. S4b). We therefore knocked-out HDAC4 in A375 melanoma cells. Initially, we obtained only heterozygous clones, suggesting an important role of HDAC4 in regulating cell fitness. Therefore, we generated Dox-inducible cell lines to conditionally express HDAC4^PAM^ before its targeting. With this strategy, five *HDAC4*^*-/-*^ (320, 304, 401, 150, 1090) and two *HDAC4*^+*/-*^ clones (159, 317) were isolated (Fig. S4c). Similarly to sarcoma cells, HDAC4KO caused the downregulation of LMNB1 (Fig. S4c). A time-course analysis in 304^-/-^ cells evidenced that HDAC4 loss triggered the rapid accumulation of DNA damage and the activation of TP53 (Fig. S4d). A pool of genes up-regulated in SK-LMS-1/*HDAC4*^*-/-*^ cells turned out to be up-regulated also in A375 *HDAC4*^*-/-*^ cells (Fig. S4e). Finally, removal of Dox-dependent HDAC4 expression elicited the acquisition of a senescent-like phenotype (Fig. S4f-i). All these phenotypes were weaker in heterozygous clones and the re-expression of HDAC4 rescued the normal phenotype (Fig. S4e-i).

### Loss of *Hdac4* accelerates senescence in non-transformed cells

Primary murine embryonic fibroblasts (MEFs) senescence rapidly when grown at atmospheric oxygen, while the maintenance of hypoxic conditions preserves their proliferation^32^. The conditional (4-OHT dependent) KO of *Hdac4* in MEFs under normoxia sped up the progressive increase in Cdkn2a/p16 levels (Fig. S5a) and the accumulation of DNA damage (Fig. S5b). Similarly, with respect to wt cells, *Hdac4*^*-/-*^ MEF earlier arrested proliferation (Fig. S5c,d) and switched on a senescence signature (Fig. S5e).

We confirmed that MEFs grown in 2% O_2_ did not reach senescence, independently from the absence or presence of HDAC4 (Fig. S5f/g). Curiously, after prolonged time of culture, *Hdac4*^*-/-*^ MEFs evidenced a proliferative crisis from which they rapidly emerged (Fig. S5g).

### Loss of H1.2 and LMNB1 triggers a second wave of DNA damage in HDAC4^-/-^ cells

To investigate the timely order of senescence appearance, we took advantage of the *HDAC4*^*-/-*^ */HDAC4*^*PAM*^*-ER* cells expressing H1.2-GFP as senescence sensor (Fig. S3b). Apparently, the loss of LMNB1 and the accumulation of DNA damage are concomitant events that are coupled to the loss of the linker histone H1 (Fig. 3a). qRT-PCR analysis suggested that the modulation of IGF1 signaling and of the cell-cycle *(CDKN1A)* are early events (Fig. 3b). Immunofluorescence analysis using different markers lead us to define six major phenotypes as exemplified in figure 3c: i) cells positive for H1.2 (H1.2+); ii) cells with a discontinuous (altered) LMNB1 ; iii) cells presenting cytosolic chromatin fragments (CCF^+^); iv) cells with multilobed nuclei; v) cells H1.2 negative (H1.2^-^) with multilobed nuclei and vi) cells with DSBs (γH2AX^+^). This detailed analysis evidenced that DNA damage increased linearly in the first 48h after HDAC4 removal and exponentially in the next 24h, when the loss of H1.2 and LMNB1 became consistent (Fig. 3d).

**Figure 3.**
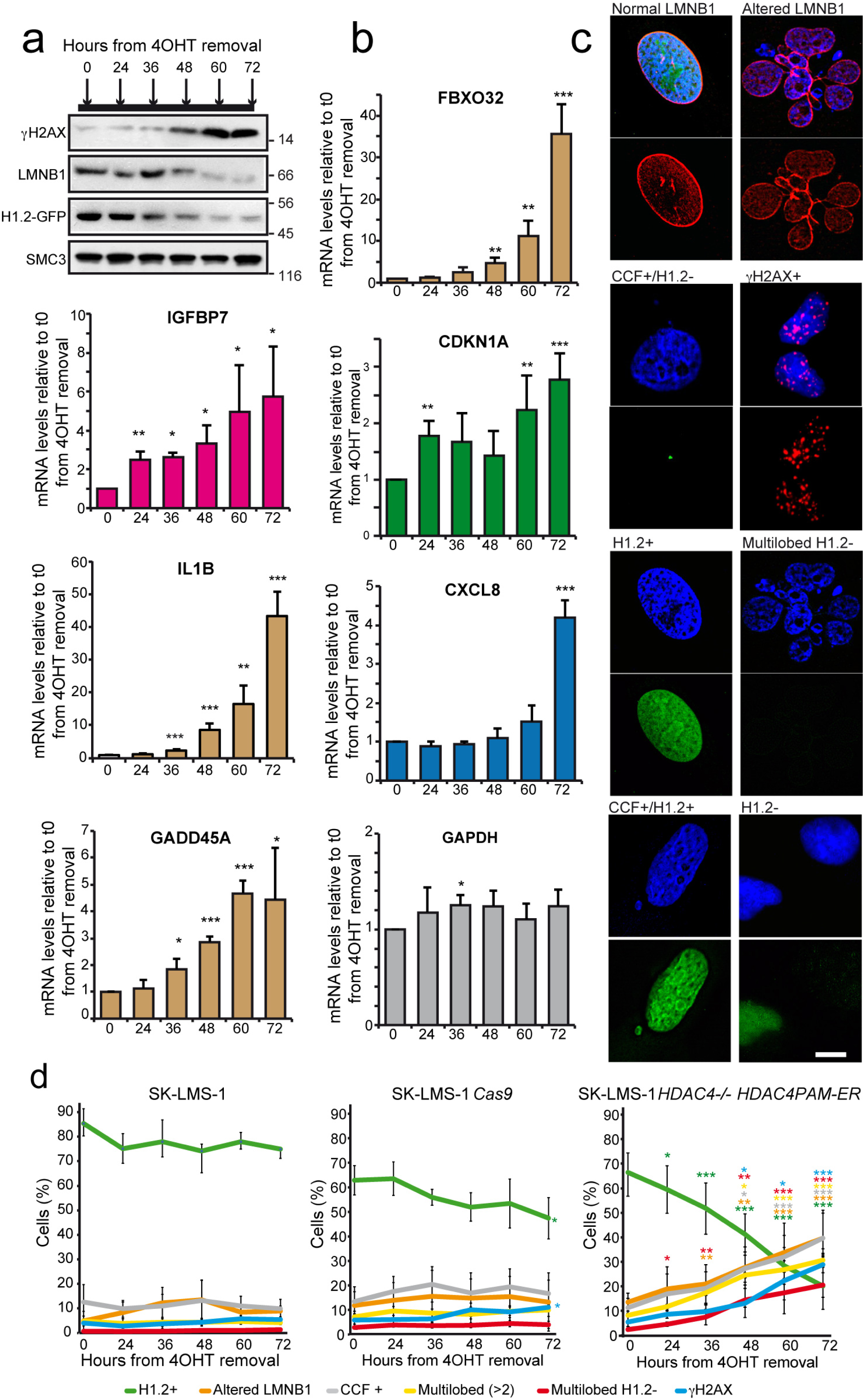
Time-course morphological and transcriptional alteration induced by HDAC4 depletion. **a.** Immunoblot analysis of LMNB1, γH2AX and H1.2-GFP in SK-LMS-1/*HDAC4*^*-/-*^ cells re-expressing *HDAC4*^*PAM*^*-ER*. Cells were harvested at the indicated time after 4-OHT removal. SMC3 was used as loading control. **b.** mRNA expression levels of the indicated genes. Mean ± SD; n = 4. **c.** Representative images of the combination of cellular phenotypes observed after the depletion of HDAC4 in SK-LMS-1 cells (scale bar 10 µm). **d.** Quantification of the time-course accumulation of the phenotypes represented in Fig. 3C in the indicated cells. Mean ± SD; n = 5. At least 200 individual cells were evaluated in each biological replicate.

### *In vivo* definition of the early response to HDAC4 depletion

To gain more insight into the early events triggered by HDAC4 depletion, we introduced in *HDAC4*^*- /-*^*HDAC4*^*PAM*^*-ER* cells two reporters (H2B-GFP and Apple-TP53BP1), to monitor mitosis^33^ and the accumulation of DNA damage^34,35^ *in vivo*, at single cell level. Unperturbed cells accumulate DNA lesions during interphase, possibly during the proceeding through the S phase. Usually these lesions are resolved in G2, before entering mitosis. After mitosis, cells emerge in G1 with few TP53BP1 positive foci defined as nuclear bodies (TP53BP1-NBs) that could represent under-replicated DNA^35–37^.

Cells were monitored over a period of 74 hours from 4-OHT removal/addition by *in vivo* time-lapse confocal microscopy. During this time, 9 out of 10 cells re-expressing HDAC4 entered mitosis twice (Fig. 4). Cell cycle length was constant in these cells (43.22±3.40 hours). The analysis of TP53BP1 dots evidenced the accumulation of DSBs (Supplementary video S1), reflecting the generation of spontaneous DNA or chromatin lesions^36^. Frequently, small TP53BP1 nuclear foci co-existed with larger TP53BP1-NBs, as previously observed^35^. The accumulation of all TP53BP1 dots was quantified and represented as a heatmap (Fig. 4 +4-OHT). The pattern of endogenous TP53BP1 distribution in NBs and in small foci was equal to what observed with the fluorescent sensor (Fig.S6a,b,c).

**Figure 4.**
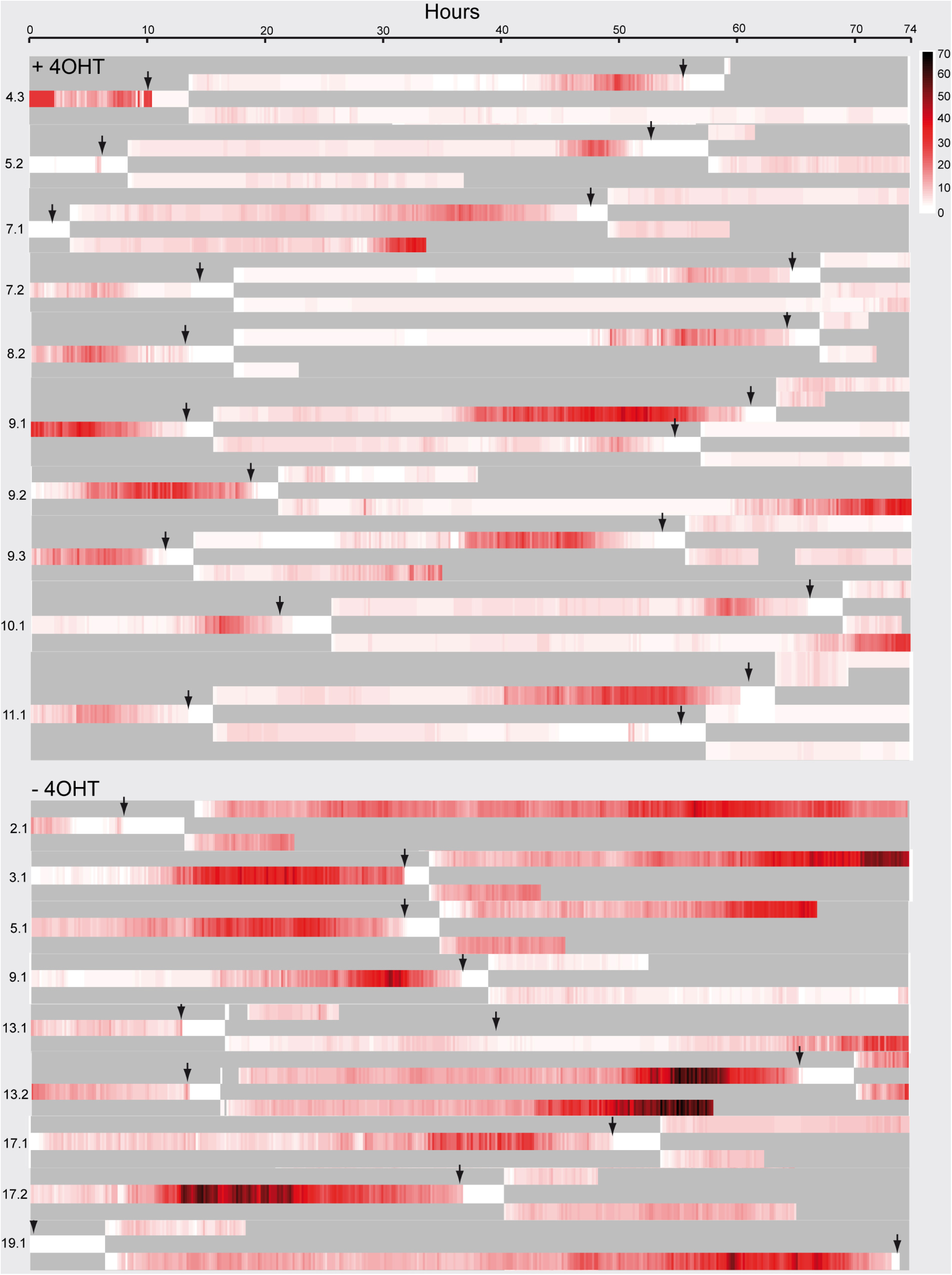
HDAC4 depletion causes the rapid accumulation of TP53BP1 foci and bodies in G2 and the subsequent mitotic slowdown and impairment. Heatmap representing the quantification of the TP53BP1 foci/bodies in SK-LMS-1/*HDAC4*^*-/-*^/*HDAC4*^*PAM*^*-ER*, during 74h of analysis starting from 6h after 4-OHT removal (time “0”), as indicated. The intensity of the red is proportional to the number of TP53BP1 spots. 10 and 9 starting cells were analyzed respectively for the +4-OHT and the -4-OHT conditions.

HDAC4 removal impaired proliferation. In fact the analyzed cells, with two exceptions (13.2 and 19.1), did not enter the second mitosis. Moreover, *HDAC4*^*-/-*^ cells were characterized by the progressive accumulation of TP53BP1 nuclear foci (Fig. 4 -4-OHT and supplementary movie S2) and by a marked increase in the frequency and dimension of TP53BP1-NBs (Fig. S7). These TP53BP1-NBs were not always symmetrically distributed among sister cells after mitosis (supplementary movie S2). In summary, the absence of HDAC4 exacerbates the trend of DNA lesion accumulation, possibly occurring in late S phase and G2.

To correlate the accumulation of pre-mitotic TP53BP1 nuclear dots with mitotic defects, we measured the average number of all TP53BP1 dots for 100 minutes preceding the appearance of chromosomes condensation (mitosis onset). In the presence of HDAC4 the number of TP53BP1 dots was comparable among the analyzed cells (range of 0-3/cell) and the mitosis length was quite homogeneous (100-150 min) (Fig. S6d). A similar behavior was observed during the second mitosis (Fig. S6d). In *HDAC4*^*- /-*^ cells the number of TP53BP1 nuclear dots in G2 was increased, as well as the length of mitosis (Fig. S6d,f).

As a consequence of the unresolved chromosomal lesions that are transmitted to daughter cells, TP53BP1-NBs re-emerged in G1^36^. In the presence of HDAC4, daughter cells emerged with few symmetrical TP53BP1-NBs and this behavior was conserved in the succeeding mitosis (Fig. S6e). In absence of HDAC4 the number of TP53BP1-NBs was increased and they were sometimes asymmetrically distributed between daughter cells. The accumulation of macroscopic nuclear alterations (CCFs, ultrafine DNA bridges (UFBs), lobate nuclei and unproductive mitosis, Fig. S6g) could explain the mitotic delay observed in the absence of HDAC4 (Fig. S6f).

### ROS and LMNB1 play a minor role in genomic instability of *HDAC4*^*-/-*^ cells

Alterations in the nuclear lamina and increased ROS (reactive oxygen species) generation are two possible sources of DNA damage^38,39^, strongly correlated to senescence onset^40^. LMNB1-GFP was overexpressed in SK-LMS-1/*HDAC4*^*-/-*^*/HDAC4*^*PAM*^-ER cells cultured in normoxia or in hypoxia to evaluate their separate or joint contribution to HDAC4-mediated senescence. The overexpression of LMNB1 had a partial effect on the appearance of senescence (Fig. 5a) and, while it did not reduce the total number of DNA CCFs, it reduced the appearance of naked (with a defective LMNB1 envelope) CCFs (Fig. 5b). Interestingly, naked CCFs were frequently positive for γH2AX (arrows and arrowheads Fig. 5b). Accordingly, in LMNB1-GFP cells the increase of γH2AX signal, observed upon HDAC4 depletion, was slightly reduced (Fig. 5d). Growing the same cells in hypoxia did not affect the accumulation of DNA damage and the appearance of senescence (Fig. 5c/d/e). However, hypoxia modestly potentiated the effects of LMNB1 re-expression in escaping senescence (Fig. 5e). As previously reported^41^, LMNB1 re-expression almost completely suppressed SASP, whereas growing cells in hypoxia showed only a modest effect on the expression of *GADD45A* and *CXCL8* (Fig. 5f).

**Figure 5.**
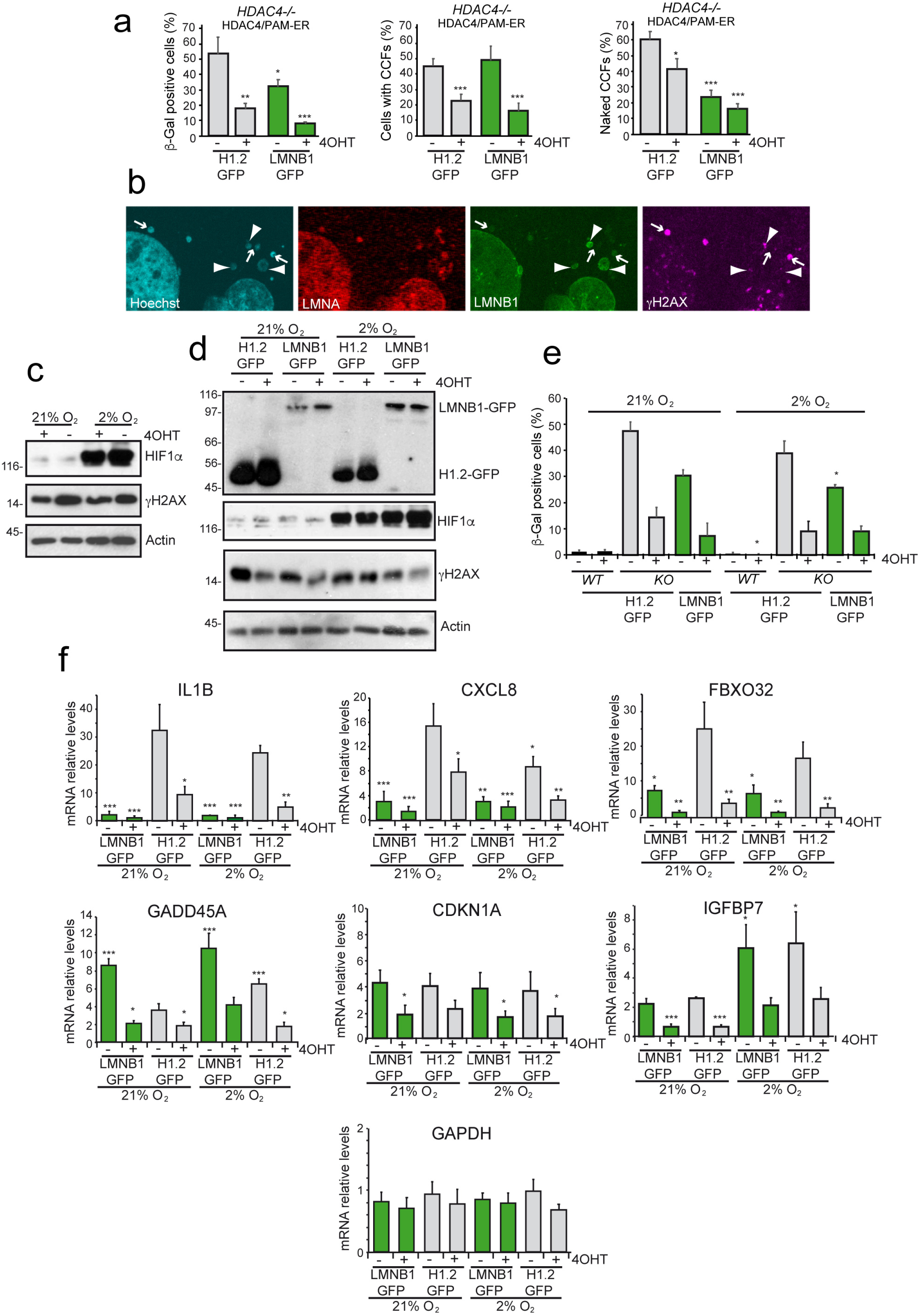
LMNB1 re-expression and hypoxic growing conditions have minimal and complementary effects on the senescence induced by HDAC4 loss in SK-LMS-1 cells. **a.** Analysis of the % of SK-LMS-1/*HDAC4*^*-/-*^ cells, grown in normoxia, expressing H1.2-GFP or GFP-LMNB1, as indicated, and re-expressing (+4-OHT) or not (−4-OHT) *HDAC4*^*PAM*^*-ER*, displaying positivity for SA-β-gal or the accumulation of CCFs and naked CCFs. Mean ± SD; n = 4. The significance is relative to H1.2-GFP re-expressing cells. **b.** Representative confocal picture of SK-LMS-1/*HDAC4*^*-/-*^ cells, expressing GFP-LMNB1 and immunostained for DNA (Hoechst, blue), LMNA (red) and γH2AX (violet). Arrows point to naked CCFs, arrowheads to LMNB1+ CCFs. **c-d.** Immunoblot analysis of HIF-1α, γH2AX, GFP-LMNB1 and H1.2-GFP (anti-GFP antibody) in SK-LMS-1/*HDAC4*^*-/-*^ cells, re-expressing (+4-OHT) or not (−4-OHT) *HDAC4*^*PAM*^*-ER* and expressing the indicated transgenes. Lysates were generated after 4d of culture in normoxia or in hypoxia, as indicated. **e.** Analysis of the SA-β-gal positivity in the cells described in Fig. 5D. The significance is relative to the same cells grown in normoxia. Mean ± SD; n = 3. **f.** mRNA expression levels of the indicated genes in SK-LMS-1 cells generated and maintained as described in Fig. 5D. The levels are relative to wt cells grown in normoxia (considered as 1). The significance is relative to SK-LMS-1/*HDAC4*^*-/-*^ cells grown in normoxia. Mean ± SD; n = 3.

### DNA lesions occurring during replication accumulate in the absence of HDAC4

DNA lesions frequently arose during DNA replication. Several intracellular and extracellular conditions can trigger RS and forks stalling or collapsing that lead to DSBs^42^. Damaged replication forks are marked by ATR-mediated hyperphosphorylation of ssDNA-bound RPA32 and by the monoubiquitylation of PCNA. These PTMs are instrumental for the recruitment of repair factors and Polη^43,44^. Starting from 12h after the switch-off of *HDAC4*^*PAM*^*-ER* expression, *HDAC4*^*-/-*^ cells accumulated Ub-PCNA and phosphorylated (S4/S8) RPA32, which became more evident after 48 hours. These accumulations were paired to the induction of DSBs (Fig. 6a). Most of this DNA damage is due to RS. In fact, inhibition of cell cycle progression, achieved through the CDK4 inhibitor Palbociclib, reduced PCNA ubiquitylation, almost abolished RPA32 phosphorylation and reduced the increase in DSBs (Fig 6b), similarly to HDAC4 re-expression (Fig. 6c). Low doses of aphidicolin (APH) or camptothecin (CPT) can trigger RS^45,46^. Depletion of HDAC4 showed additive effects to APH or CPT for DSBs accumulation (Fig. 6d-e). Similarly, the recovery after the APH block was delayed by the absence of HDAC4 (Fig. 6f). Overall, these experiments indicate RS as an important source of DNA damage in *HDAC4*^*-/ -*^ cells.

**Figure 6.**
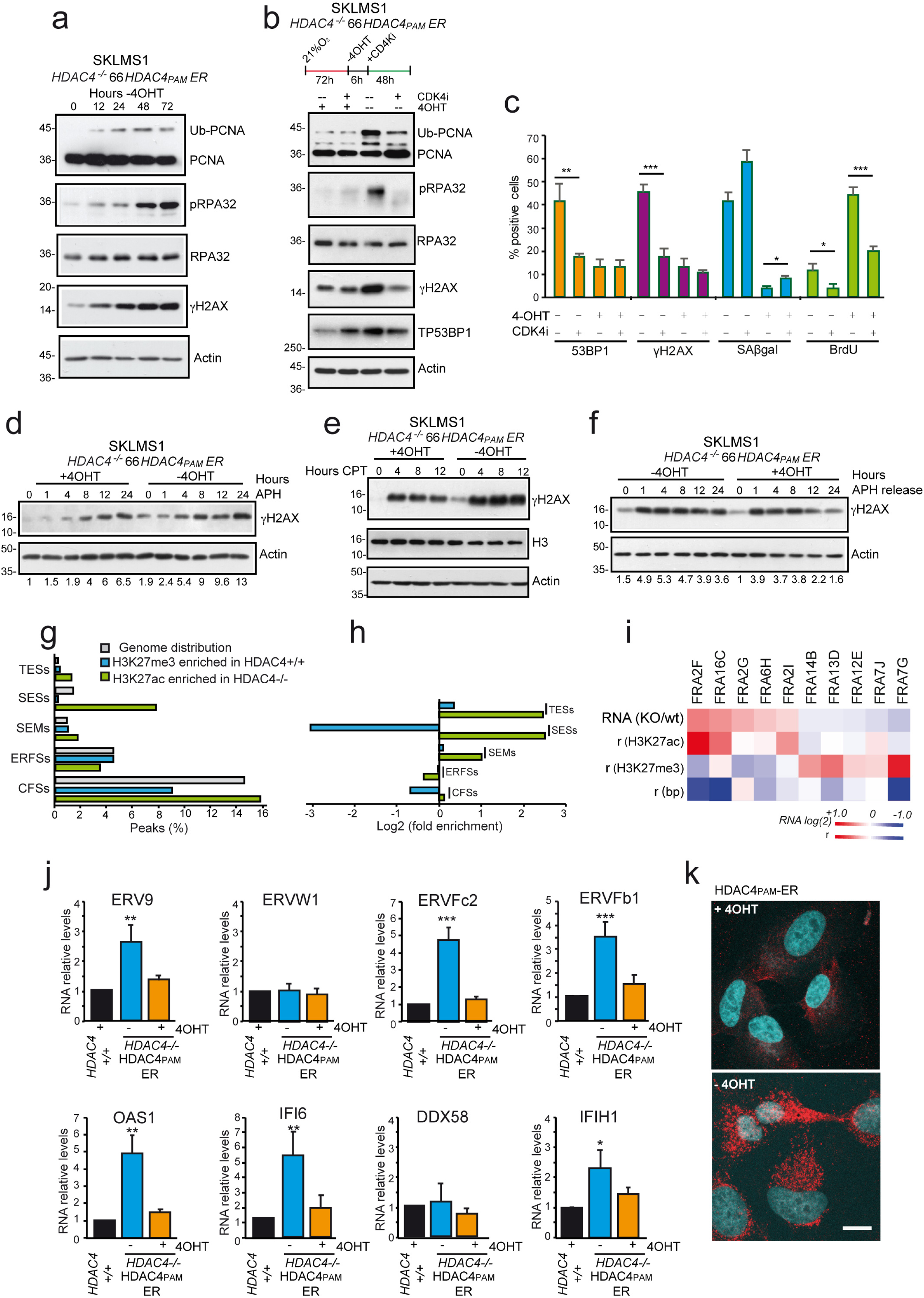
HDAC4 depletion causes the accumulation of replication stress and of cytoplasmic dsRNAs of retroviral origin. **a.** Immunoblot analysis in SK-LMS-1/*HDAC4*^*-/-*^ cells re-expressing *HDAC4*^*PAM*^*-ER*. Cells were harvested at the indicated time after 4-OHT wash out. Actin was used as loading control. **b.** Immunoblot analysis in SK-LMS-1/*HDAC4*^*-/-*^ cells re-expressing *HDAC4*^*PAM*^*-ER*. 4-OHT was removed 6h before the treatment with CDK4i (1µM), as indicated. Actin was used as loading control. **c.** Analysis of BrdU, SA-β-gal, TP53BP1 foci and γH2AX positivity in the indicated cells treated as in Fig. 6B. Mean ± SD; n = 4. The significance is relative to untreated cells. **d-e.** Time-course immunoblot analysis in SK-LMS-1/*HDAC4*^*-/-*^ cells re-expressing or not *HDAC4*^*PAM*^*-ER* (48h) and treated for the indicated time with 500nM Aphidicolin (APH) or 3.125µm Camptothecin (CPT). Actin was used as loading control. Densitometric analysis of γH2AX/Actin ratio is provided. **f.** Immunoblot analysis on the same cells described in Fig. 6D, harvested at the indicated time after the release from 1h APH treatment. Densitometric analysis of γH2AX/Actin ratio is provided. **g.** Histogram representing the percentage of hyper-acetylated (green bar) or demethylated (light blue) H3K27 peaks in SK-LMS-1/*HDAC4*^*-/-*^ cells and falling in the indicated genomic elements or displaying the indicated epigenomic features. The genome coverage of each element is indicated by gray bars. **h.** Histogram representing the enrichment of each element described in Fig. 6g in respect to the expected distribution calculated according to the genome coverage. **i.** Histogram of 10 CFSs associated to the transcripts (median of the associated transcripts, indicated as RNA (KO/wt)) more up-regulated (red) or down-regulated (blue) in SK-LMS-1/*HDAC4*^*-/-*^ in respect to *HDAC4*^+*/*+^ cells. The correlation between the RNA (KO/wt) and H3K27ac/me3 levels and the gene length are indicated. **j.** Histogram representing the RNA levels of the indicated genes/ERVs in SK-LMS-1*/HDAC4*^*-/-*^/*HDAC4*^*PAM*^*-ER* cells, at 72h from last 4-OHT treatment (+4-OHT) or wash-out (−4-OHT). Mean ± SD; n = 4. The significance is relative to wt cells. **k.** Representative confocal pictures of the indicated SK-LMS-1 cells at 48h from 4-OHT removal/addition. Immunofluorescence was performed to visualize dsRNAs (red) and nuclei (blue). Scale bar 10µM.

*HDAC4*^*-/-*^ cells did not show evident marks of epigenetic stress that could lead to transcription/replication conflicts^47–49^ in correspondence of genomic common fragile sites (CFSs)^50^ or early replicating fragile sites (ERFSs)^51^. Indeed, some epigenetic alterations fall in CFSs (5.5% with altered H3K27ac and 7.4% with altered H3K27me3) (Fig.6g/h) and, even if they correlate well with the expression of the associated genes (ρac(H3K27ac KO/wt)=0.37, ρme(H3K27me3 KO/wt)= -0.32) (Fig.6i), only 16% of HDAC4-regulated genes (13% up-regulated, 20% down-regulated) fall in CFSs, that cover 15% of the genome. In conclusion, the source of RS that characterizes *HDAC4*^*-/-*^ cells cannot be ascribed to abrupt epigenetic and transcriptional changes at ERFSs and CFSs.

### HDAC4 depletion causes the activation of ERV transcripts

Epigenetic perturbations elicit the transcription of ERVs^52,53^ during cellular senescence, aging and cancer^54,55^. Demethylating agents and HDACi trigger the expression of ERVs^52,54^. All tested ERVs, with the exception of *ERVW1* that contains a coding gene (*SYCY1*), were induced in *HDAC4*^*-/-*^ cells (Fig. 6j). Similarly, the IFN response was switched on (Fig. 6j), as a consequence of dsRNAs accumulations (Fig. 6k). The time-course analysis confirmed the up-regulation of ERVs and of the IFN response. The levels of *ERV9* and *ERVFc2* peaked at 60h from HDAC4 removal and then decreased, probably because of an OAS1-dependent degradation (Fig. 7a). The same pattern of ERVs was upregulated during replicative senescence in human fibroblasts (Fig. 7b). ERVs and the IFN response were induced in IMR90-*E1A/RAS* transformed cells after HDAC4 KO (Fig. S8a), in dermis and liver of aged mice (Fig. S8b/c), in *HDAC4*^*-/-*^ A375 cells (Fig. S8e). ERVs up-regulation was not due to Cas9 activity or nuclear lamina dismantling, as they were unperturbed by the KO of unrelated genes (Fig. S8d) or by the re-expression of LMNB1 in *HDAC4*^*-/-*^ SK-LMS-1 cells (Fig. S8f). Interestingly, in MCF10A cells the KO of HDAC7, the most abundant class IIa HDAC, induced ERVs expression and this up-regulation was increased in the presence of RAS (Fig. S8g).

**Figure 7.**
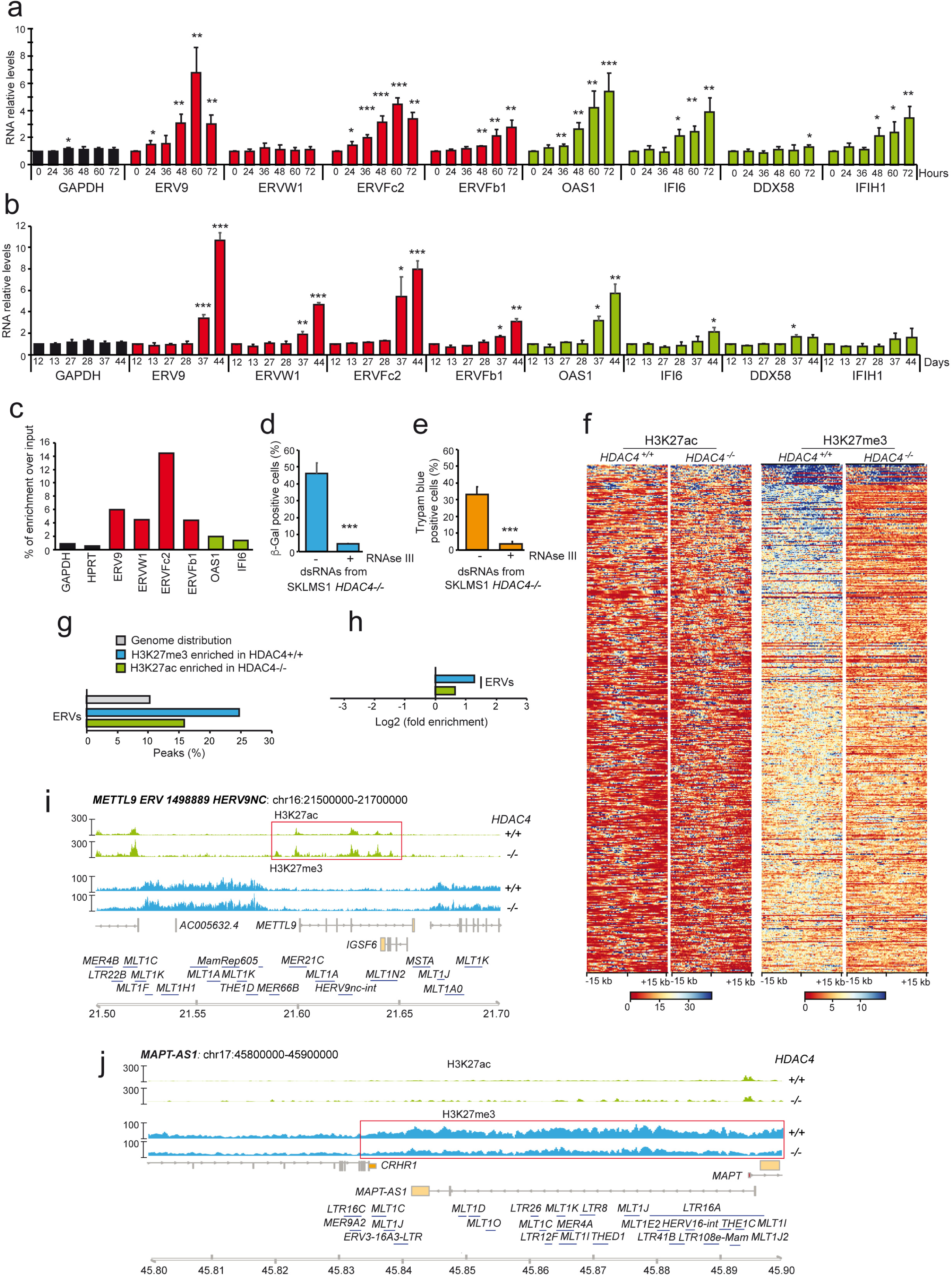
The epigenetic stress induced by HDAC4 depletion causes the accumulation of ERVs and triggers the IFN-response. **a-b.** mRNA expression levels of the indicated genes, in SK-LMS-1/*HDAC4*^*-/-*^ */HDAC4*^*PAM*^*-ER* re-expressing cells harvested at the indicated time after HDAC4 depletion (a) and in IMR90 cells harvested at the indicated cellular splittings (b). Mean ± SD; n = 3. The significance is relative to time 0 (a) or to split 12 (b). **c.** Quantification of the enrichment over input of the indicated species of RNAs in the preparation of dsRNAs used to treat the cells in Fig. 7d/e. **d-e.** Analysis of SA-β-gal (c) and Trypan blue (d) positivity in SK-LMS-1 cells transfected for 72h with 30pmoles of double-stranded enriched RNAs obtained from *HDAC4*^*-/-*^ cells, and pre-digested or not for 30’ with 50u RNAseIII. **f.** Heat-map of the top (1%) H3K27ac and H3K27me3 enriched peaks in the indicated SK-LMS-1 cells in a 30kb region around an ERV; each row represents the same ERV in wt and KO cells. **g.** Histogram representing the percentage of hyper-acetylated (green bar) or demethylated (light blue) H3K27 peaks in SK-LMS-1/*HDAC4*^*-/-*^ cells and falling in ERVs. The genome coverage of each element is indicated by gray bars. **h.** Histogram representing the enrichment of hyper-acetylated or demethylated ERVs as explained in Fig. 7g in respect to the expected distribution calculated according to the genome coverage. **i-j.** Detailed view of H3K27ac (green) and H3K27me3 (light-blue) tracks at two ERVs rich regions on Chr16 (g) and Chr17 (hH). Regions H3K27 hyper-acetylated (g) or demethylated (h) in SK-LMS-1/*HDAC4*^*-/-*^ in respect to the wt are indicated.

Importantly, the exogenous delivery of dsRNAs species immunopurified from *HDAC4*^*-/-*^ cells triggered senescence and cell death in SK-LMS-1 cells (Fig. 7c-e). In *HDAC4*^*-/-*^ cells, ERVs are characterized by a depletion of H3K27me3 and by an increase in the H3K27ac/H3K27me3 ratio (Fig. 7f). The strong demethylation of K27 of H3 wrapping the genomic DNA of ERVs represents the second most pronounced epigenetic alteration related to HDAC4 removal after the activation of TESs and SESs (Fig. 7g-j).

### A MAVS/ERV dependent pathway sustains HDAC4-dependent senescence

To better define the contribution of ERVs and of the DDR in this senescence response, we suppressed these pathways by taking advantage of dominant negative (DN) versions of MAVS (MAVSΔCARD)^56^ and TP53 (TP53^R175H^)^57^. MAVS was selected since MDA5-RIG1-MAVS recognizes the dsRNA shape of ERV transcripts and triggers the activation of a NFKB/IFN/TNF antiviral responses, which sustain senescence^58,59^.

The expression of TP53-DN and of MAVS-DN in *HDAC4*^*-/-*^ cells reduced SA-β-gal positivity more efficiently than MEF2-DN (Fig. 8a,h). The impact on counteracting the DDR showed a gradient of efficiency with HDAC4>MAVS-DN>TP53-DN>MEF2-DN (Fig. 8b). The effect of MAVS-DN on DNA damage was unexpected. Hence, we evaluated this suppressive effect in relation to the RS using the CDK4i. Figure 8c shows that MAVS-DN did not completely suppress DNA damage and a stronger effect was observed with the co-expression of HDAC4. As expected, the level of DNA damage was further dropped after the block of the cell cycle operated by CDK4i. This result demonstrates that dsRNA/MAVS are secondary sources of RS that add to the primary and earlier generation, triggered by HDAC4 depletion. The senescence rescue of TP53-DN and MAVS-DN was confirmed by LMNB1 expression (Fig. 8d/e). The role of MAVS in this senescent response was verified by silencing its expression and that of RIG1, using RNAi (Fig. 8f). Next, we evaluated the influences on a pattern of genes (ERVs, SASP, IFNs and cell cycle genes) defining the senescent signature (Fig.8g/h). Re-expression of HDAC4 efficiently rescued all observed alterations. MAVS-DN impacted on SASP and IFN genes, but only partially on cell cycle genes. TP53-DN was ineffective in suppressing ERVs and some SASP genes but effective for the IFN response. LMNB1, NFKBIA and MEF2-DN partially impacted on the senescent signature and mainly at the level of SASP expression (Fig. 8g/h).

**Figure 8.**
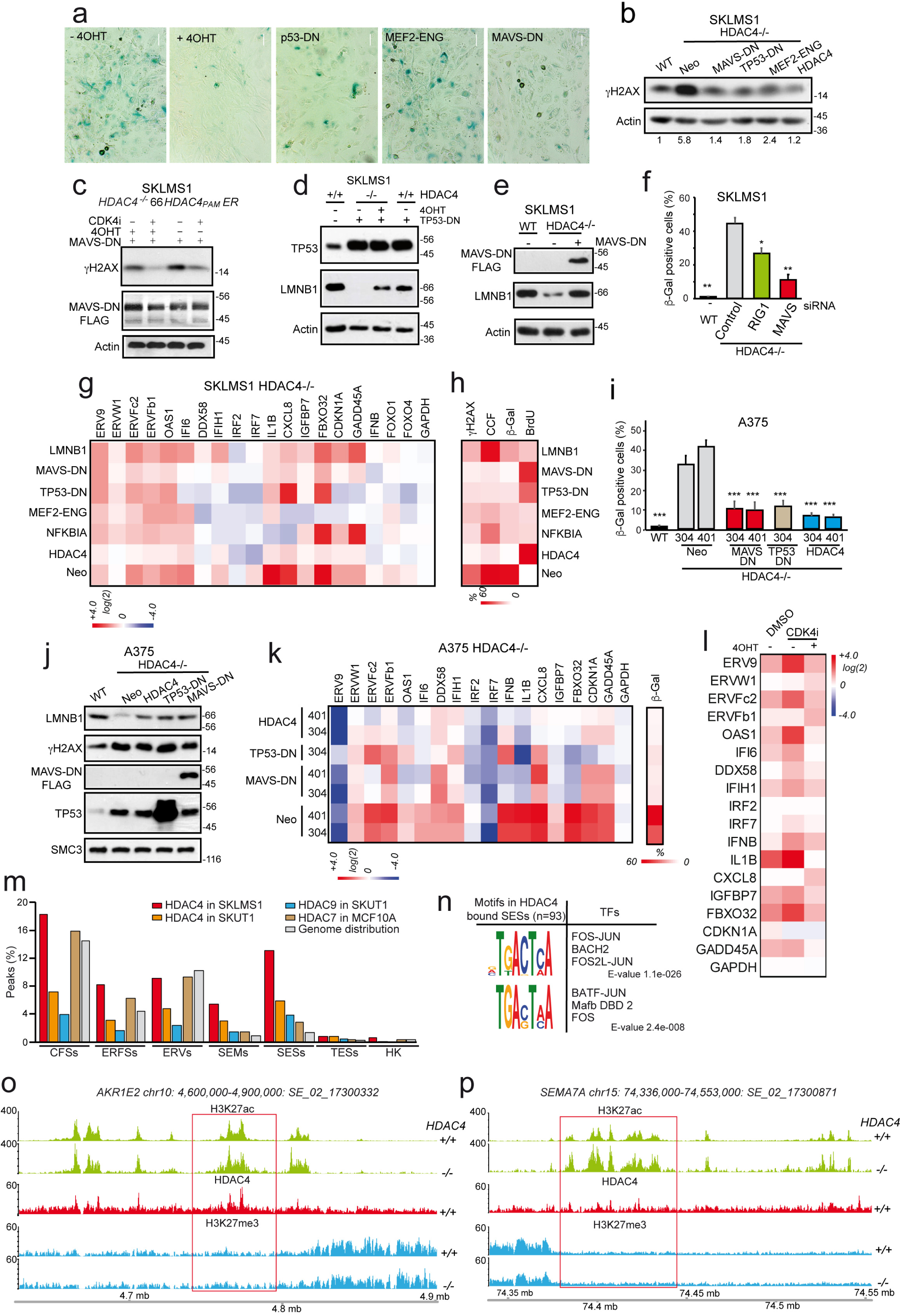
A senescence signature triggered by the activation of RIG-1/MAVS pathway or by the HDAC4-dependent de-repression and activation of SE sustains the premature senescence entrance. **a.** Microscopic images of SA-β-gal stained SK-LMS-1/*HDAC4*^*-/-*^ cells re-expressing HDAC4 (+4-OHT) or the indicated transgenes, at 72h from HDAC4 depletion. Scale bar 50 µm. **b.** Immunoblot analysis in the cells described in Fig.8a. Densitometric analysis of γH2AX/Actin ratio is provided. n=3. **c.** Immunoblot analysis in the indicated cells treated as in fig. 6b. **d-e.** Immunoblot analysis in the indicated cells harvested after 72h of HDAC4 depletion. Actin was the loading control. **f.** Analysis of SA-β positivity in wt or *HDAC4*^*-/-*^ SK-LMS-1 cells transfected with the indicated siRNAs. Mean ± SD; n =3. **g.** Heat-map reporting the expression levels (log2 fold change relative to wt cells) of the indicated genes and ERVs in SK-LMS-1/*HDAC4*^*-/-*^ cells expressing the indicated transgenes as described in Fig. 8b. **h.** Heat-map reporting the % of the indicated SK-LMS-1 cells displaying positivity to γH2AX, CCF, SA-β-gal and BrdU. **i.** Analysis of SA-β-gal positivity in wt or *HDAC4*^*-/-*^ A375 cells re-expressing the indicated transgenes, 5d after HDAC4 removal. Mean ± SD; n =3. The significance is relative to *HDAC4*^*-/-*^-Neo cells. **j.** Immunoblot analysis in the indicated cells harvested as in Fig. 8i. SMC3 was the loading control. **k.** Heat-map reporting the expression levels (log2 fold change relative to wt cells) of the indicated genes and ERVs (left) and SA-β-gal positivity (right) in the same A375 cells described in fig. 8i. **l.** Heat-map reporting the expression levels (log2 fold change relative to DMSO treated *HDAC4*^*PAM*^*-ER* cells) of the indicated genes and ERVs, in the same SK-LMS-1 cells described in fig. 6b. **m.** Histogram representing the percentage of class IIa HDACs bound peaks in the indicated cells and falling in the described genomic/epigenomic elements. The genome coverage is indicated by gray bars. **n.** Motif analysis of 93 HDAC4-gained SESs for putative transcription factor binding sites. Motifs with p-value<0.5*10^−4^ were selected. **o-p.** Detailed view of 2 representative SESs directly bound by HDAC4 in SK-LMS-1 wt cells in correspondence to H3K27ac-defined SEs.

We confirmed these observations in melanoma A375 cells. MAVS-DN and TP53-DN (more strongly than in SK-LMS-1) inhibited the HDAC4-dependent senescence, rescued LMNB1 levels (Fig. 8i/j) and modulated the senescent signature (Fig. 8k) similarly to what observed in SK-LMS-1 cells.

### HDAC4 binds and control the acetylation status of super-enhancers

Removal of HDAC4 triggers several dysfunctions including the induction of RS, SEs and the activation of ERVs, which ultimately lead to senescence. Importantly, HDAC4 was still able to repress the senescent signature when CD4Ki was used to induce senescence and cell-cycle arrest with only minimal RS (Fig. 8l). We therefore hypothesized that HDAC4 could act as a safeguard of cell fate by repressing senescent genes. HDAC4 ChIP-seq profile evidenced that HDAC4 binding was highly enriched at the level of specific SE, mainly those activated in senescence (SES) (Fig. 8m). Even though this property is partially shared with the other class IIa HDACs, it particularly marks HDAC4 (Fig. 8m). By binding these SE, possibly through the engagement of transcription factors previously associated with SE like BACH2^60^ and AP-1^3^ (Fig.8n) and/or MEF2^61^ (Table S3), HDAC4 could regulate the expression of BRD4-dependent inflammatory genes (8o,p, Table S4; ρac(H3K27ac KO/wt)=0.35 p=0.02 in BRD4 associated^3^ SESs, ρac=0.04 p=0.86 in SESs not associated to BRD4). This response is reinforced and integrated by the H3K27 demethylation and subsequent de-repression of ERVs, which engages about 10% of HDAC4 peaks (Fig. 8m). In conclusion, HDAC4 has a dual role in regulating senescence, by controlling the replication stress and by repressing directly (SE) or indirectly (through ERV) the inflammatory response.

## DISCUSSION

We have identified HDAC4 as the class IIa HDAC member downregulated in all the tested models of senescence and aging. GSK3β^23^ supervises the UPS-mediated degradation of HDAC4 during senescence. HDAC4 depletion forces the senescence entrance in transformed and pre-transformed cells. These responses stem from the activation of two early events: DNA damage and inflammation. *HDAC4*^*-/-*^ cells become more susceptible to RS. Whether this is due to increased replication fork stalling and collapsing or decreased fork restarting^62^ is presently unknown and deserves further studies. HDAC4-regulated genes falling in CFSs and ERFs underwent normal epigenetic control. This makes it unlikely that a) replication/transcription conflicts^47^ arising in these sites or b) other influences of RNA Pol-II on replication origin firing^48,49^ could be at the origin of HDAC4-dependent RS. Interestingly, class I HDACs were found to bind nascent DNA and protect from RS without altering the acetylation of the histones proximal to the fork^63,64^.

HDAC4 depletion elicits H3K27 hyper-acetylation and activation of TE and SE^3,7^ and the H3K27 de-methylation and de-repression of repetitive elements of retroviral origin (ERVs)^52,55^. The activation of these two complementary epigenetic mechanisms confirms that class IIa HDACs are pivotal nodes for the formation of high molecular weight complexes with deacetylase and methyltransferase activities^65,66^. One third of HDAC4 peaks supervises directly the chromatin remodeling of these elements and promotes the RS-independent activation of a pro-inflammatory signaling. Licensing of SE is emerging as a key aspect of the senescence response^3,7,60,67^. The activation of this pathway could increase the genomic fragility due to HDAC4 depletion, as demonstrated in other studies^68,69^. The dismantling of the nuclear lamina and the loss of the linker H1 can further increase transcriptional dysregulation and DNA damage.

The complex circuit of events that orbit around HDAC4 demonstrate the centrality of the role played by epigenetic regulators in preserving the correct expression and the integrity of the genome.

## ACKNOWLEDGEMENTS

We thank Francesca D’Este for the support with the confocal microscope, Danilo Licastro for the support with ChIP-seq experiments and Matteo Corsano for the support with RS experiments. HDAC4 conditional mutant mice were generously provided by Prof. Eric N Olson and Rhonda Bassel-Duby. This study was supported by PRIN 2017 JL8SRX “Class IIa HDACs as therapeutic targets in human diseases: new roles and new selective inhibitors”, Interreg Italia-Osterreich rITAT1054 EPIC and the Sarcoma Foundation of America (SFA) to C.B.

## AUTHOR CONTRIBUTIONS

CB conceived the project. EDG and CB designed and analyzed all experiments. EDG generated all knocked-out cell lines, prepared the libraries for ChIP-seq and performed all gene expression studies. HP completed RNAi experiments and supported EDG in immunoblot experiments. EDG, ED and RF performed bioinformatics analysis. LR performed some western blotting. CB did the confocal microscopy and time-lapse analysis. EDG, HP and CB analyzed the time-lapse experiments. EDG, VC, HP, AR, VM generated cell lines used in the study. EDG, ED and CB interpreted data. CB designed the figures with the help of EDG, ED and RP. EDG and CB wrote the paper.

## METHODS

### Cell culture and reagents

BJ/hTERT and IMR90 cells (previously characterized^57^), SK-LMS-1 (*TP53*^wt/G245S^) and SK-UT-1 leiomyosarcoma cells (previously characterized^14^), HEK293T, LinXE, Ampho Phoenix and MEF *HDAC4*^*loxp/loxp*^ cells were cultured as previously described^14^ in 10% FBS DMEM (Euroclone). A375^70^ *(TP53*^*wt/wt*^) and WM115^71^ *(TP53*^*wt/wt*^) melanoma cells were grown in RPMI, MCF10A were grown as previously described^72^. MCF10A/*HDAC7*^*-/-*^, U87MG/*GSK3β*^*-/-*^, SK-UT-1/*HDAC4*^*-/-*^ and *HDAC9*^*-/-*^ were previously described^14,28,73^. For the conditioning of BJ/hTERT cells, the medium obtained from BJ/*hTERT/HRAS*^*G12V*^ or *HYGRO* cells cultured in 60mm plates for 8 days, was filtered and diluted 1:1 with fresh medium and used to treat cells twice for 96h, with a change after 48h. For the experiments performed in Hypoxia, cells were grown in hypoxic chambers at 37°C, 5% CO_2_, 2% O_2_ (Baker Ruskinn). The following chemicals were used: 250nM 4-OHT (Sigma-Aldrich), 1µM Doxycycline (Sigma-Aldrich), 1µM MG132 (Sigma-Aldrich), 10µM Chloroquine (Sigma-Aldrich), 0.4% Trypan Blue (Sigma-Aldrich), 200µM H_2_O_2_ (Sigma-Aldrich), 1µM PD0332991 (Sigma-Aldrich), 500nM Aphidicolin (Sigma-Aldrich), 3.125µm Camptothecin (Enzo Life Sciences), 20µM Etoposide (Enzo Life Sciences), 100nM ABT-263 (Clinisciences).

### Generation and culture of *Hdac4*^*fl/fl*^ and *Hdac4* ^*-/-*^ murine embryonic fibroblasts

*Hdac4*^*fl/fl*^ mice were previously described^74^. MEFs were generated following standard procedures^75^ from 13.5 days-old embryos. Single cell suspensions were expanded in DMEM/10% FBS in hypoxia. 5*10^6^ cells were retrovirally infected to express Cre-ER. Not infected cells were removed from culture by puromycin (2µg/ml, Sigma-Aldrich) selection. The recombination was achieved through the treatment for 48h with 4-OHT (Sigma-Aldrich). At the end of this incubation, half of the culture was kept in normoxia and half in hypoxia. At each splitting the total number of cells was counted (Countess II, LifeTechnologies) and the doubling time (dt) was calculated as previously described^76^.

### Plasmid construction, transfection, retroviral and lentiviral infection, silencing

pLENTI-*CRISPR/V2* (Plasmid #52961), p*SpCas9*(BB)-2A-*GFP*(PX458) (Plasmid #48138), p*SpCas9*(BB)-2A-*Puro* (PX459) (Plasmid #62988), pCW-*Cas9* (Plasmid #50661), pCW57/Hygro-MCS1-2A-MCS2 (Plasmid #80922), *Apple-53BP1trunc* (Plasmid #69531), *pEGFP-N1*/H2B-GFP (Plasmid #11680), pBabe-*Puro-IKBalpha* (NFKBIA) S32A/S36A (Plasmid #15291), MSCV-*CreERT2-Puro* (Plasmid #22776) were obtained from Addgene. pLKO-Puro *shHDAC4* 1 (TRCN0000314667) and 2 (TRCN000004832) were obtained from Sigma-Aldrich. pWZL-*Hygro-HDAC4*^*PAM*^ (V31L or P16A), pWZL-*Hygro-HDAC4*^*PAM*^*-ER* pCW-*Hygro-HDAC4*^*PAM*^ were obtained by sub-cloning a mutagenized HDAC4 (QuikChange Site-Directed Mutagenesis Kit, Agilent) into the linearized empty backbones by a restriction-based approach. *Apple-53BP1trunc* and *H2B-GFP* were sub-cloned respectively in pBABE-*Zeo* and pWZL-*Neo*, NFKBIA-S32A/S36A into pWZL-*Neo-GFP*. pWZL-*Neo-MCS1-2A-MCS2* was obtained by sub-cloning the MCS of pCW57 *Hygro MCS1-2A-MCS2* into pWZL-*Neo* through a recombination-based approach. The generated plasmid was used as acceptor vector for the cloning of *HRAS*^*G12V*^ (*Nhe*I/*Sal*I) and *E1A*/1-143 (*Mlu*I/*Bgl*II-*Bam*HI) to generate pWZL-*Neo-HRAS/G12V-*2A*-E1A 1-143*. pWZL-*Neo/GFP-LMNB1*, pWZL-*Neo-MAVSDN*(ΔCARD 1-100), pBABE-Z*eo-MAVSDN* and pWZL-*Neo/H1.2-GFP* were obtained by amplifying the relative cDNAs from IMR90 cells. pWZL-*Hygro-MEF2/ENG*, pWZL-*Hygro-HRAS*^*G12V*^, pBABE-*Puro-HRAS*^*G12V*^ and pBABE-*Puro-myrAKT1* were previously described^57^. pCW-Puro-*HRAS*^*G12V*^ and *myrAKT1* were obtained by sucloning. pLKO-*Hygro* plasmid expressing the same shRNAs were obtained by oligo cloning. All the generated plasmids were checked by restriction and sequencing. The primers used for cloning are listed in Table S5. Transfections, viral infections and siRNA delivery were done as previously described^28,77^. The following siRNAs (148 pmol) were used: HDAC4 (CCACCGGAAUCUGAACCACUGCAUU, Invitrogen Stealth), MAVS 1 (CCACCUUGAUGCCUGUGAA), MAVS 2 (CAGAGGAGAAUGAGUAUAA), RIG-1 (AAUUCAUCAGAGAUAGUCA).

### CRISPR*/*Cas9 Genome editing

SpCas9 was stably transduced to generate SK-LMS1 *HDAC4*^*-/-*^ clones (76,1231,205,1254,66), *HDAC4*^+*/-*^ clone 275 and BJ/E1A-RAS *HDAC4*^*-/-*^. SpCas9 was transiently transfected (Lipofectamine 2000, LifeTechnologies) to generate SK-LMS-1 *HDAC4*^*-/-*^ clones (635, 273, 274, 275, 277, 279), A375 *HDAC4*^*-/-*^ clones (304, 401, 150, 1090) and *HDAC4*^+*/-*^ clones (317, 159). A Cas9 resistant HDAC4 (PAM mutated, using a strategy previously described^28^), was stably expressed prior to the KO (in SK-LMS-1 clones 273,274,275, 277, 279) or continuously re-expressed in a 4-OHT dependent (SK-LMS-1 clone 66) or DOX dependent (A375 clones 304, 401, 150, 1090) manner. The sgRNA used are listed in Table S5. Monoclonal cultures were generated by seeding n=1 (SK-LMS-1 and A375), n=3 (BJ/E1A-RAS) cells in each well of 96-well plates (Sarstedt). The successful generation of KO clones was screened by immunoblotting and confirmed by Sanger sequencing.

### Immunofluorescence and immunoblotting

Cells were fixed with 3% paraformaldehyde and permeabilized with 0.3% Triton X-100. The secondary antibodies were Alexa Fluor 488-, 546-or 633-conjugated anti-mouse and anti-rabbit secondary antibodies (Molecular Probes). Actin was labelled with phalloidin-AF546 or AF-660 (Molecular Probes). For the intracellular staining of dsRNA, the permeabilization step was performed for 5’ with 0.5% Triton X-100. ICC blocking solution (3% w/v BSA (Sigma-Aldrich), 3% v/v goat serum (Abcam), 0.02% v/v Tween-20 in PBS) was applied for 1h to block nonspecific binding and for the incubation at 37°C for 1h with 250 ng J2 antibody (Scicons). For S phase analysis, cells were grown for 3 h with 50 µM Bromodeoxyuridine (BrdU). After fixation, coverslips were treated with HCl (1% and 2%), quenched with Borate and processed for immunofluorescence. Cells were imaged with a confocal microscope Leica AOBS SP8 or with Leica AF6000 LX. Nuclei were stained with Hoechst 33258 or DAPI (Sigma-Aldrich).

Cell lysates after SDS-PAGE and immunoblotting on nitrocellulose (Whatman) were incubated with primary antibodies. HPR-conjugated secondary antibodies were obtained from Cell Signalling and blots were developed with Super Signal West Dura (Thermo Fisher Scientific). Primary and secondary antibodies were removed by using Restore PLUS Western Blot Stripping Buffer (Thermo Fisher Scientific), according to manufacturer. Unless otherwise indicated, all the immunoblot figures were representative of at least two biological replicates. The primary and secondary antibodies used in this work are listed in Table S6. Images represent maximum intensity projections of 3D image stacks and were adjusted for brightness and contrast for optimal visualization.

### Time-lapse video microscopy

SK-LMS-1 and SK-LMS-1/*HDAC4*^-/-^/*HDAC4*^*PAM*^*-ER* cells engineered to express H2B-GFP and Apple-53BP1 trunc or H2B-GFP alone were seeded on fibronectin coated 35mm Glass bottom dishes (MatTek) at low density (0.3*10^5^ cells). After 24h, medium was refreshed and 4-OHT was added in HDAC4 re-expressing cells. 6h later, the dishes were housed in the live cell imaging chamber of a Leica AOBS SP8 confocal microscopy, maintained in a humidified atmosphere at 37°C and 5%CO_2_ and imaged every 10 minutes for 74h under four dimensions. 5 z-stacks were collected for each time-point. Laser power, exposure time, pinhole aperture and acquisition intervals were chosen appropriately to minimize toxicity and bleaching.

### Proteomics and transcriptomics from *in vivo* murine aging models

C57BL/6J female mice were obtained from Shared Ageing Research Models (ShARM, UK). Tissues explanted from 4 months (128 days) and 26 months (774 days) old mice were snap-frozen in liquid nitrogen. For protein lysates generation, subsections of the liver and of the skin were grinded into a powder with a pestle and lysed for 1h at 4°C respectively with 400 and 200µl RIPA lysis buffer for 10mg of tissue. 4x Laemli sample buffer was added to the clarified lysates and after boiling the samples were loaded on SDS/PAGE gels. For RNA extraction, 1ml Tri Reagent (Molecular Research Center) was added to 10mg of smashed tissues. After 1h incubation at 4°C, RNA was recovered by phenol-chloroform extraction/ethanol precipitation and resuspended in 20 μL RNase-free water.

### Immunoprecipitation

Cells were lysed for 10’ into hypotonic lysis buffer (20mM Tris-HCl pH7.4, 10mM KCl, 10mM MgCl_2_, 1% Triton X-100, 10% glycerol, 50mM Iac, 1 mM phenylmethylsulphonylfluoride, 5 mM NaF, 1 mM Na_3_VO_4_), supplemented with protease inhibitors and 10µM MG-132 and 10µM G5. Lysates were incubated for 5 h with 1µg anti-HDAC4^78^ or rabbit IgG and for 1h with 30µl slurry protein A (GE). After 4 washes, the immunocomplexes were reversed with 2x Laemli sample buffer, boiled, resolved by SDS-PAGE and subjected to western-blotting. 1/100 of total lysate has been collected as input.

### dsRNA immunoprecipitation

Total RNA was extracted with Tri Reagent (Molecular Research Center) from a pellet of 35*10^6^ SK-LMS-1 HDAC4 KO cells. IP was performed O/N in Polysomal Lysis Buffer^79^ in the presence of 10μg J2 antibody (Scicons). 50µl slurry protein A (GE) was added and incubated in continuous rotation at 4°C for 4h. After 5 washes of the collected immunocomplexes, RNA was recovered by phenol-chloroform extraction/ethanol precipitation and resuspended in 20μL RNase-free water. The precipitation was repeated until a final amount of 1.6 µg of purified RNA was reached. 800ng were treated for 20’ at 37°C with 50u ShortCut Rnase III (NEB), in the digestion buffer supplied by the manufacturer. The digestion was stopped with 10x EDTA. The remaining 800ng were treated in the same manner but without the addition of RNAse III. 30pmoles of purified dsRNA treated or not with RNAseIII were transfected in recipient cells, by using 15µl Lipofectamine 3000 (LifeTechnologies) and 250µl Optimem (Gibco). The enrichment of dsRNA in the preparation was evaluated by qPCR and expressed as % of enrichment over input.

### SA-β-gal assay

Cells seeded on coverslips in 12-well plates were fixed for 5’ (PBS 2% formaldehyde/0.2% glutaraldehyde), washed twice with 0.9% NaCl and stained for 16h at 37°C with staining solution: 40 mM citric acid/Na phosphate buffer, 5 mM K4[Fe(CN)6]3H2O, 5 mM K3[Fe(CN)6], 150 mM sodium chloride, 2 mM magnesium chloride and 1 mg/mL X-gal (Panreact Applichem). Images were acquired with Leica LD bright field optical microscope.

### Transformation assay

Soft agar assay was performed as previously described^57,77^. Briefly, a total of 0.8*10^5^ cells were seeded in 0.3% top agar/DMEM layer above a 0.6% agar/DMEM basement. Fresh medium was added twice/week. After 15 days of culture the supernatant was discarded and the MTT [3-(4,5-dimethylthiazol-2-yl)-2,5-diphenyltetrazolium bromide] staining (0.5mg/ml in PBS) was applied for 2h. Images were acquired with a Leica DN6000 microscope. Foci were automatically counted with Clono Counter.

### RNA extraction and quantitative qRT-PCR

Cells were lysed using Tri Reagent (Molecular Research Center). 1.0 µg of total RNA was DNAseI treated (NEB #T2010) and retro-transcribed by using 100 units of M-MLV Reverse transcriptase (Life Technologies) in the presence of 1.6 µM oligo(dT) (Sigma-Aldrich) and 4 µM Random hexamers (Euroclone). qRT-PCRs were performed using SYBR green technology (KAPA Biosystems). Data were analyzed by comparative threshold cycle (delta delta Ct) using *HPRT* and *GAPDH* or *ACTB* and *GAPDH* as normalizer. The primers used for qRT-PCR are listed in Table S5.

### RNA array expression and data analysis

Total RNA was purified with Quick-RNA Miniprep (ZymoResearch), amplified according to the specifications of the Illumina TotalPrep RNA Amplification Kit (Ambion) and hybridized on Illumina whole-genome HumanHT-12 v 4.0 chip (Illumina). Acquisition and data analysis were performed as previously described^14^. Principal component analysis (PCA) was performed by using R function prcomp. Differentially expressed genes (DEGs) were called accordingly to the following criteria: |fold change|>2 and p adj.<0.05. The list of the DEGs is provided as Table S7. GSEA analysis in Fig.2 was performed as previously described^14^. The transcripts defining the “NFκβ”, “SASP”^80^ and “RIS up-regulated genes” gene sets are listed in Table S8. Gene list enrichment in Table S4 was performed by interrogating MSigDB collections (BP,C6,CGP,H,MF) with the transcripts associated to promoters (with 2kb from TSS), TEs or SEs bound by HDAC4; the obtained enrichments were considered significant for p and FDR <0.05 and if at least three Gene Sets fall in the same category. For the expression levels and Kaplan-Meier analysis of TCGA Skin Cutaneous Melanoma samples, data were retrieved from CBioPortal^81^ and expressed as z-score. Z-scores > |1.75| were selected as cut-off. For bioinformatics analysis in Fig.2, 7, S3 and S4, the following GEO datasets were analyzed: GSE38410, GSE74324, GSE40349, GSE3189, GSE78138, GSE45276, GSE36640, GSE40349, GSE132569.

### ChIP, library construction, ChIP-seq and NGS data analysis

Chromatin was obtained from SK-LMS-1 cells, 36h after or not HDAC4 removal, and immunoprecipitated with 2 µg of anti-H3K27ac, 3µg of anti-H3K27me3, 4µg of anti-LMNB1 and 4µg of anti-HDAC4 antibodies or control IgG, as previously described^14^. Three independent biological replicates were pulled according to BLUEPRINT requirements and 5 ng of total DNA were used to prepare ChIP-seq libraries, according to TruSeq ChIP Sample Preparation guide (Illumina). Libraries were sequenced on the Illumina HiSeq 2000 sequencer. The ShortRead R*/*Bioconductor package was used to evaluate the quality of sequencing reads and Bowtie 2 was used to align them to NCBI *GRCh38* human genome reference. Peak calling was performed against input sequences using HOMER for HDAC4 ChIP (“factor” mode) and MACS2 for H3K27ac and H3K27me3 (“sharp” mode and “broad” mode, respectively); gene annotations were performed as previously described^14^. gplots, biomaRt and Gviz R*/*Bioconductor packages and the deepTools suite were used to generate peak heatmaps and for the visualization of genomic loci. The H3K27ac and H3K27me3 enriched genomic regions between HDAC4 KO and wt were called according to 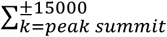 *f*(*k*), *where* 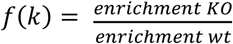. |log_2_ (Fc)|≥1 was used as cut-off. The following databases were consulted for the dissection of the ChIP-seq peaks as reported in Fig.6g/h, 7g/h, 8m, S3j/k: SEdb^82^ for super-enhancers, HERVd^83^ for endogenous retroviruses, HumCFS^84^ and the refinement^50^ for common fragile sites; ERFs^51^ and HK exons^85^. sample_01_066_SE of smooth muscle SE was used as lineage reference. “SES”, defined according to the ROSE algorithm^86^, represents the SEs activated during senescence. SES is the union of the SE identified during OIS^3^ and replicative senescence^7^ and is provided as Table S9. Liftover tool was used to convert genome coordinates between assemblies and to remap homologous sequences between genomes. The bedtools toolset (−-intersect option)^87^ was used to identify overlaps of at least one nucleotide. The investigated classes of genomic elements have been considered as separated, not redundant and not overlapping. The enrichment has been calculated with respect to the genome coverage of each genomic element (length of female haploid human genome: 3184709445 nucleotides). Known and novel motif discovery was performed using the MEME-ChIP tool from the MEME Suite^88^. The following parameters were used: -ccut 0; -order 1; -meme-maxsize 100000000; -meme-minsites 2; -meme-maxsites 100; -meme-minw 6; -meme-maxw 10; -meme-nmotifs 10; -meme-mod anr; -dreme-e 0.05; -centrimo-score 5.0; -centrimo-ethresh 10. The identified enriched motifs were compared to the Jolma2013, JASPAR2018_CORE_vertebrates_non_redundant and uniprobe_mouse databases for annotation. Enrichr (TRRUST) (http://amp.pharm.mssm.edu/Enrichr/) was used for the motif enrichment analysis in Table S2.

### Statistics

For experimental data, Student t-test was employed. Mann–Whitney test was applied when normality could not be assumed. *p*<0.05 was chosen as statistical limit of significance. For comparisons between more than two samples, the Anova test was applied coupled to Kruskal–Wallis and Dunn’s Multiple Comparison Test. For correlation between two variables, Pearson correlation or Spearman correlation were calculated for normal or non normal distributions, respectively. Excel and GraphPad Prism were used for routineer analysis, R/Bioconductor packages for large data analysis and heatmap generation. We marked with \**p*<0.05, \*\**p*<0.01, \*\*\**p*<0.001. Unless otherwise indicated, all the data in the figures were represented as arithmetic means ± the standard deviations from at least three independent experiments.

### Data availability

Raw data corresponding to ChIP-seq experiments are uploaded with GEO accession GSE149644. For reviewers, to access the data, https://www.ncbi.nlm.nih.gov/geo/query/acc.cgi?acc=GSE149644 Enter token clyvsuiwrlqfrgn into the box.

