## Supplementary material for "HDAC4 controls senescence and aging by safeguarding the epigenetic identity and ensuring the genomic integrity"

### Supplementary data

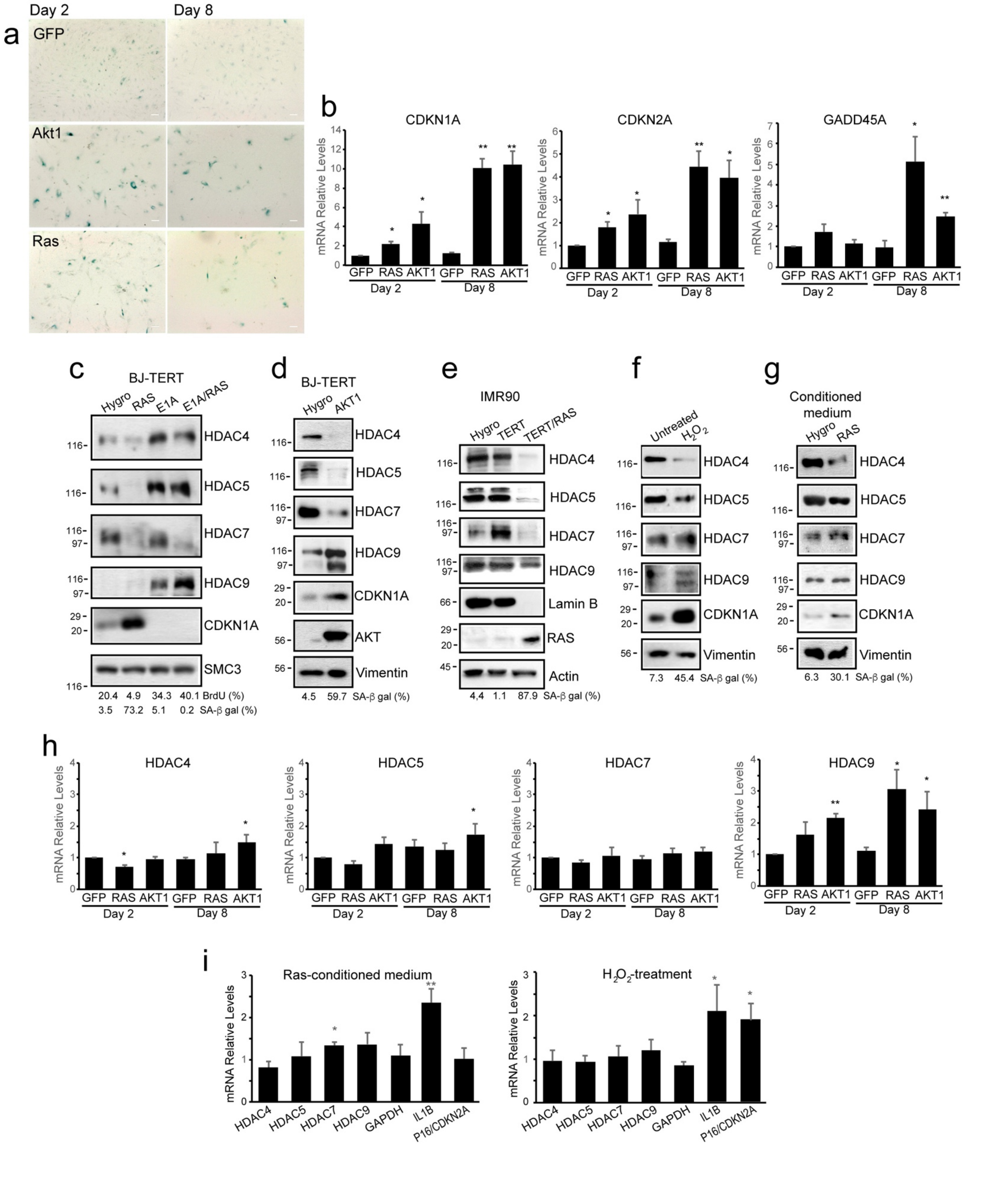

**Figure S1. Class IIa HDACs are dysregulated in different types of cellular senescence. a.** Microscopic images of SA-β-gal stained BJ/hTERT cells expressing the indicated transgenes for 2 or 8 days. Scale bar 50 µm. **b.** mRNA expression levels of the indicated genes in the indicated BJ/*hTERT* cells. Mean ± SD; n = 3. **c-g.** Immunoblot analysis of Class IIa HDACs, LMNB1, p21, RAS and AKT1 levels in the indicated cells stably expressing the indicated transgenes or treated with $H_2O_2$ (200µM for 2h and harvested after 4d) or the conditioned medium (1:1 conditioned/fresh medium, 2 treatments of 48h) obtained from BJ/hTERT cells stably expressing HYGRO$^R$ or HRAS$^{G12V}$. SA-β-gal and BrdU positivity are indicated. Actin, Vimentin and SMC3 were used as loading control. **h-i.** mRNA expression levels of the indicated genes in BJ/hTERT cells generated and treated as described in Fig.S1a, f, g. Mean ± SD; n = 3.

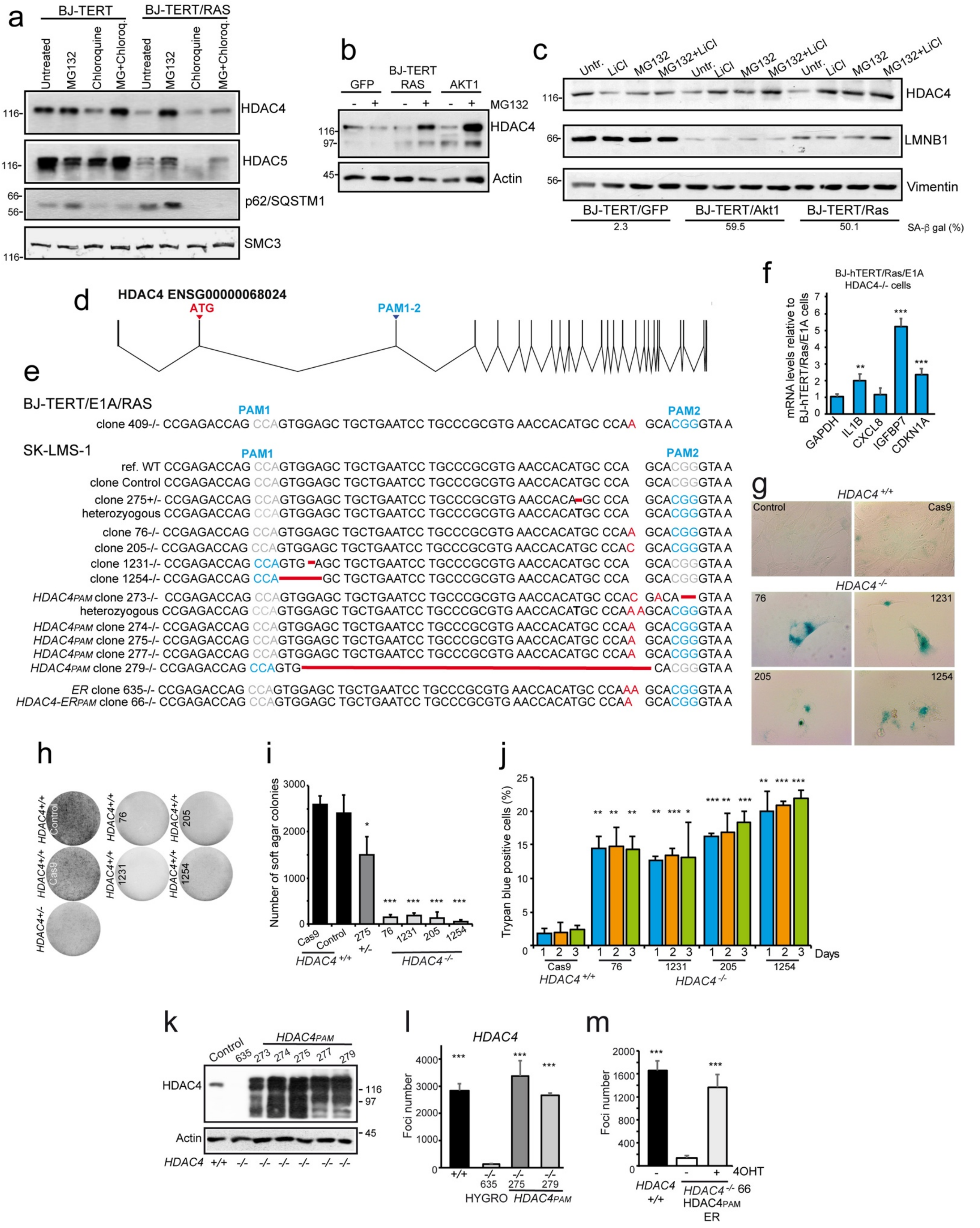

**Figure S2. The proteasomal mediated degradation or the CRISPR/Cas9 mediated targeting of HDAC4 allow premature senescence entrance. a.** Immunoblot analysis of HDAC4, HDAC5 and p62 in BJ/hTERT and BJ/*hTERT/HRAS*$^{G12V}$ expressing cells after 12 days of culture and 8h of treatment with MG132 (1 µM) and Cloroquine (10µM), as indicated. SMC3 was used as loading control. **b.** Immunoblot analysis of HDAC4 and Actin in the input (1/100 total lysate) related to the immunoprecipitation shown in Fig. 1e. **c.** Immunoblot analysis of HDAC4 and LMNB1 in BJ/hTERT cells expressing for 6 days the indicated transgenes and treated for the last 12h with LiCl (10mM) and/or MG132 (1µM), as indicated. Vimentin was used as loading control. **d.** Schematic representation of HDAC4 genomic organization with indicated: the exons (vertical bars), the introns (junctions between the bars) and the PAM sequences utilized for the CRISPR/Cas9 genome editing. **e.** Genomic sequences of the *HDAC4*$^{-/-}$ BJ/RAS/E1A and SK-LMS-1 cells used in this study. **f.** mRNA expression levels of the indicated genes in BJ/hTERT/RAS/E1A *HDAC4*$^{-/-}$ cells, relative to wt. Mean ± SD; n = 3. **g.** Microscopic images of SA-β-gal stained SK-LMS-1 clones, as indicated. **h.** Representative images of MTT stained foci formed from the indicated SK-LMS-1 clones and grown for 15 days in soft-agar. **i.** Quantification of the stained foci displayed in Fig. S2h. Data are presented as mean ± SD; n = 4. **j.** Histogram representing the percentage of TB positivity in the indicated SK-LMS-1 clones. Mean ± SD; n = 3. **k.** Immunoblot analysis of HDAC4 and Actin in the indicated SK-LMS-1 clones. Mean ± SD; n = 4. **l-m.** Histogram representing the quantification of the foci formed from the indicated SK-LMS-1 clones grown for 15 days in soft-agar. 250nM 4-OHT was added where indicated every three days. Mean ± SD; n = 3.

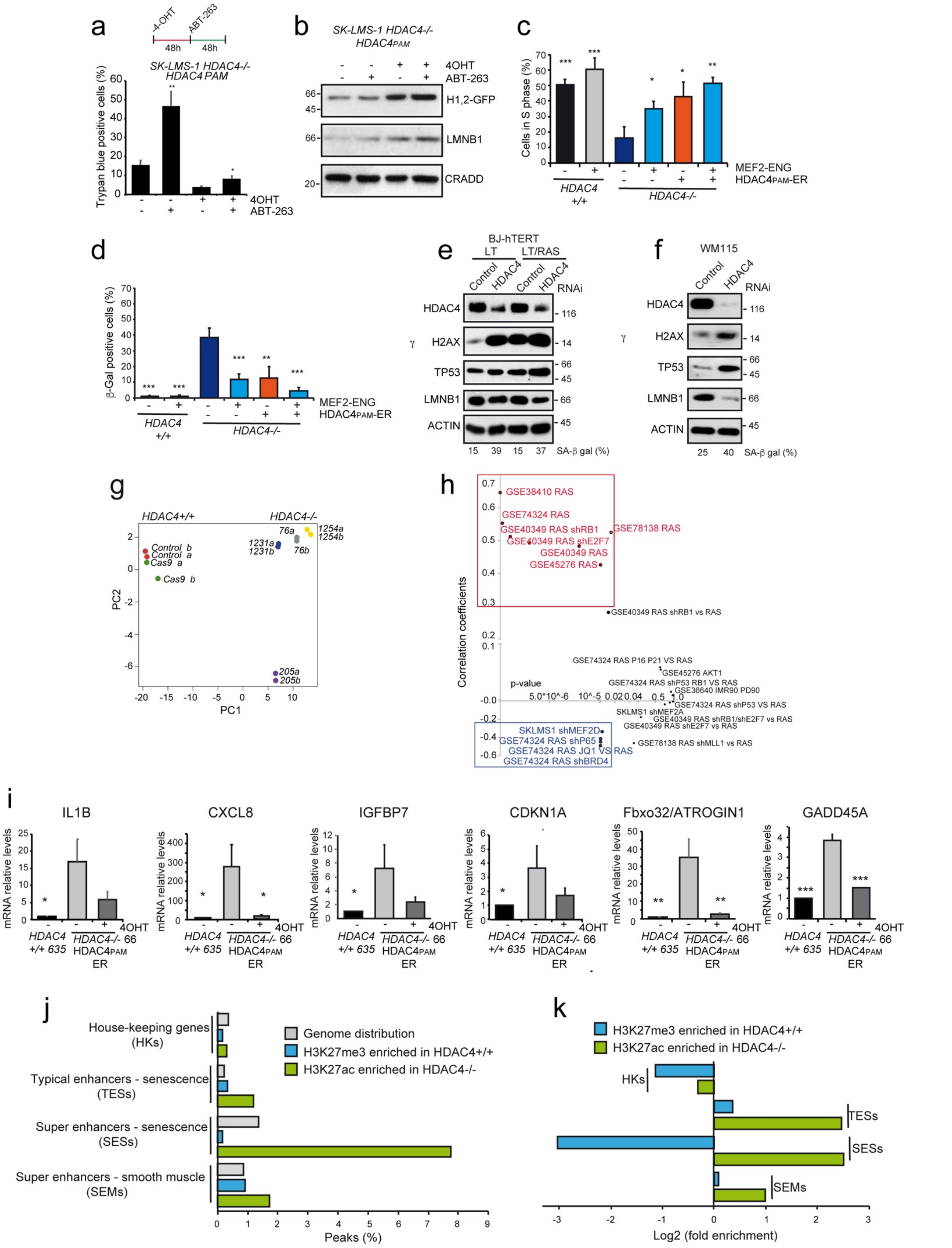

**Figure S3. Characterization of the pro-senescent properties arising from HDAC4 depletion. a.** Histogram representing the percentage of TB positivity in HDAC4 KO SK-LMS-1 cells, re-expressing or not HDAC4 and treated or not with

100nM ABT-263, as indicated. Mean ± SD; n = 3. **b.** Immunoblot analysis of HDAC4 and H1.2 GFP in the same cells described in Fig. S3a. CRADD was used as loading control. **c-d.** Histograms representing the percentage of BrdU (c) and SA-β-gal (d) positivity in HDAC4 KO SK-LMS-1 re-expressing HDAC4 or the super-repressive isoform of MEF2 (MEF2-ENG), as indicated. The significance is relative to the KO clone not re-expressing HDAC4. Mean ± SD; n = 4. **e-f.** Immunoblot analysis of HDAC4, γH2AX, TP53 and LMNB1 in the indicated cells transfected for 72h with siRNAs against HDAC4 or control. SA-β-gal positivity is indicated. Actin was used as loading control. **g.** PCA analysis performed on the expression profile of the indicated SK-LMS-1 clones. **h.** Dot plot representing the correlation coefficient (ρ, y-axis) and significance (p, x-axis) of SASP signature (as defined in materials and methods) between HDAC4 KO SK-LMS-1 cells and the indicated models of senescence. $|\rho|>0.3$ and $p<0.01$ were considered as significant. **i.** mRNA expression levels of the indicated genes in the indicated SK-LMS-1 clones. The significance is relative to the KO clone not re-expressing HDAC4. Mean ± SD; n = 3. **j.** Histogram representing the percentage of hyper-acetylated (green bar) or demethylated (light blue) H3K27 peaks in SK-LMS-1 *HDAC4*$^{-/-}$ cells and falling in genomic regions displaying the indicated epigenomic features. The genome coverage of each element is indicated by gray bars. **k.** Histogram representing the enrichment of each element described in Fig. 3j in respect to the expected distribution calculated according to the genome coverage.

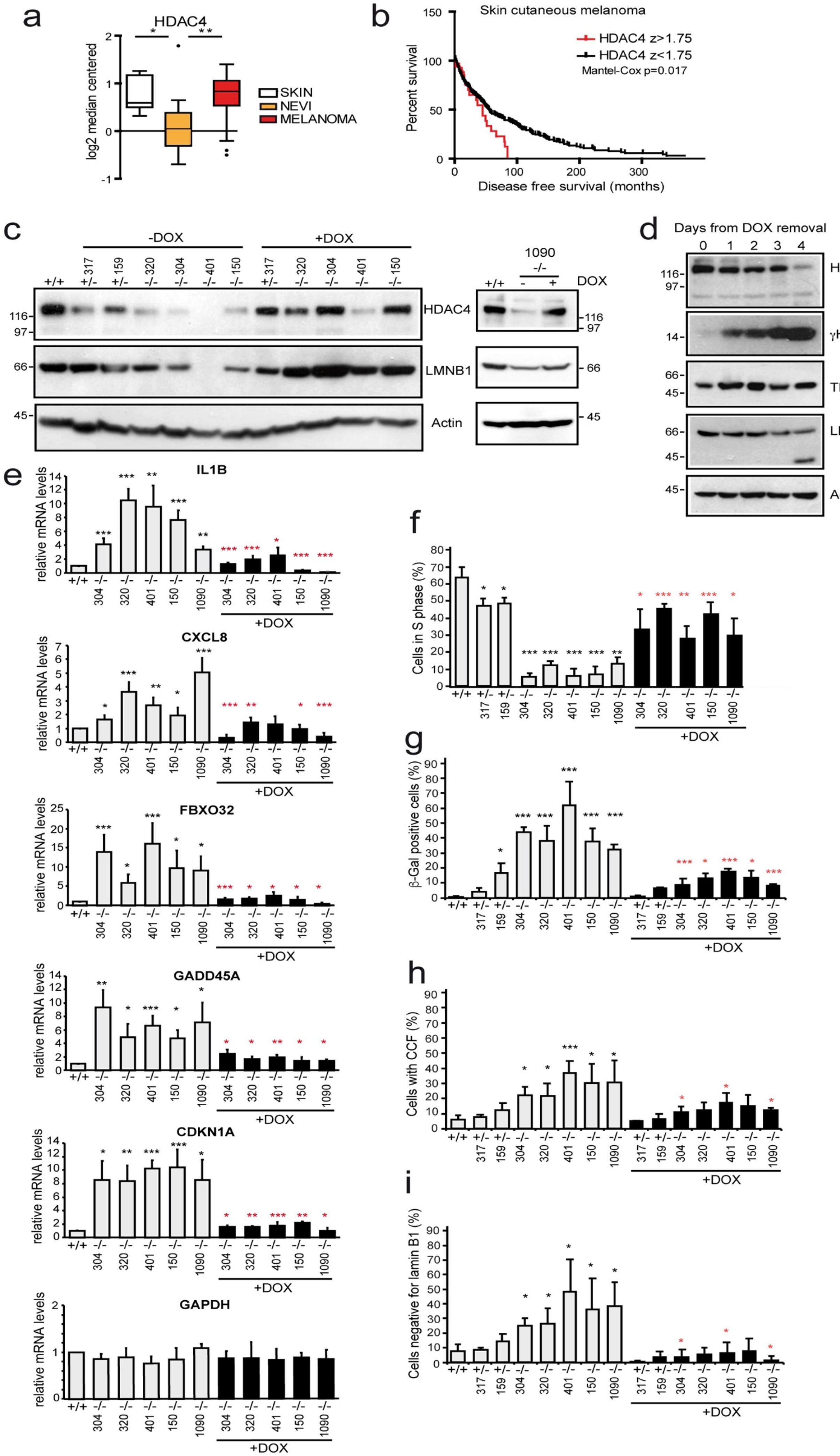
a
HDAC4
log2 median centered
SKIN
NEVI
MELANOMA
b
Skin cutaneous melanoma
HDAC4 z>1.75
HDAC4 z<1.75
Mantel-Cox p=0.017
Percent survival
Disease free survival (months)
c
-DOX
+DOX
1090
DOX
HDAC4
LMNB1
Actin
d
Days from DOX removal
HDAC4
γH2AX
TP53
LMNB1
Actin
e
IL1B
CXCL8
FBXO32
GADD45A
CDKN1A
GAPDH
relative mRNA levels
+DOX
f
Cells in S phase (%)
+DOX
g
β-Gal positive cells (%)
+DOX
h
Cells with CCF (%)
+DOX
i
Cells negative for lamin B1 (%)
+DOX

**Figure S4. Characterization of the role played by HDAC4 in sustaining senescence escape and melanomagenesis. a.** Box plot representing the log2 median centered levels of HDAC4 transcripts in normal skin (n=7), nevi (n=18) and melanoma (n=45) biopsies (GSE3189). **b.** Kaplan-Meier survival analysis related to the expression levels of HDAC4 (z-score>1.75, n=25) in 425 TCGA melanoma samples. Median months overall survival: 47.54 and 81.20 respectively in HDAC4 high and HDAC4 low clusters of patients. **c.** Immunoblot analysis of HDAC4 and LMNB1 in the indicated A375 clones, re-expressing or not a doxycycline (DOX)-inducible CRISPR/Cas9 resistant form of HDAC4 ($HDAC4^{PAM}$). Actin was used as loading control. **d.** Time-course immunoblot analysis of HDAC4, γH2AX, TP53 and LMNB1 levels in the A375 clone 1090, in lysates harvested at the indicated days after HDAC4 (DOX) removal. Actin was used as loading control. **e.** mRNA expression levels of the indicated genes in the indicated A375 clones, grown for 4d in the presence or absence of DOX. The significance is relative to wt cells (black *) or to clones grown in absence of DOX (red *). Mean ± SD; n = 4. **f-i.** Analysis of the percentage of cells displaying BrdU (f) positivity, SA-β-gal (g) positivity, chromatin cytoplasmic fragments (CCFs) (h) and altered or absent LMNB1 (i). The significance and cell treatments are as explained in Fig. S4e. Mean ± SD; n = 4.

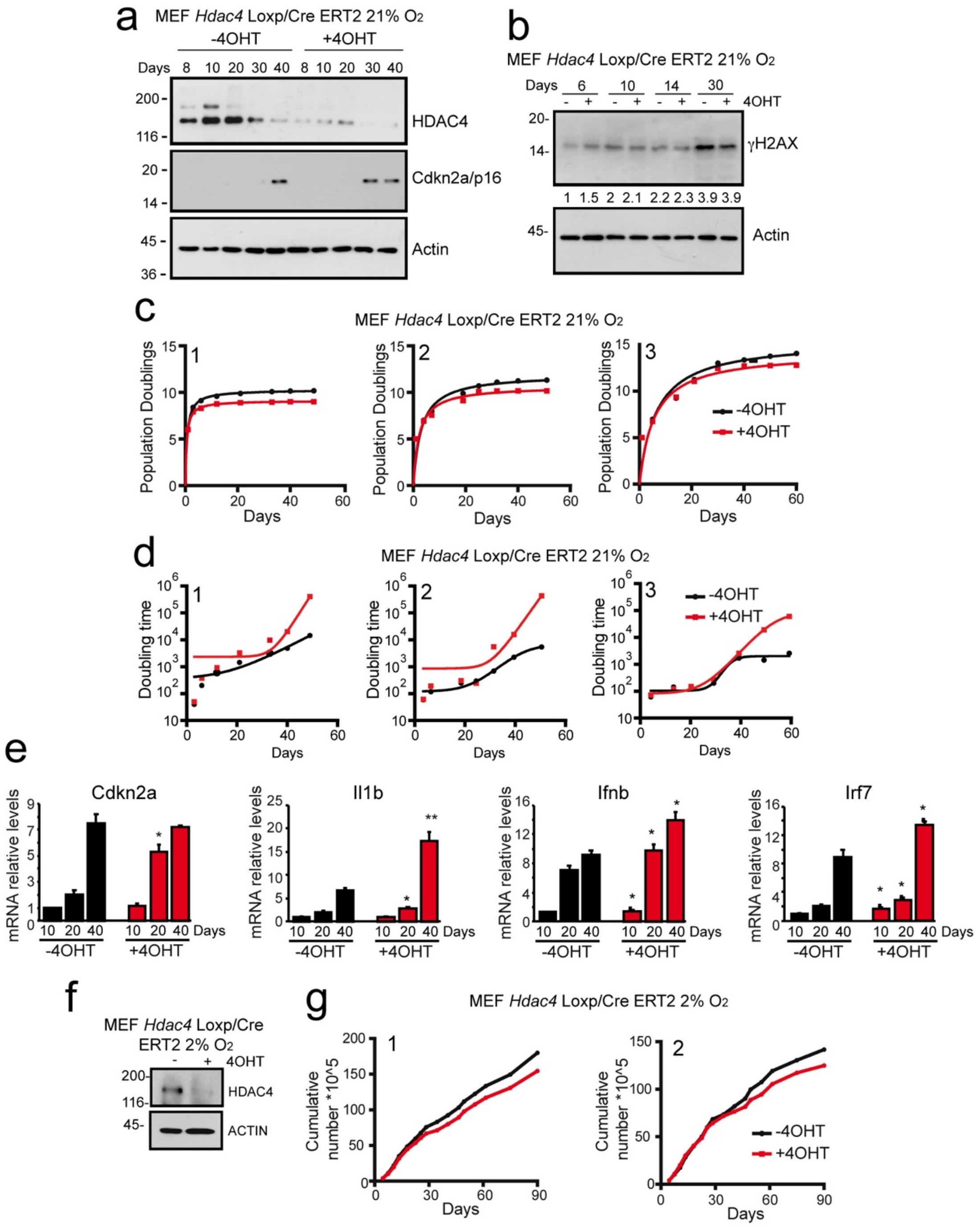

**Figure S5. HDAC4 depletion stimulates the premature senescence entrance of primary murine fibroblasts. a-b.** Immunoblot analysis of HDAC4 and p16 (a) and γH2AX (b) levels in MEF$^{loxp/loxp}$ x CreER T2 cells, maintained in normoxia and treated or not for 48h with 0.25µM 4-OHT at day 6 of culture and harvested after regular splitting at the indicated days. Actin was used as loading control. **c-d.** Analysis of the cumulative population doublings (c) or the doubling time (d) observed during the culture of thee independent batches of MEF$^{loxp/loxp}$ x CreER T2 cells maintained as described in Fig.S5a. Third order polynomial regression curves (c) and sigmoidal regression curves (d) are shown. **e.** mRNA expression levels of the same MEF cells described in Fig.S5a. Mean ± SD; n = 2. For significance calculation, a paired comparison between wt and *Hdac4*$^{-/-}$ cells was made for each growth point. **f.** Immunoblot analysis of HDAC4 levels in MEF$^{loxp/loxp}$ x CreER T2 cells, maintained in hypoxia and treated or not for 48h with 0.25µM 4-OHT at day 6 of culture and harvested after regular splitting at day 30. Actin was used as loading control. **g.** Analysis of the cumulative population doublings observed during the culture of two independent batches of MEF$^{loxp/loxp}$ x CreER cells maintained as described in Fig.8f.

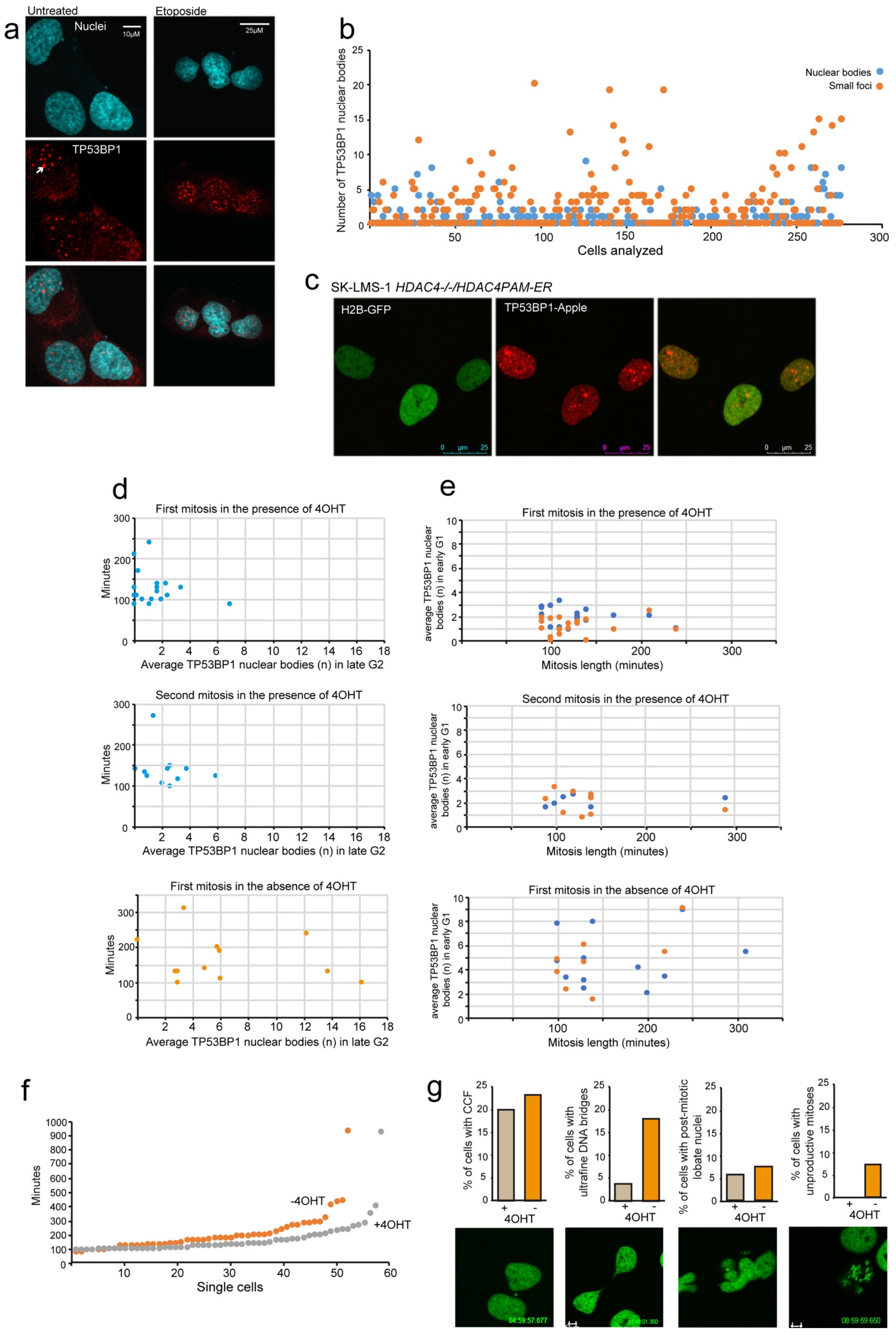
a
Untreated
Etoposide
Nuclei
10μM
25μM
TP53BP1
b
Number of TP53BP1 nuclear bodies
Cells analyzed
Nuclear bodies
Small foci
c
SK-LMS-1 HDAC4-/-/HDAC4PAM-ER
H2B-GFP
TP53BP1-Apple
d
First mitosis in the presence of 4OHT
Minutes
Average TP53BP1 nuclear bodies (n) in late G2
Second mitosis in the presence of 4OHT
First mitosis in the absence of 4OHT
e
average TP53BP1 nuclear bodies (n) in early G1
Mitosis length (minutes)
f
Minutes
Single cells
-4OHT
+4OHT
g
% of cells with CCF
% of cells with ultrafine DNA bridges
% of cells with post-mitotic lobate nuclei
% of cells with unproductive mitoses
4OHT

**Figure S6. Validation of the Apple-53BP1-trunc fluorescent sensor for the visualization of DNA damage foci and TP53BP1 bodies. a.** Representative confocal pictures of SK-LMS-1 untreated or Etoposide treated (2h, 20μM) cells immunostained with anti-53BP1 and secondary AF546 antibodies (red) to visualize endogenous TP53BP1 bodies and foci, while DAPI was used to stain nucleic acids (blue). The Arrow points to a TP53BP1 body. **b.** Dot plot representing the quantification of DNA damage foci and TP53BP1 bodies identified by immunostaining of Etoposide treated SK-LMS-1 cells (n=270). **c.** Representative video frame of untreated SK-LMS-1/*HDAC4*$^{-/-}$ cells re-expressing HDAC4 engineered to stably express H2B-GFP and Apple-TP53BP1 trunc and subjected to *in vivo* video microscopy. **d.** Dot plot representing the correlation between the duration of the first and second mitosis (y) and the average number of TP53BP1 nuclear bodies (x) observed in the last 10 frames (100 minutes) preceding the prophase in SK-LMS-1/*HDAC4*$^{-/-}$ cells re-expressing (+4-OHT) or not (-4-OHT) HDAC4-ER. The observation period is of 74h. Mitosis duration: +4-OHT: 1st 127 ± 39.34 min; 2nd 139.09±53.37; -4-OHT: 1st 166.67±65.55; 2nd not observed. **e.** Dot plot representing the correlation between the duration of the first and second mitosis (x) and the average number of TP53BP1 nuclear bodies (y) observed during the first 10 frames (100 minutes) after cytokinesis in sister cells (orange spot: mother cells, blue spot: offspring) subjected to the same observation explained in Fig. S6d. **f.** Dot plot representing the duration of mitosis in SK-LMS-1/*HDAC4*$^{-/-}$ cells re-expressing *HDAC4*$^{PAM}$*-ER* (+4-OHT) or not (-4-OHT). **g.** Quantification of the mitotic defects evidenced by H2B-GFP and observed in the cells subjected to *in vivo* video microscopy as explained in Fig. S6d, during 74h of analysis. For each kind of mitotic defect, a representative video frame is provided.

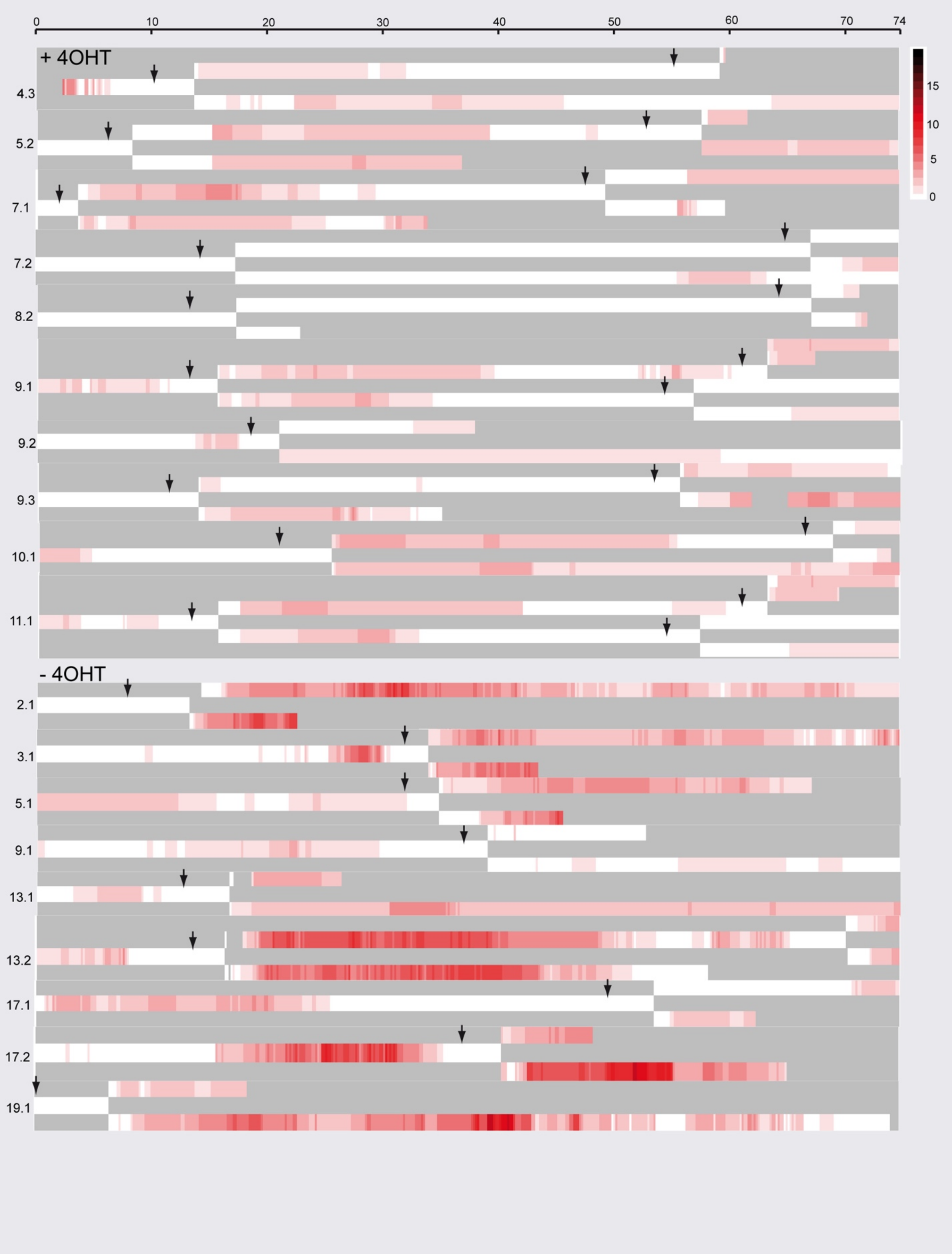

**Figure S7. The progressive accumulation of TP53BP1 bodies in HDAC4 depleted cells is correlated to mitotic slowdown and impairment.** Heatmap representing the quantification of the TP53BP1 bodies in SK-LMS-1/*HDAC4*$^{-/-}$/*HDAC4*$^{PAM}$*-ER* cells, during 74h of analysis starting 6h after 4-OHT removal (time "0"), as indicated. The intensity of the red is proportional to the number of TP53BP1 bodies. 10 and 9 starting cells were analyzed respectively for the +4-OHT and the -4-OHT conditions.

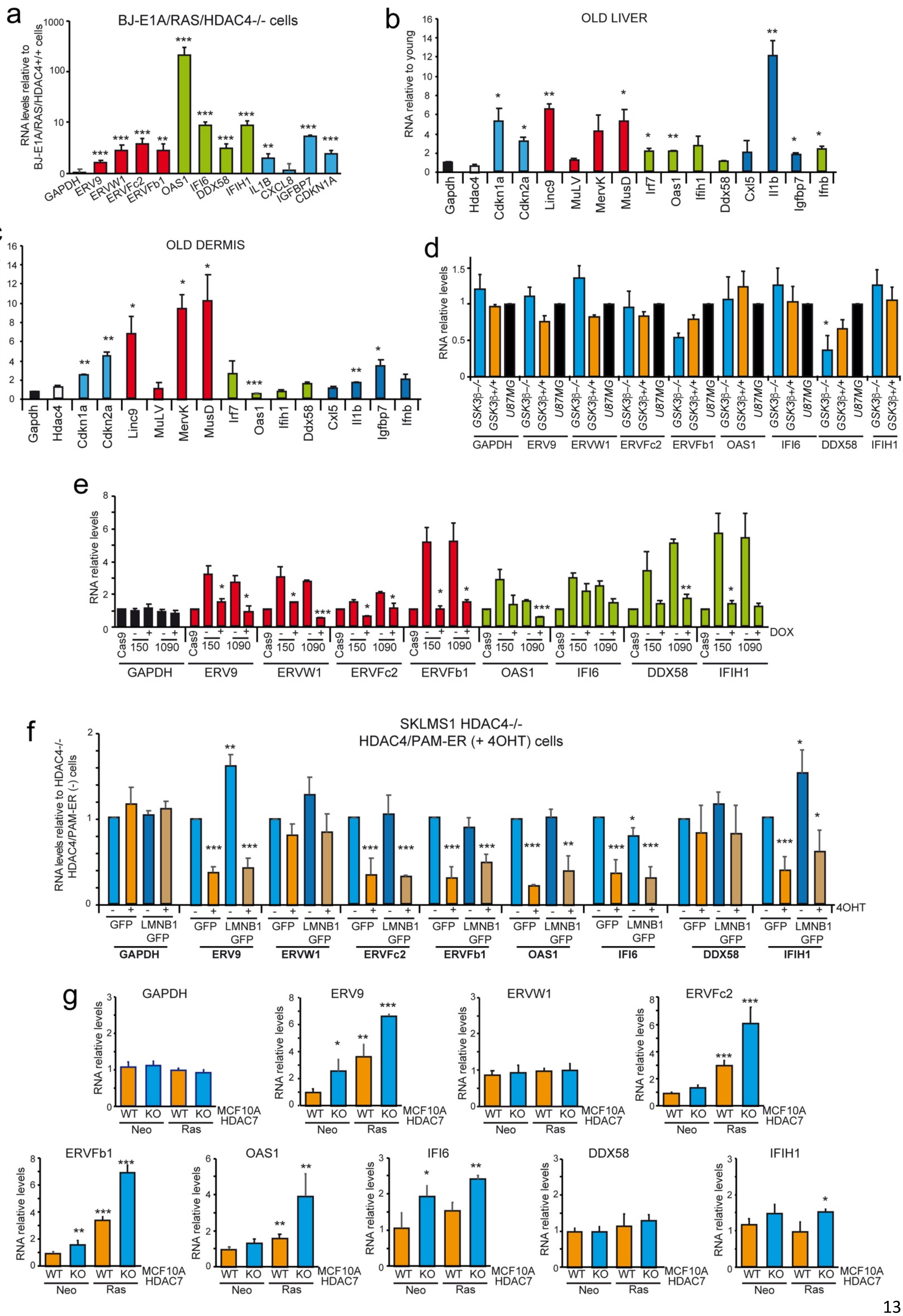
a
BJ-E1A/RAS/HDAC4-/- cells
RNA levels relative to BJ-E1A/RAS/HDAC4+/+ cells
GAPDH ERV9 ERVW1 ERVFc2 ERVFb1 OAS1 IFI6 DDX58 IFIH1 IL1B CXCL8 IGFBP7 CDKN1A
b
OLD LIVER
RNA relative to young
Gapdh Hdac4 Cdkn1a Cdkn2a Linc9 MuLV MervK MusD Irf7 Oas1 Ifih1 Ddx58 Cxl5 Il1b Igfbp7 Ifnb
c
OLD DERMIS
RNA relative to young
Gapdh Hdac4 Cdkn1a Cdkn2a Linc9 MuLV MervK MusD Irf7 Oas1 Ifih1 Ddx58 Cxl5 Il1b Igfbp7 Ifnb
d
RNA relative levels
GSK3β-/- GSK3β+/+ U87MG
GAPDH ERV9 ERVW1 ERVFc2 ERVFb1 OAS1 IFI6 DDX58 IFIH1
e
RNA relative levels
Cas9 150 1090
DOX
GAPDH ERV9 ERVW1 ERVFc2 ERVFb1 OAS1 IFI6 DDX58 IFIH1
f
SKLMS1 HDAC4-/-
HDAC4/PAM-ER (+ 4OHT) cells
RNA levels relative to HDAC4-/- HDAC4/PAM-ER (-) cells
4OHT
GFP LMNB1 GFP
GAPDH ERV9 ERVW1 ERVFc2 ERVFb1 OAS1 IFI6 DDX58 IFIH1
g
GAPDH ERV9 ERVW1 ERVFc2 ERVFb1 OAS1 IFI6 DDX58 IFIH1
RNA relative levels
WT KO WT KO
Neo Ras
MCF10A HDAC7

**Figure S8. ERVs regulation is HDAC4-dependent but LMNB1 and CRISPR/Cas9 independent. a.** RNA expression levels of the indicated genes and ERVs in BJ-E1A/RAS /*HDAC4*$^{-/-}$ cells in respect to wt cells. Mean ± SD; n = 3. **b-c.** RNA expression levels of the indicated genes and ERVs in the liver and in the dermis of aged mice, in respect to young mice, as explained in Fig.1c. Mean ± SD; n = 2. **d.** RNA expression levels of the indicated genes and ERVs in U87-MG/*GSK3β*$^{+/+}$ or *GSK3β*$^{-/-}$ generated through a CRISPR/Cas9 editing approach, in respect to normal U87-MG not edited cells. Mean ± SD; n = 3. **e.** RNA expression levels of the indicated genes and ERVs in A375 *HDAC4*$^{-/-}$ cells (clones 150 and 1090), re-expressing or not HDAC4$^{PAM}$ (+DOX), in respect to wt cells transfected with Cas9. Mean ± SD; n = 3. **f.** RNA expression levels of the indicated genes and ERVs in SK-LMS-1/*HDAC4*$^{-/-}$ cells, re-expressing *HDAC4*$^{PAM}$*-ER* (+4-OHT), GFP or LMNB1-GFP as indicated. The significance is relative to *HDAC4*$^{-/-}$ cells expressing the GFP. Mean ± SD; n = 3. **g.** RNA expression levels of the indicated genes and ERVs in MCF10A/Neo$^{R}$ and MCF10A/HRAS$^{G12V}$ *HDAC7*$^{-/-}$ cells in respect to MCF10A/Neo$^{R}$ wt cells. Mean ± SD; n = 3.

| Gene | IMR90 PD44 vs PD8 | BJ/hTERT RAS vs Hygro | BJ/hTERT E1A vs Hygro | BJ/hTERT RAS E1A vs Hygro |
|---|---|---|---|---|
| HDAC4 | 0.92±0.19 | 1.16±0.29 | 4.18±0.74 (**) | 3.21±0.24 (***) |
| HDAC5 | 1.56±0.28 (*) | 0.81±0.13 | 2.97±0.19 (***) | 2.42±0.49 (**) |
| HDAC7 | 1.26±0.3 | 0.85±0.11 | 1.87±0.36 (*) | 1.07±0.39 |
| HDAC9 | 5.16±0.89 (**) | 3.45±0.42 (***) | 8.67±0.98 (***) | 12.06±1.43 (***) |

**Table S1. mRNA expression levels of class IIa HDACs in replicative senescence, OIS and OIS escape.** Data are presented as mean ± SD, n=3. *p<0.05, **p<0.01, ***p<0.005.

| Term | Overlap | P-value | Odds Ratio | Combined Score |
|---|---|---|---|---|
| NFKBIA | 21,05 | 1,53E-07 | 80,97 | 1270,7 |
| RELA | 2,66 | 1,02E-06 | 10,22 | 141,07 |
| JUN | 4,03 | 2,37E-06 | 15,49 | 200,6 |
| NFKB1 | 2,31 | 1,28E-05 | 8,89 | 100,14 |
| REL | 13,64 | 2,47E-05 | 52,45 | 556,5 |
| FOS | 5,26 | 4,39E-04 | 20,24 | 156,49 |
| STAT3 | 2,82 | 5,06E-04 | 10,83 | 82,22 |
| CEBPB | 5 | 5,11E-04 | 19,23 | 145,76 |
| HDAC1 | 4,23 | 8,36E-04 | 16,25 | 115,16 |
| ETS1 | 3,8 | 0,00114 | 14,61 | 98,97 |
| IKBKB | 16,67 | 0,0155 | 64,1 | 267,11 |
| MEF2A | 14,29 | 0,01806 | 54,95 | 220,55 |
| NCOA3 | 14,29 | 0,01806 | 54,95 | 220,55 |
| MAZ | 12,5 | 0,02062 | 48,08 | 186,62 |
| ZFP36 | 12,5 | 0,02062 | 48,08 | 186,62 |
| FOSL1 | 11,11 | 0,02316 | 42,74 | 160,91 |
| GATA6 | 11,11 | 0,02316 | 42,74 | 160,91 |
| ZNF148 | 11,11 | 0,02316 | 42,74 | 160,91 |
| FOXA1 | 10 | 0,0257 | 38,46 | 140,81 |
| NFIL3 | 10 | 0,0257 | 38,46 | 140,81 |
| IRF8 | 9,09 | 0,02824 | 34,97 | 124,72 |
| PAX6 | 9,09 | 0,02824 | 34,97 | 124,72 |
| NR5A2 | 8,33 | 0,03077 | 32,05 | 111,58 |
| SF1 | 8,33 | 0,03077 | 32,05 | 111,58 |
| TCF7L2 | 8,33 | 0,03077 | 32,05 | 111,58 |
| HDAC7 | 7,69 | 0,03329 | 29,59 | 100,67 |
| ING4 | 6,67 | 0,03831 | 25,64 | 83,64 |
| JUNB | 6,67 | 0,03831 | 25,64 | 83,64 |
| NR4A1 | 6,67 | 0,03831 | 25,64 | 83,64 |
| NFE2L2 | 6,25 | 0,04081 | 24,04 | 76,89 |
| HMGA1 | 5,88 | 0,04331 | 22,62 | 71,03 |
| DDIT3 | 5,56 | 0,0458 | 21,37 | 65,89 |
| FOXA2 | 5,56 | 0,0458 | 21,37 | 65,89 |
| XBP1 | 5,26 | 0,04828 | 20,24 | 61,35 |

**Table S2. List of TFs predicted to regulate the transcripts associated to SEs activated and hyper-acetylated in SK-LMS-1/*HDAC4*$^{-/-}$ cells.**

| Genomic coordinates | Motif name | Sequence | Position | Strand | Score | p-value | E-value |
|---|---|---|---|---|---|---|---|
| chr15:32850157-32850480 | MEF2A(MA0052.1) | CTATTCTTAG | 229 | - | 6,82 | 0,0009 | 0,283 |
| chr18:29154734-29155057 | MEF2A(MA0052.1) | CTATATTTAG | 194 | + | 11,85 | 5,00E-05 | 0,0157 |
| chr3:55169686-55170009 | MEF2A(MA0052.1) | CTATTTATAT | 78 | - | 8,22 | 0,000525 | 0,165 |
| chr12:75690734-75691057 | MEF2A(MA0052.1) | CTACTTTTAG | 304 | + | 10,34 | 0,000125 | 0,039 |
| chr1:186974870-186975193 | MEF2A(MA0052.1) | CTATTTTTAG | 83 | - | 13,49 | 5,00E-05 | 0,0157 |
| chr4:156778618-156778941 | MEF2A(MA0052.1) | CTACTTATAA | 19 | + | 7,16 | 0,00075 | 0,236 |
| chr5:111997815-111998138 | MEF2A(MA0052.1) | CTTTTTATAG | 9 | + | 7,49 | 0,000625 | 0,196 |
| chr6:142885458-142885781 | MEF2A(MA0052.1) | CTACATTTAG | 192 | + | 8,7 | 0,0003 | 0,094 |
| chr20:7856752-7857075 | MEF2A(MA0052.1) | CTTTTTATAG | 47 | - | 7,49 | 0,000625 | 0,196 |
| chr10:33129569-33129892 | MEF2A(MA0052.1) | CTATTTTTAA | 8 | - | 8,75 | 0,0003 | 0,094 |
| chr9:3875315-3875638 | MEF2A(MA0052.1) | CGATTTTTAG | 191 | - | 6,67 | 0,0009 | 0,283 |
| chr20:51360531-51360854 | MEF2A(MA0052.1) | CTATTTTTAA | 162 | + | 8,75 | 0,0003 | 0,094 |
| chr5:151063281-151063604 | MEF2A(MA0052.1) | CTATTTTTAA | 126 | + | 8,75 | 0,0003 | 0,094 |
| chr8:66130020-66130343 | MEF2A(MA0052.1) | CTATTTTTGG | 188 | - | 10,49 | 0,0001 | 0,0314 |
| chr8:66130020-66130343 | MEF2A(MA0052.1) | CTATTTATTG | 16 | + | 7,52 | 0,000625 | 0,196 |
| chr2:217239656-217239979 | MEF2A(MA0052.1) | CTATTTTTAA | 42 | - | 8,75 | 0,0003 | 0,094 |

**Table S3. List of 16 putative MEF2 binding sites in 15 out of 93 SESs bound by HDAC4.**

| PROMOTER | | |
|---|---|---|
| *EXTRACELLULAR MATRIX REMODELLING* | *p-value* | *FDR* |
| GO collagen catabolic process | 4,17E-11 | 5,3E-07 |
| GO extracellular matrix disassembly | 1,22E-09 | 3,78E-06 |
| GO extracellular structure organization | 1,48E-09 | 3,78E-06 |
| GO metalloendopeptidase activity | 4,94E-09 | 8,34E-06 |
| *INFLAMMATION* | *p-value* | *FDR* |
| Phong TNF response not via p38 | 3,39E-07 | 0,000359 |
| GO cellular response to virus | 4,66E-05 | 0,0147 |
| Dasu IL6 signaling up | 7,52E-05 | 0,0197 |
| GO regulation of defense response to virus | 0,000134 | 0,0266 |
| *TUMORIGENESIS AND SENESCENCE* | *p-value* | *FDR* |
| Cromer tumorigenesis up | 1,11E-06 | 0,000971 |
| GO negative regulation of cell growth | 0,000107 | 0,0231 |
| Ly aging middle up | 0,000156 | 0,0296 |
| Markey RB1 acute LOF dn | 0,000247 | 0,037 |
| *CELL DEATH* | *p-value* | *FDR* |
| Dutta apoptosis via NFKB | 9,66E-06 | 0,0057 |
| GO cysteine endopeptidase activity | 2,44E-05 | 0,0101 |
| Gali TP53 targets apoptotic up | 4,82E-05 | 0,0147 |
| GO negative regulation of necrotic cell death | 0,000357 | 0,0452 |
| **TE** | | |
| *POLYCOMB TARGETS* | *p-value* | *FDR* |
| Benporath SUZ12 targets | 1,52E-18 | 1,94E-14 |
| Benporath ESC with H3K27me3 | 7,05E-16 | 4,49E-12 |
| Meissner brain HCP with H3K4me3/H3K27me3 | 1,46E-14 | 6,19E-11 |
| Mikkelsen MEF HCP with h3k27me3 | 7E-07 | 0,00011 |
| *SIGNAL TRANSDUCTION* | *p-value* | *FDR* |
| GO molecular transducer activity | 1,2E-12 | 2,24E-09 |
| GO cell cell signaling | 2,11E-12 | 3,36E-09 |
| GO positive regulation of signaling | 1,38E-09 | 7,33E-07 |
| GO regulation of system process | 1,55E-09 | 7,67E-07 |
| *CELLULAR ADHESION* | *p-value* | *FDR* |
| GO biological adhesion | 6,73E-13 | 2,08E-09 |
| GO regulation of cell adhesion | 5,78E-09 | 2,16E-06 |
| GO positive regulation of cell adhesion | 1,7E-08 | 5,29E-06 |
| *TUMORIGENESIS AND SENESCENCE* | *p-value* | *FDR* |
| KRAS 600 up in lung and breast cancer | 2,45E-07 | 4,88E-05 |
| Perez TP53 targets | 3,13E-07 | 5,66E-05 |
| KRAS 600 upregulated genes | 1,19E-06 | 0,000167 |
| **SE** | | |
| *EXTRACELLULAR MATRIX REMODELLING* | *p-value* | *FDR* |
| GO collagen catabolic process | 4,09E-08 | 9,21E-05 |
| GO extracellular matrix disassembly | 1,18E-07 | 0,000187 |
| GO extracellular structure organization | 5,4E-07 | 0,000471 |
| GO metallopeptidase activity | 6,11E-06 | 0,00299 |
| *NFKB SIGNALLING* | *p-value* | *FDR* |
| Hinata NFKB targets keratinocyte up | 1,72E-07 | 0,000212 |
| Hinata NFKB targets fibroblast up | 1,84E-06 | 0,00118 |
| Hallmark TNFA signaling via NFKB | 1,42E-05 | 0,00549 |
| Tian TNF signaling via NFKB | 3,25E-05 | 0,0104 |
| *TUMORIGENESIS AND SENESCENCE* | *p-value* | *FDR* |
| Cromer tumorigenesis up | 1,83E-07 | 0,000212 |
| Chicas RB1 targets senescent | 6,94E-07 | 0,00052 |
| Sabates colorectal adenoma up | 7,61E-06 | 0,00346 |
| Osman bladder cancer dn | 5,48E-05 | 0,0148 |
| *INFLAMMATION* | *p-value* | *FDR* |
| Mahajan response to IL1A up | 1,42E-06 | 0,00101 |
| Browne HCMV infection 1hr up | 1,85E-06 | 0,00118 |
| Phong TNF targets up | 5,79E-06 | 0,00295 |
| Dasu IL6 signaling up | 7,39E-05 | 0,0188 |

**Table S4. Functional analysis of the nearest genes associated to HDAC4 binding in SK-LMS-1 cells.**

**Table S5. List of the primers used for cloning, genome editing and PCR.**

| Antibody | Catalogue or reference | Company | Application |
|---|---|---|---|
| RPA32 | sc56770 | Santa Cruz Biotechnology | WB |
| PCNA | sc56770 | Santa Cruz Biotechnology | WB |
| MAVS | sc166583 | Santa Cruz Biotechnology | WB |
| LMNA/C | sc376248 | Santa Cruz Biotechnology | IF |
| LMNB1 | sc377000 | Santa Cruz Biotechnology | IF |
| RACK1 | sc17754 | Santa Cruz Biotechnology | WB |
| GFP | Paroni, 2004[79] | Home-made | WB,IP |
| HDAC4 | Paroni, 2004[79] | Home-made | ChIP,IF,IP,WB |
| H3 | H0164 | Sigma-Aldrich | WB |
| H3K27ac | ab177178 | Abcam | ChIP,WB |
| H3K27me3 | ab6002 | Abcam | ChIP,WB |
| RPA32(pS4/S8) | A300-245A | Bethyl Laboratories | WB |
| HDAC5 | Clocchiatti, 2016[73] | Home-made | WB |
| HDAC7 | Cutano, 2018 | Home-made | ChIP,WB |
| HDAC9 | Di Giorgio, 2020[14] | Home-made | ChIP,WB |
| ACTB | #4970 | Cell Signalling Technology | WB |
| H2AX (pS139) | #9718 | Cell Signalling Technology | IF,WB |
| AKT1 | #2938 | Cell Signalling Technology | WB |
| HRAS | ab97488 | Abcam | WB |
| Ub | MMS-257P-200 | Covance | WB |
| VIM | sc-6260 | Santa Cruz Biotechnology | WB |
| p21 | P1484 | Sigma-Aldrich | WB |
| p53 | DO-1 | Home-made | WB |
| GSK3B | sc81462 | Santa Cruz Biotechnology | WB |
| E1A | sc-58658 | Santa Cruz Biotechnology | WB |
| SMC3 | A300-060A | Bethyl Laboratories | WB |
| 53BP1 | #88439 | Cell Signalling Technology | IF,WB |
| dsRNA | J2-1909 | Scicons | IF,IP |
| FLAG (M2) | F1804 | Sigma-Aldrich | WB |
| p16 | #92803 | Cell Signalling Technology | WB |
| p62/SQSTM1 | A302-856A | Bethyl Laboratories | WB |
| CRADD | #4899 | Cell Signalling Technology | WB |

**Table S6. List of the antibodies used.**

**Table S7. List of 230 DEGs identified by comparing wt and HDAC4KO SK-LMS-1 transcriptomes.**

**Table S8. List of the genes belonging to the NFκβ, SASP and RIS signatures interrogated in this paper.**

**Table S9. .bed formatted file of 1053 SESs, identified in replicative senescence[7] and OIS[3].**

**Video S1-2. Time-lapse videomicroscopy of SK-LMS-1/*HDAC4*$^{-/-}$/*HDAC4*$^{PAM}$-ER cells, over-expressing H2B-GFP and Apple-53BP1 and re-expressing (video S1) or not (video S2) HDAC4.** Each frame was acquired every 10 minutes for 74h.

| Primer name |
|---|
| APPLE BAMHI FW |
| 53BP1 TRUNC ECORI RV |
| 53BP1 TRUNC XHOI RV |
| MYR FW ECORI |
| AKT1 RV ECORI |
| MYR FW NHEI |
| AKT1 RV XBAI |
| AKT1 RV SALI NO STOP |
| AKT1 STOP RV BAMHI |
| Cas9 BAMHI FW |
| Cas9 ECORI RV |
| NLS RV ECORI |
| E1A FW NHEI |
| E1A RV BAMHI STOP |
| E1A MLUI FW |
| E1A 143 BGLII RV STOP |
| E1A 143 SALI RV STOP |
| GFP FW MLUI |
| GFP BGLII RV |
| H1.2 ECORI FW |
| H1.2 BGLII RV |
| hHDAC4 ECORI FW |
| hHDAC4 ECORI RV |
| HDAC4 MLUI FW |
| ERa ECORI RV STOP |
| ERa SAL RV NO STOP |
| HRAS FW NHEI |
| HRAS RV BAMHI |
| HRAS BAMHI FW |
| HRAS XHOI RV no stop |
| LMNB1 FW BAMHI |
| LMNB1 RV SALI NO STOP |
| BGLII FLAG AGEI MAVS FW |
| MAVSdCARD AGEI FW |
| MAVS SAL RV STOP |
| MAVSdCARD FW BGLII |
| MAVS RV NO STOP SALI |
| PURO FW NCOI |
| PURO RV SACI |
| FLAG FW BAMHI |
| FLAG SNABI FW |
| RIG1 XHOI RV |
| sgRNA1 HDAC4 FW |
| sgRNA1 HDAC4 RV |
| sgRNA2 HDAC4 FW |
| sgRNA2 HDAC4 RV |
| sgRNA4 HDAC4 FW |
| sgRNA4 HDAC4 RV |
| sgRNA5 HDAC4 FW |
| sgRNA5 HDAC4 RV |

| |
|---|
| HDAC4 P16A FW |
| HDAC4 P16A RV |
| HDAC4 P22A FW |
| HDAC4 P22A RV |
| HDAC4 V31L FW |
| HDAC4 V31L RV |

| Primer name |
|---|
| hACTB FW rt |
| hACTB RV rt |
| hCDKN1A RT FW |
| hCDKN1A RT RV |
| hCDKN2A FW RT |
| hCDKN2A RV RT |
| hCSF2 RT FW |
| hCSF2 RT RV |
| hDDX58 RT FW |
| hDDX58 RT RV |
| hERV9-1(pol) FW |
| hERV9-1(pol)RV |
| hERV-Fb1(env) FW |
| hERV-Fb1(env) RV |
| hERV-Fc2 FW |
| hERV-Fc2 RW |
| hERVW1(Syncitin1)FW |
| hERVW1(Syncitin1)RV |
| hFBXO32 FW |
| hFBXO32 RV |
| hFOXO1 RT FW |
| hFOXO1 RT RV |
| hGADD45a FW |
| hGADD45a RV |
| hGAPDH RT fw |
| hGAPDH RT RV |
| hHDAC4 RT FW |
| hHDAC4 RT RV |
| hHDAC5 RT FW |
| hHDAC5 RT RV |
| hHDAC7 RT FW |
| hHDAC7 RT RV |
| hHDAC9 RT FW |
| hHDAC9 RT RV |
| hHPRT RT FW |
| hHPRT RT RV |
| hIFI6 RT FW |
| hIFI6 RT RV |
| hIFIH1 RT FW |
| hIFIH1 RT RV |
| hIL1B RT FW |
| hIL1B RT RV |
| hIL32 FW |

| |
|---|
| hIL32 RV |
| hIL6 RT FW |
| hIL6 RT RV |
| hIL8 FW RT |
| hIL8 RV RT |
| hIRF2 RT FW |
| hIRF2 RT RV |
| hIRF3 FW |
| hIRF3 RV |
| hIRF5 FW |
| hIRF5 RV |
| hIRF7 FW |
| hIRF7 RV |
| hIRF9 FW |
| hIRF9 RV |
| hMAVS rt fw |
| hMAVS rt rv |
| hOAS1 RT FW |
| hOAS1 RT RV |
| hIGFBP7 FW RT |
| hIGFBP7 RV RT |
| mOas1 fw |
| mOas1 rv |
| m and h IRF7 FW |
| m and h IRF7 RV |
| mActb fw rt |
| mActb rv rt |
| mB2m fw |
| mB2m rv |
| mCdkn1a FW RT |
| mCdkn1a RV RT |
| mCdkn2a FW RT |
| mCdkn2a RV RT |
| CHERRY RT FW |
| CHERRY RT RV |
| mCxcl15 fw |
| mCxcl15 rv |
| mDdx58 fw |
| mDdx58 rv |
| mHprt fw |
| mHprt rv |
| mIfih1 fw |
| mIfih1 rv |
| mIgfbp7 fw |
| mIgfbp7 rv |
| mIl1b fw |
| mIl1b rv |
| mLINC9 RV |
| mLINC9 RV |
| mMMTV FW |
| mMMTV RV |

| |
|---|
| mMULV FW |
| mMULV RV |
| mMUSD FW |
| mMUSD RV |

| Sequence (5'->3') | Application |
|---|---|
| AAAGGATCCATGGTGAGCAAGGGCGAG | Cloning |
| AAAGAATTCCTACCCGGTAGAATTATCTAGATC | Cloning |
| AAA CTC GAG CTA CCC GGT AGA ATT ATC | Cloning |
| AAAGAATTCATGGGGAGCAGCAAGAGCAAG | Cloning |
| AAAGAATTCGGCCGTGCCGCTGGCCGAGT | Cloning |
| AAAAGCTAGCATGGGGAGCAGCAAGAGCAAG | Cloning |
| AAATCTAGATCAGGCCGTGCCGCTG | Cloning |
| AAAGTCGACGGCCGTGCCGCTGGCC | Cloning |
| AAAAGGATCCTCAGGCCGTGCCGCTG | Cloning |
| GAGGATCCATGTACCCATACGATGTTCC | Cloning |
| GAGAATTCTTAGCTGGCCTCCACCTTTC | Cloning |
| AAAGAATTCGACCTTCCGCTTCTTCTTTG | Cloning |
| aaaGCTAGCATGAGACATATTATCTGCCA | Cloning |
| aaaGGATCCttatggcctggggcgtttac | Cloning |
| AAAACGCGTATGAGACATATTATCTGCCACG | Cloning |
| AAAAGATCTTTATTCAGACACAGGACCCTCT | Cloning |
| AAAGTCGACTTATTCAGACACAGGACC | Cloning |
| AAAACGCGTATGGTGAGCAAGGGCGAGG | Cloning |
| AAAAGATCTCTTGTACAGCTCGTCCATGCCG | Cloning |
| CATCACGAATTCAATGTCCGAGACTGCTCCT | Cloning |
| TACTACAGATCTAATTTCTTCTTGGGCGCCGC | Cloning |
| CATGAATTCATGAGCTCCCAAAGCCATCCA | Cloning |
| CATGAATTCCAGGGGCGGCTCCTCTTCCAT | Cloning |
| AAAACGCGTATGAGCTCCCAAAGCCATCCAG | Cloning |
| AAACATGAATTCCTACGTACTCGTGTTGGGGAAGC | Cloning |
| AAACATGTCGACCGTACTCGTGTTGGGGAAGC | Cloning |
| AAAGCTAGCATGACGGAATATAAGCTGGT | Cloning |
| AAAGGATCCTCAGGAGAGCACACACTTG | Cloning |
| AAAGGATCCATGACGGAATATAAGCTG | Cloning |
| AAACTCGAGGGAGAGCACACACTTGCA | Cloning |
| AAAGGATCCATGGCGACTGCGACC | Cloning |
| AAAGTCGACCATAATTGCACAGCTTCTATTG | Cloning |
| AAAAGATCTATGGATTACAAGGACGATGACAAGACCGGTCCGTTTGCTGAAGAC | Cloning |
| AAAACCGGTATGCCGTTTGCTGAAGACAAGACCTATAGAGCTACCAGCCTCGGACCT | Cloning |
| AAAGTCGACCTAGTGCAGACGCCGCCGGTAC | Cloning |
| AAA AGA TCT ATG CCG TTT GCT GAA GAC AAG | Cloning |
| AAA GTC GAC GTG CAG ACG CCG CCG GT | Cloning |
| AAACCATGGATGACCGAGTACAAGCCCACG | Cloning |
| AAAGAGCTCTCAGGCACCGGGCTTGCG | Cloning |
| AAAGGATCCATGGACTATAAGGACCACG | Cloning |
| AAATACGTAATGGATTACAAGGACGACGATGACAAG | Cloning |
| AAACTCGAGTCATTTGGACATTTCTGCTGGATC | Cloning |
| CACCGGCAGGATTCAGCAGCTCCAC | Crispr/Cas9 |
| AAACGTGGAGCTGCTGAATCCTGCC | Crispr/Cas9 |
| CACCGCGTGAACCACATGCCCAGCA | Crispr/Cas9 |
| AAACTGCTGGGCATGTGGTTCACGC | Crispr/Cas9 |
| CACCGGATTCAGCAGCTCCACTGGC | Crispr/Cas9 |
| AAACGCCAGTGGAGCTGCTGAATCC | Crispr/Cas9 |
| CACCGTTCTGGCCGAGACCAGCCAG | Crispr/Cas9 |
| AAACCTGGCTGGTCTCGGCCAGAAC | Crispr/Cas9 |

| | |
|---|---|
| CTGGCCGAGACCAGGCAGTGGAGCTGCTG | Mutagenesis |
| CAGCAGCTCCACTGCCTGGTCTCGGCCAG | Mutagenesis |
| GGAGCTGCTGAATGCTGCCCGCGTGAAC | Mutagenesis |
| GTTCACGCGGGCAGCATTCAGCAGCTCC | Mutagenesis |
| CACATGCCCAGCACCCTGCATGTGGCCAC | Mutagenesis |
| GTGGCCACATCCAGGGTGCTGGGATGTG | Mutagenesis |

| Sequence (5'->3') | Application |
|---|---|
| CGC CGC CAG CTC ACC ATG | qPCR |
| CAC GAT GGA GGG GAA GAC GG | qPCR |
| AGTCAGTTCCTTGTGGAGCC | qPCR |
| CATGGGTTCTGACGGACAT | qPCR |
| GTTACGGTCGGAGGCCG | qPCR |
| GTGAGAGTGGCGGGGTC | qPCR |
| TCCTGAACCTGAGTAGAGACAC | qPCR |
| TGCTGCTTGTAGTGGCTGG | qPCR |
| TGTGCTCCTACAGGTTGTGGA | qPCR |
| CACTGGGATCTGATTCGCAAAA | qPCR |
| CTTGGAGTCCTCACTCAAACTC | qPCR |
| ACTGCTGCAACTACCCTTAAACA | qPCR |
| ATATCCCTCACCACGATCCTAATA | qPCR |
| CCCTCTGTAGTGCAAAGACTGATA | qPCR |
| TCCAATCCCACCATCTGCTT | qPCR |
| TGTCAGGCGATAGGTGTGTT | qPCR |
| TGGCCCAAGATTCCATTCCT | qPCR |
| TTGCCAAAATGTTACCGGGG | qPCR |
| ATGCCACTCAGGGATGTGA | qPCR |
| TTCTCAACTGCCATTCTGGA | qPCR |
| AAGAGCGTGCCCTACTTCAA | qPCR |
| GCACACGAATGAACTTGCTG | qPCR |
| CTCTTGGAGACCGACGCTG | qPCR |
| GCAGGATCCTTCCATTGAGA | qPCR |
| CCCTTCATTGACCTCAACTACATG | qPCR |
| TGGGATTTCCATTGATGACAAGC | qPCR |
| CAAGAACAAGGAGAAGGGCAAAG | qPCR |
| GGAGAACTCTGGTCAAGGGAACTG | qPCR |
| ACTTCTCTGCACAGCATCCC | qPCR |
| AGTGTGGGGTCCACAGAGC | qPCR |
| CTCACTGTCAGCCCCAGAG | qPCR |
| CTGGTGCTTCAGCATGACC | qPCR |
| AGTAGAGAGGCATCGCAGAGA | qPCR |
| GGAGTGTCTTTCGTTGCTGAT | qPCR |
| AGACTTTGCTTTCCTTGGTCAGG | qPCR |
| GTCTGGCTTATATCCAACACTTCG | qPCR |
| CCATCTATCAGCAGGCTCCG | qPCR |
| TTTCTTACCTGCCTCCACCC | qPCR |
| ACAGAAACCGGATTGCTGCTG | qPCR |
| GCTCTTCCACTTGAGGACCA | qPCR |
| GAAGCTGATGGCCCTAAACA | qPCR |
| AAGCCCTTGCTGTAGTGGTG | qPCR |
| CCTGAGCAGAAGTAGGGAGG | qPCR |

| | |
|---|---|
| TTGGCTCCTTGAACTTTTGG | qPCR |
| ACTCACCTCTTCAGAACGAATTG | qPCR |
| CCATCTTTGGAAGGTTCAGGTTG | qPCR |
| CGGAAGGAACCATCTCACTG | qPCR |
| AGCACTCCTTGGCAAAACTG | qPCR |
| CATGCGGCTAGACATGGGTG | qPCR |
| GCTTTCCTGTATGGATTGCCC | qPCR |
| ACC AGC CGT GGA CCA AGA G | qPCR |
| TAC CAA GGCCCT GAG GCA C | qPCR |
| AAG CCG ATC CGG CCA A | qPCR |
| GGA AGT CCC GGC TCT TGT TAA | qPCR |
| TGG TCC TGG TGA AGC TGG AA | qPCR |
| GAT GTC GTC ATA GAG GCT GTT GG | qPCR |
| CAT GGC TCT CTT CCC AGA AA | qPCR |
| AGC TCT TCA GAA CCG CCT AC | qPCR |
| CAGGCCGAGCCTATCATCTG | qPCR |
| GGGCTTTGAGCTAGTTGGCA | qPCR |
| AGCTTCGTACTGAGTTCGCTC | qPCR |
| CCAGTCAACTGACCCAGGG | qPCR |
| GCTCAAGTACACCTGGGCAC | qPCR |
| CATCACCCAGGTCAGCAAG | qPCR |
| GGGCCTCTAAAGGGGTCAAG | qPCR |
| TCAAACTTCACTCCACAACGTC | qPCR |
| GTG ACC ATC ATG TAC AAG GG | qPCR |
| TAC ACC TTG CAC TTG CCC | qPCR |
| GGCTGTATTCCCCTCCATCG | qPCR |
| CCAGTTGGTAACAATGCCATGT | qPCR |
| GTCTCACTGACCGGCCTGTATG | qPCR |
| CCCGTTCTTCAGCATTTGGATTTC | qPCR |
| CAGATCCACAGCGATATCCA | qPCR |
| ACGGGACCGAAGAGACAAC | qPCR |
| CGTGAACATGTTGTTGAGGC | qPCR |
| GCAGAAGAGCTGCTACGTGA | qPCR |
| GCCCCGTAATGCAGAAGAAG | qPCR |
| GTGTAGTCCTCGTTGTGGGA | qPCR |
| TCGAGACCATTTACTGCAACAG | qPCR |
| CATTGCCGGTGGAAATTCCTT | qPCR |
| CAGATCCGAGACACTAAAGGGA | qPCR |
| TCCTCATCAGCCTTGCTTTCA | qPCR |
| GTGTTGGATACAGGCCAGACTTTG | qPCR |
| ACTAACAGGACTCCTCGTATTTGC | qPCR |
| AGATCAACACCTGTGGTAACACC | qPCR |
| CTCTAGGGCCTCCACGAACA | qPCR |
| AAGAGGCGGAAGGGTAAAGC | qPCR |
| TGGGGTAGGTGATGCCGTT | qPCR |
| GAAATGCCACCTTTTGACAGTG | qPCR |
| TGGATGCTCTCATCAGGACAG | qPCR |
| TGA GTG GAA CAC AAC TTC TGC | qPCR |
| CAG GCA AGC TCT CTT CTT GC | qPCR |
| TTT CCC GAA GAA GGA GGA TT | qPCR |
| GCT TCT GCG GAT AGC AAA AC | qPCR |

| | |
|---|---|
| TTC TGC TCC TCT TCT GCC CT | qPCR |
| GAG GAC CCT GGG CAA GAA AC | qPCR |
| ATA GAG GCC GCT TCT TTG C | qPCR |
| TGA GAC TCC ACC AAA TGT CC | qPCR |

| Gene | FC (KO/wt) | pvalue adj |
|---|---|---|
| LCE2A | 23.29058749 | 0.000163804 |
| LCP1 | 19.29060605 | 0 |
| IL1B | 17.08732118 | 0.000163804 |
| CSF2 | 15.1394832 | 0.000271569 |
| MARCH11 | 10.73534703 | 0.003969089 |
| IGFBP7 | 9.854857439 | 0.000163804 |
| SCG5 | 9.493261285 | 0.000644977 |
| EDN1 | 8.908144906 | 0.002540217 |
| NCKAP1L | 8.742933397 | 0 |
| IL1A | 7.81650163 | 0.004432832 |
| CCDC148 | 7.810287639 | 0.000163804 |
| PSG4 | 7.582220796 | 0.000163804 |
| HECW2 | 7.209763827 | 0.000271569 |
| BAMBI | 6.954256872 | 0 |
| CLDN1 | 6.873085446 | 0.000163804 |
| PTPRR | 6.817666506 | 0 |
| IL32 | 6.65098877 | 0.000163804 |
| CXCL10 | 6.415040262 | 0.03268545 |
| IAPP | 6.37450829 | 0.030140371 |
| AKR1C3 | 5.983583326 | 0.003969089 |
| ALPL | 5.960806154 | 0.009105558 |
| DNER | 5.941435077 | 0.031556402 |
| LUM | 5.59931778 | 0.000163804 |
| MMP9 | 5.551522905 | 0.001483869 |
| ALPK2 | 5.356094407 | 0.016023797 |
| DDIT4L | 4.963889714 | 0.0440915 |
| BIRC3 | 4.859787642 | 0.000163804 |
| IGFBP3 | 4.498924563 | 0.007082643 |
| FBXO32 | 4.268287381 | 0.000687976 |
| LOC284998 | 4.230555815 | 0.027853199 |
| LOC653580 | 4.188209519 | 0.000163804 |
| PAPPA | 4.174054821 | 0.000163804 |
| MMP3 | 4.039767838 | 0.011055248 |
| VCAN | 3.973934996 | 0.009157191 |
| HS.311428 | 3.867323513 | 0.0197883 |
| GBP4 | 3.820847144 | 0.039664476 |
| PLA2G4C | 3.78649461 | 0.001464272 |
| WNT5B | 3.77165086 | 0.004452283 |
| HS.374023 | 3.715652481 | 0 |
| MAMDC2 | 3.660380149 | 0.001154644 |
| LOX | 3.613384685 | 0.000469074 |
| PDGFRL | 3.55909218 | 0.016411397 |
| FAP | 3.50438738 | 0.024453821 |
| CXCR4 | 3.431881447 | 0.01095796 |
| PSG1 | 3.421113098 | 0 |
| FOLR3 | 3.405574756 | 0.046549929 |
| ARHGDIB | 3.379249376 | 0.043186842 |
| UBD | 3.378391161 | 0.007882332 |
| TGM2 | 3.376949134 | 0.000163804 |
| SNORD13 | 3.364731434 | 0.001547945 |

| | | |
|---|---|---|
| MPP4 | 3.278102644 | 0.000163804 |
| SLCO1B3 | 3.258823741 | 0.000271569 |
| LOC100133984 | 3.231799959 | 0.004452283 |
| RFPL4A | 3.177469325 | 0.042431606 |
| C6ORF58 | 3.127744755 | 0.000163804 |
| HS.390407 | 3.125262463 | 0.009217308 |
| MTSS1 | 3.120248859 | 0.00687033 |
| CXCL2 | 3.10550618 | 0.000163804 |
| SNCAIP | 3.080961007 | 0.001502225 |
| PID1 | 3.057399791 | 0.005143436 |
| KRT34 | 3.056483251 | 0.000163804 |
| SNORA12 | 3.040341699 | 0.000687976 |
| FLJ14213 | 3.039984218 | 0.002109538 |
| PSG3 | 3.036343116 | 0.003309249 |
| CXCL1 | 3.023549711 | 0.001799936 |
| LOC441061 | 2.994777901 | 0.040491223 |
| LOC728855 | 2.99140399 | 0.004432832 |
| PSG6 | 2.986587774 | 0.000986435 |
| IFIH1 | 2.98024314 | 0.022570277 |
| IL8 | 2.977538585 | 0.001799936 |
| RNF182 | 2.962563616 | 0.000644977 |
| ICAM2 | 2.96091052 | 0.000271569 |
| FAM176A | 2.944037933 | 0 |
| UCA1 | 2.899856724 | 0.000377548 |
| AKR1C4 | 2.855659063 | 0.012913742 |
| NRIP1 | 2.825893298 | 0.006334529 |
| IRX3 | 2.825179795 | 0.009157191 |
| ABI3BP | 2.820120237 | 0.003309249 |
| MST4 | 2.810759814 | 0.000687976 |
| HBE1 | 2.748505443 | 0.015200982 |
| SOD2 | 2.744330724 | 0.002163794 |
| ASB5 | 2.7257994 | 0.009463519 |
| POU3F2 | 2.703358801 | 0.003309249 |
| DCN | 2.701904158 | 0.005061221 |
| DRAM1 | 2.701448137 | 0.001272283 |
| TMEM132A | 2.685853291 | 0.004432832 |
| TNFAIP3 | 2.682628749 | 0.003613879 |
| GBP5 | 2.682197601 | 0.001483869 |
| MSC | 2.674571542 | 0.003309249 |
| TMEM166 | 2.669436607 | 0.002630495 |
| CD83 | 2.665085874 | 0.000163804 |
| LOC100133866 | 2.641408903 | 0.014451031 |
| LOC389300 | 2.639674562 | 0.015049464 |
| NGF | 2.62890541 | 0.007237502 |
| LOC644173 | 2.610703235 | 0.027306319 |
| ESM1 | 2.605895462 | 0.000163804 |
| CFLAR | 2.581306702 | 0.005597619 |
| AKR1C2 | 2.573502288 | 0.020546055 |
| RNF150 | 2.522479831 | 0.030348206 |
| NLGN1 | 2.501904148 | 0.005609169 |
| HLA-B | 2.47017215 | 0.000163804 |

| | | |
|---|---|---|
| PRDM1 | 2.465525976 | 0.027853199 |
| SOX4 | 2.461232996 | 0.019593542 |
| PSG2 | 2.4387809 | 0.000687976 |
| DEFB103B | 2.438714125 | 0.004788849 |
| LOC644350 | 2.416217417 | 0.015963902 |
| CTHRC1 | 2.390721962 | 0.008291645 |
| BCL11A | 2.386125511 | 0.000163804 |
| LOC388681 | 2.383772798 | 0.008322284 |
| HS.21177 | 2.359148113 | 0.000163804 |
| IFI6 | 2.357883823 | 0.039835999 |
| LOC642477 | 2.354510518 | 0.011629742 |
| ZEB2 | 2.341290594 | 0.000986435 |
| ITGAV | 2.333066224 | 0.009002232 |
| STC2 | 2.32539743 | 0.001483869 |
| C21ORF7 | 2.320107443 | 0.002827976 |
| LOC653506 | 2.316713422 | 0.007292189 |
| ERN1 | 2.311428114 | 0.003354942 |
| ZNF697 | 2.287985081 | 0.002827976 |
| JAM2 | 2.275411275 | 0.027254413 |
| ZSWIM4 | 2.27443395 | 0.001838555 |
| LOC440160 | 2.260101354 | 0.000644977 |
| LOC651876 | 2.252090015 | 0.001636039 |
| KRT222 | 2.241691797 | 0.017090027 |
| DSG2 | 2.237115842 | 0.000163804 |
| KCNG1 | 2.223162445 | 0.002131228 |
| WWC1 | 2.223056653 | 0.003309249 |
| M160 | 2.194555006 | 0.000687976 |
| ETS2 | 2.185091141 | 0.022822263 |
| DLC1 | 2.182426298 | 0.022957175 |
| LOC650369 | 2.170334591 | 0.000163804 |
| NYNRIN | 2.143256065 | 0.000687976 |
| F2RL2 | 2.14067147 | 0.000687976 |
| DNAJC12 | 2.140099554 | 0.000163804 |
| SPANXD | 2.138412678 | 0.012670215 |
| FAM43B | 2.134042654 | 0.011550602 |
| MARCH4 | 2.111644627 | 0.008383109 |
| XYLT1 | 2.10690494 | 0.019996797 |
| BDKRB1 | 2.102228194 | 0.000163804 |
| BCAT1 | 2.090885493 | 0.001039387 |
| MYO1D | 2.090158019 | 0.002250558 |
| PSG7 | 2.034087543 | 0.008658423 |
| DIRAS3 | -2.041008713 | 0.002554364 |
| RELN | -2.045825247 | 0.000163804 |
| LOC730413 | -2.048805951 | 0.003354942 |
| TSPAN8 | -2.06548224 | 0.024453821 |
| INSIG1 | -2.083184931 | 0.000163804 |
| ENPP1 | -2.089466828 | 0.001838555 |
| CA12 | -2.102596022 | 0.016411397 |
| EDIL3 | -2.121716539 | 0.005609169 |
| ATOH8 | -2.131502105 | 0.000687976 |
| LOC646723 | -2.136775127 | 0.000163804 |

| | | |
|---|---|---|
| BAIAP2L1 | -2.188181908 | 0.029074988 |
| ABLIM1 | -2.203364564 | 0.014960916 |
| C14ORF23 | -2.218228669 | 0.007609426 |
| CA9 | -2.218235361 | 0.007065193 |
| HOXA11AS | -2.221130345 | 0.015455071 |
| TMEM233 | -2.22266379 | 0.012473635 |
| NTN4 | -2.223691628 | 0.00607576 |
| DDX10 | -2.239992346 | 0.000271569 |
| LOC100130111 | -2.241744445 | 0.000687976 |
| MAP7 | -2.247770493 | 0.000737117 |
| PEX11G | -2.276578819 | 0.006935688 |
| MBP | -2.288419757 | 0.005597619 |
| CD96 | -2.307182937 | 0.019969158 |
| RGS17 | -2.314953362 | 0.000271569 |
| CYSLTR1 | -2.320283601 | 0.000377548 |
| SEMA3A | -2.323300175 | 0.009661897 |
| SPTLC3 | -2.362900707 | 0.000163804 |
| HGF | -2.368708194 | 0.000687976 |
| ADORA2B | -2.378276997 | 0.010052169 |
| EPHB6 | -2.378358953 | 0.007850199 |
| ZNF385D | -2.378758213 | 0.000271569 |
| EVL | -2.392745383 | 0.009078356 |
| TIMP3 | -2.396609321 | 0.000377548 |
| F2RL1 | -2.443624561 | 0.015438691 |
| KRT19 | -2.481038536 | 0.042088403 |
| KAT2B | -2.482944263 | 0.002827976 |
| HS.176498 | -2.491807466 | 0.002554364 |
| CMBL | -2.523765098 | 0.017159905 |
| AGPAT9 | -2.553455647 | 0.000687976 |
| SNORD114-1 | -2.561204995 | 0.000644977 |
| ALDH2 | -2.567805996 | 0.039305018 |
| HS.370359 | -2.582443669 | 0.027504397 |
| JMJD8 | -2.602721287 | 0.041471969 |
| DKFZP761P0423 | -2.616585967 | 0.003354942 |
| FOXA2 | -2.690432208 | 0.003309249 |
| MAL2 | -2.694331093 | 0.03089747 |
| AK5 | -2.753252357 | 0 |
| APCDD1L | -2.789409802 | 0.045571811 |
| BLK | -2.794882373 | 0.014665652 |
| ARL14 | -2.833024064 | 0.00931341 |
| SMAD6 | -2.835500594 | 0.000469074 |
| GKN2 | -2.837378762 | 0.000687976 |
| FST | -2.907585695 | 0.012670215 |
| DOCK2 | -2.964034243 | 0.000163804 |
| MGC42105 | -2.985092829 | 0.000163804 |
| PPAP2B | -3.024453563 | 0.000163804 |
| RGS7 | -3.029634384 | 0.000579755 |
| NR2F1 | -3.039188383 | 0.00457891 |
| FCGRT | -3.067689402 | 0.003613879 |
| DHRS9 | -3.072163227 | 0.032926414 |
| C10ORF116 | -3.201844181 | 0.00342578 |

| | | |
|---|---|---|
| PRAGMIN | -3.204748572 | 0.003495359 |
| KIAA0363 | -3.251316515 | 0.000163804 |
| DENND2A | -3.310639653 | 0.000687976 |
| PAMR1 | -3.325295557 | 0.000163804 |
| CNTNAP2 | -3.349659985 | 0.010993567 |
| PRICKLE1 | -3.356169376 | 0.000469074 |
| SPANXC | -3.426079002 | 0.003309249 |
| LAMA5 | -3.630810322 | 0.013945936 |
| MALL | -3.680008558 | 0.003338705 |
| HSPB6 | -3.759687957 | 0.002827976 |
| FAM5C | -3.93910614 | 0.000163804 |
| CPA4 | -4.161306244 | 0.000986435 |
| NCRNA00161 | -4.277910249 | 0.00687033 |
| C21ORF100 | -4.599085644 | 0.026576433 |
| CSAG1 | -4.626035172 | 0.028939152 |
| SYTL2 | -4.839914876 | 0.000469074 |
| SLC14A1 | -4.969218678 | 0.001592536 |
| FAM107A | -5.28235475 | 0.009108408 |
| C20ORF103 | -5.529810076 | 0.000163804 |
| COX7A1 | -6.390701111 | 0.028200504 |
| HS.554324 | -6.395202215 | 0.000271569 |
| PPARG | -7.349895803 | 0.000163804 |
| PTGER2 | -7.406641179 | 0.000163804 |
| MEG3 | -10.07070058 | 0.001502225 |
| SFRP1 | -13.5620957 | 0.000377548 |
| C5ORF46 | -23.7454918 | 0 |
| ANXA10 | -27.84183349 | 0 |

| NFKB signature | SASP signature | RIS up-regulated genes |
|---|---|---|
| A4 | ANG | BPGM |
| ABCA1 | AREG | C6orf192 |
| ABCB1 | AXL | CA12 |
| ABCB4 | BMP2 | CEBPD |
| ABCB9 | BMP6 | CXCL6 |
| ABCC6 | CCL1 | FOLR3 |
| ABCG5 | CCL13 | HEY1 |
| ABCG8 | CCL16 | HIST2H2AC |
| ADAM19 | CCL2 | HSD17B14 |
| ADH1A | CCL26 | IL1RAPL1 |
| ADORA1 | CCL3 | INSIG1 |
| ADORA2A | CCL7 | JUP |
| ADRA2B | CCL8 | KCNJ15 |
| AFP | CD55 | LOC283050 |
| AGER | CD9 | MGC87042 |
| AGT | CSF2 | MMP10 |
| AHCTF1 | CSF2RB | MT1X |
| AICDA | CXCL1 | NTSR1 |
| AKR1C1 | CXCL5 | NUDT14 |
| ALOX12 | EGFR | OTUB2 |
| ALOX5 | ENA78 | RAB31 |
| AMACR | EREG | SERPINA9 |
| AMH | ETS2 | SERPIND1 |
| ANGPT1 | FAS | SLC25A37 |
| APOBEC2 | FGF2 | STEAP1 |
| APOC3 | FGF7 | STX3 |
| APOD | GCP2 | TNFAIP3 |
| APOE | GDF15 | TRAF3IP2 |
| AQP4 | GEM | TRIB1 |
| AR | GMFG | TSC22D1 |
| ARFRP1 | HGF | WNT7B |
| ART1 | ICAM1 | LOC654103 |
| ASPH | ICAM3 | CTSK |
| ASS1 | IGF1 | SLC20A1 |
| ATP1A2 | IGF2R | LPXN |
| B2M | IGFBP1 | PDE4B |
| BACE1 | IGFBP2 | CST4 |
| BAX | IGFBP6 | CXCL10 |
| BCL2 | IGFBP7 | RAC2 |
| BCL2A1 | IL13 | ODC1 |
| BCL2L1 | IL15 | NINJ1 |
| BCL2L11 | IL1A | SLC6A15 |
| BCL3 | IL1B | ABCA1 |
| BDKRB1 | IL6 | SQRDL |
| BDKRB1 | IL7 | PDPN |
| BDNF | INHBA | SLC11A2 |
| BGN | IQGAP2 | TMEM132A |
| BLIMP1 /PRDM1 | ITGA2 | DUSP6 |
| BLNK | ITPKA | SLC2A6 |
| BLR1 | JUN | GPR68 |

| | | |
|---|---|---|
| BMI1 | MIF | SP140 |
| BMP2 | MMP1 | AKR1B1 |
| BMP4 | MMP10 | GK |
| BNIP3 | MMP2 | CCL3 |
| BRCA2 | MMP3 | CCL3L3 |
| BTK | NAP2 | OAF |
| C3 | NRG1 | LCE2A |
| C4A | PECAM1 | CYGB |
| C4BPA | PIGF | MX1 |
| C69 | PLAUR | LOC647650 |
| CALCB | PTGES | KYNU |
| CASP4 | RPS6KA5 | HIST2H2AA3 |
| CAV1 | SERIPINE1 | MME |
| CCL1 | TGFB1 | IRAK2 |
| CCL15 | TIMP2 | ANGPTL4 |
| CCL17 | TNFRSF11B | NAMPT |
| CCL19 | VEGFA | BMP2 |
| CCL2 | VEGFC | IL11 |
| CCL20 | WNT2 | DIO2 |
| CCL20 | | GFPT2 |
| CCL22 | | C8orf4 |
| CCL23 | | NRG1 |
| CCL28 | | ACP5 |
| CCL3 | | COL10A1 |
| CCL4 | | M160 |
| CCL4 | | TAGLN3 |
| CCL5 | | CSF3 |
| CCL5 | | SERPINB4 |
| CCND1 | | ZC3H12A |
| CCND2 | | QPCT |
| CCND3 | | IL24 |
| CCR5 | | NFKBIZ |
| CCR7 | | SAT1 |
| CD209 | | LOC387763 |
| CD274 | | TFPI2 |
| CD38 | | IER3 |
| CD3G | | SOD2 |
| CD3G | | PAPPA |
| CD40 | | LIF |
| CD40LG | | MMP1 |
| CD44 | | IL8 |
| CD48 | | CCL20 |
| CD54 | | MMP3 |
| CD80 | | NDP |
| CD83 | | CXCL2 |
| CD86 | | CXCL5 |
| CDK6 | | FOXQ1 |
| CDKN1A | | GDF15 |
| CDX1 | | C15orf48 |
| CEBPD | | CXCL1 |
| CFB | | SLC16A6 |

CFB
CFLAR
CGM3
CHI3L1
CIDEA
CLDN2
COL1A2
CR2
CR2
CREB3
CRP
CSF1
CSF2
CSF3
CTSB
CTSL1
CXCL1
CXCL1
CXCL1
CXCL1
CXCL10
CXCL10
CXCL11
CXCL3
CXCL3
CXCL5
CXCL5
CXCL5
CXCL6
CXCL9
CYP19A1
CYP27B1
CYP2C11
CYP2E1
CYP7B1
DCTN4
DEFB2
DIO2
DMP1
DNASE1L2
DPYD
DUSP1
E2F3
EBI3
EDN1
EGFR
EGR1
ELF3
ENG
ENO2
EPHA1
CYP26B1
SCG5
SERPINB2
IL1A
PTGS2
CST1
IL6
CSF2
IL1B

EPO
ERBB2
ERVWE1
F11R
F3
F8
FABP6
FAM148A
FAS
FASLG
FCER2
FCER2/CD23
FCGRT
FGF8
FN1
FOS
FSTL3
FTH1
FTH1
G6PC
G6PD
GAD1
GADD45B
GATA3
GBP1
GCLC
GCLC
GCLC
GCLC
GCLM
GCLM
GCNT1
GJB1
GNAI2
GNB2L1
GNRH2
GRIN1
GRIN2A
GRM2
GSTP1
GUCY1A2
GZMB
HAMP
HAS1
HBE1
HBZ
HGF
HIF1A
HLA-B
HLA-G
HMGN1

HMOX1
HOXA9
HPSE
HSD11B2
HSD17B8
HSP90AA1
ICOS
ICOS
IDO1
IER2
IER3
IER3
IER3
IFI44L
IFNB1
IFNG
IGFBP1
IGFBP2
IGFBP2
IGHE
IGHG1
IGHG2
IGHG4
IGKC
Iigp1
IL10
IL11
IL12A
IL12B
IL13
IL15
IL17
IL1A
IL1B
IL1RN
IL2
IL23A
IL27
IL2RA
IL6
IL8
IL8RA
IL8RB
IL9
INHBA
IRF1
IRF2
IRF4
IRF7
JMJD3
JUNB

KC
KCNK5
KCNN2
KISS1
KITLG
KLF10
KLK3
KLRA1
KRT15
KRT3
KRT5
KRT6B
LAMB2
LBP
LCN2
LCN2
LEF1
LGALS3
LIPG
LTA
LTA
LTB
LTF
LYZ
MADCAM1
MAP4K1
MBP
MDK
MMP1
MMP3
MMP9
MMP9
MT3
MTHFR
MUC2
MX1
MYB
MYC
MYLK
MYOZ1
NCAM
NFKB1
NFKB2
NFKBIA
NFKBIE
NFKBIZ
NGFB
NK4
NLRP2
NOD2
NOS1

NOS2A
NOS2A
NOX1
NPY1R
NQO1
NR3C1
NR4A2
NRG1
NUAK2
OLR1
OPN1SW
OPRD1
OPRM1
ORM1
Osterix
OXTR
PAFAH2
PAX8
PDE7A
PDGFB
PDYN
PENK
PGK1
PGLYRP1
PGR
PI3KAP1
PIGF
pIgR
PIK3CA
PIM1
PLA2
PLAU
PLCD1
PLK3
POMC
PPARGC1B
PPP5C
PRF1
PRKACA
PRKCD
PRL
PSMB9
PSME1
PSME2
PTAFR
PTEN
PTGDS
PTGES
PTGIS
PTGS2
PTHLH

PTPN1
PTPN13
PTS
PTX3
PYCARD
RAG1
RAG2
RBBP4
Rdh1
Rdh7
REL
RELB
REV3L
RIPK2
S100A10
S100A4
S100A6
SAA1
SAA2
SAA3
SAT1
SCNN1A
SDC4
SELE
SELP
SELS
SENP2
SERPINA1
SERPINA2
SERPINA3
SERPINB1
SERPINE1, PAI-1
SERPINE2
SH3BGRL
SKALP, PI3
SKP2
SLC11A2
SLC16A1
SLC3A2
SLC6A6
Slfn2
SNAI1
SOD1
SOD1
SOD2
SOX9
SPATA19
SPI1
SPP1
ST6GAL1
ST8SIA1

STAT5A
SUPV3L1
TACR1
TAP1
TAPBP
TCRB
TERT
TF
TFEC
TFF3
TFPI2
TGM1
TGM2
THBS1
THBS2
TICAM1
TIFA
TLR2
TLR9
TNC
TNF
TNFAIP2
TNFAIP3
TNFRSF1B
TNFRSF4
TNFRSF9
TNFSF10
TNFSF13B
TNFSF13B
TNFSF15
TNIP1
TNIP3
TP53
TRAF1
TRAF2
TREM1
TRPC1
TWIST1
UBE2M
UCP2
UGCGL1
UPK1B
UPP1
VCAM1
VEGFC
VIM
VPS53
WNT10B
WT1
XDH
XIAP

| |
|---|
| YY1 |
| ZNF366 |

| Chromosome (hg38) | Start | End |
|---|---|---|
| chr1 | 7247506 | 7.27E+06 |
| chr1 | 7993530 | 8.06E+06 |
| chr1 | 8076808 | 8.10E+06 |
| chr1 | 9160898 | 9.21E+06 |
| chr1 | 15158118 | 1.52E+07 |
| chr1 | 23095593 | 2.31E+07 |
| chr1 | 25227080 | 2.53E+07 |
| chr1 | 28109459 | 2.81E+07 |
| chr1 | 31470205 | 3.15E+07 |
| chr1 | 31927522 | 31956944 |
| chr1 | 36343094 | 36389051 |
| chr1 | 39146603 | 39206641 |
| chr1 | 40362532 | 40405060 |
| chr1 | 40867712 | 40879254 |
| chr1 | 41595151 | 41684323 |
| chr1 | 58747035 | 58766595 |
| chr1 | 64879384 | 64936362 |
| chr1 | 66206678 | 66295518 |
| chr1 | 66521079 | 66599937 |
| chr1 | 67679528 | 67771276 |
| chr1 | 83132960 | 83165842 |
| chr1 | 85272887 | 85345071 |
| chr1 | 87220438 | 87237516 |
| chr1 | 91554597 | 91586250 |
| chr1 | 93661535 | 93703518 |
| chr1 | 94636324 | 94676712 |
| chr1 | 94707455 | 94802320 |
| chr1 | 98075209 | 98091786 |
| chr1 | 99586589 | 99622550 |
| chr1 | 108813842 | 108832835 |
| chr1 | 112722377 | 112744683 |
| chr1 | 114453406 | 114474449 |
| chr1 | 115411206 | 115433269 |
| chr1 | 119631225 | 119649470 |
| chr1 | 120035243 | 120154589 |
| chr1 | 144976275 | 145018530 |
| chr1 | 144935189 | 144948002 |
| chr1 | 149149496 | 149163764 |
| chr1 | 148915380 | 148998666 |
| chr1 | 148840319 | 148895802 |
| chr1 | 148818251 | 148821390 |
| chr1 | 146194624 | 148797646 |
| chr1 | 147026631 | 147057241 |
| chr1 | 150602978 | 150621453 |
| chr1 | 153552172 | 153574334 |
| chr1 | 156095813 | 156130630 |
| chr1 | 161111000 | 161120566 |
| chr1 | 163071848 | 163091631 |
| chr1 | 165893343 | 165914274 |
| chr1 | 167773445 | 167824815 |

| | | |
|---|---|---|
| chr1 | 170519093 | 170563931 |
| chr1 | 170660723 | 170713341 |
| chr1 | 171737735 | 171755124 |
| chr1 | 172897260 | 172955605 |
| chr1 | 178122026 | 178146059 |
| chr1 | 179127946 | 179147349 |
| chr1 | 183020956 | 183046283 |
| chr1 | 186182407 | 186192801 |
| chr1 | 186937650 | 186985650 |
| chr1 | 192517349 | 192574928 |
| chr1 | 198098448 | 198129885 |
| chr1 | 198771907 | 198795891 |
| chr1 | 198878813 | 198938507 |
| chr1 | 199222931 | 199230230 |
| chr1 | 199270416 | 199298206 |
| chr1 | 201500673 | 201565124 |
| chr1 | 203642850 | 203676007 |
| chr1 | 206350433 | 206363533 |
| chr1 | 206873314 | 206906414 |
| chr1 | 207705397 | 207709049 |
| chr1 | 209872743 | 209902174 |
| chr1 | 214394041 | 214473135 |
| chr1 | 218459646 | 218500410 |
| chr1 | 219770039 | 219800927 |
| chr1 | 221429339 | 221459422 |
| chr1 | 221707859 | 221767037 |
| chr1 | 221793049 | 221888209 |
| chr1 | 221981935 | 222002864 |
| chr1 | 222060448 | 222082715 |
| chr1 | 223697947 | 223749903 |
| chr1 | 223801975 | 223844321 |
| chr1 | 224047657 | 224092427 |
| chr1 | 224488544 | 224582015 |
| chr1 | 234494279 | 234546143 |
| chr1 | 234599411 | 234632316 |
| chr1 | 234955082 | 235026533 |
| chr1 | 235900753 | 235924156 |
| chr1 | 240230575 | 240259666 |
| chr1 | 240332629 | 240357308 |
| chr1 | 243250859 | 243296091 |
| chr1 | 243617520 | 243626577 |
| chr1 | 245082029 | 245094532 |
| chr1 | 246565652 | 246633301 |
| chr10 | 575265 | 627794 |
| chr10 | 3738432 | 3788802 |
| chr10 | 3801743 | 3907960 |
| chr10 | 4239988 | 4270462 |
| chr10 | 4646830 | 4707204 |
| chr10 | 4762288 | 4776049 |
| chr10 | 13681664 | 13725005 |
| chr10 | 13860072 | 13895056 |

| | | |
|---|---|---|
| chr10 | 16944300 | 17090791 |
| chr10 | 17199133 | 17232621 |
| chr10 | 22596796 | 22633334 |
| chr10 | 24428982 | 24468134 |
| chr10 | 30964209 | 31008420 |
| chr10 | 31739703 | 31769750 |
| chr10 | 32938356 | 33015870 |
| chr10 | 33128653 | 33158549 |
| chr10 | 33237501 | 33307576 |
| chr10 | 35889445 | 35916588 |
| chr10 | 36489430 | 36500444 |
| chr10 | 36524015 | 36542926 |
| chr10 | 43128938 | 43167490 |
| chr10 | 48588303 | 48635137 |
| chr10 | 49159198 | 49181906 |
| chr10 | 61872918 | 61906008 |
| chr10 | 62024939 | 62084607 |
| chr10 | 62731897 | 62746764 |
| chr10 | 63263367 | 63271366 |
| chr10 | 63697620 | 63720145 |
| chr10 | 68070642 | 68107306 |
| chr10 | 68133682 | 68203202 |
| chr10 | 71831288 | 71868790 |
| chr10 | 73883424 | 73917433 |
| chr10 | 77156206 | 77166362 |
| chr10 | 77348079 | 77361562 |
| chr10 | 77578100 | 77630239 |
| chr10 | 78253551 | 78268684 |
| chr10 | 78970119 | 79001346 |
| chr10 | 79111017 | 79160682 |
| chr10 | 79317749 | 79338496 |
| chr10 | 87816060 | 87849283 |
| chr10 | 88149519 | 88159989 |
| chr10 | 91335514 | 91368310 |
| chr10 | 91587135 | 91617914 |
| chr10 | 93435069 | 93480663 |
| chr10 | 95246170 | 95280177 |
| chr10 | 99657921 | 99695436 |
| chr10 | 103730287 | 103793569 |
| chr10 | 104092829 | 104122057 |
| chr10 | 104246303 | 104264488 |
| chr10 | 104291084 | 104352615 |
| chr10 | 110350223 | 110427646 |
| chr10 | 110486566 | 110540091 |
| chr10 | 126040925 | 126065798 |
| chr10 | 126241858 | 126268985 |
| chr11 | 9138150 | 9176826 |
| chr11 | 9564525 | 9578396 |
| chr11 | 10350482 | 10408362 |
| chr11 | 12030167 | 12048955 |
| chr11 | 12064183 | 12090893 |

| | | |
|---|---|---|
| chr11 | 12110374 | 12243832 |
| chr11 | 15715174 | 15755043 |
| chr11 | 19711641 | 19739966 |
| chr11 | 22453413 | 22468482 |
| chr11 | 26217908 | 26252464 |
| chr11 | 27903088 | 27937974 |
| chr11 | 29013862 | 29026756 |
| chr11 | 31642861 | 31655629 |
| chr11 | 33508154 | 33558100 |
| chr11 | 33669415 | 33734489 |
| chr11 | 34625329 | 34657728 |
| chr11 | 34677915 | 34686799 |
| chr11 | 35028984 | 35088901 |
| chr11 | 35138314 | 35184373 |
| chr11 | 35321428 | 35392693 |
| chr11 | 35538813 | 35556214 |
| chr11 | 43784033 | 43791749 |
| chr11 | 43893588 | 43944665 |
| chr11 | 44567037 | 44593103 |
| chr11 | 58335008 | 58340974 |
| chr11 | 58571483 | 58583912 |
| chr11 | 65275424 | 65344032 |
| chr11 | 65415645 | 65427589 |
| chr11 | 65470336 | 65510821 |
| chr11 | 69296017 | 69322333 |
| chr11 | 69326789 | 69368733 |
| chr11 | 74924806 | 74943623 |
| chr11 | 86624412 | 86644539 |
| chr11 | 89496264 | 89500016 |
| chr11 | 94127790 | 94153859 |
| chr11 | 96146126 | 96169327 |
| chr11 | 96199198 | 96263505 |
| chr11 | 96302166 | 96340351 |
| chr11 | 102138164 | 102198431 |
| chr11 | 102342684 | 102350349 |
| chr11 | 102577638 | 102604740 |
| chr11 | 102749565 | 102803689 |
| chr11 | 102824688 | 102853733 |
| chr11 | 102981004 | 102998449 |
| chr11 | 107583270 | 107590284 |
| chr11 | 109195092 | 109198376 |
| chr11 | 110068165 | 110095933 |
| chr11 | 112502020 | 112505290 |
| chr11 | 115905577 | 115944266 |
| chr11 | 122135467 | 122200422 |
| chr11 | 123165513 | 123240759 |
| chr11 | 127620029 | 127657464 |
| chr11 | 127915542 | 127926608 |
| chr11 | 128447938 | 128531187 |
| chr11 | 128573783 | 128605140 |
| chr11 | 131743771 | 131774387 |

| | | |
|---|---|---|
| chr11 | 132025437 | 132077740 |
| chr12 | 11697175 | 11732901 |
| chr12 | 12425481 | 12457446 |
| chr12 | 13097469 | 13104259 |
| chr12 | 13165085 | 13225015 |
| chr12 | 14325615 | 14329373 |
| chr12 | 24708216 | 24722193 |
| chr12 | 24897226 | 24949665 |
| chr12 | 26266987 | 26278017 |
| chr12 | 27570020 | 27583920 |
| chr12 | 28021315 | 28037282 |
| chr12 | 28972435 | 28975840 |
| chr12 | 29732066 | 29746109 |
| chr12 | 31731471 | 31766764 |
| chr12 | 43276785 | 43311866 |
| chr12 | 45215841 | 45236176 |
| chr12 | 46429104 | 46497212 |
| chr12 | 46546253 | 46564085 |
| chr12 | 46657952 | 46668532 |
| chr12 | 47503922 | 47510487 |
| chr12 | 51882513 | 51906212 |
| chr12 | 57117891 | 57171697 |
| chr12 | 58499673 | 58511957 |
| chr12 | 58526556 | 58537523 |
| chr12 | 64157526 | 64190501 |
| chr12 | 64612802 | 64634018 |
| chr12 | 65522917 | 65539918 |
| chr12 | 65601014 | 65700152 |
| chr12 | 65818359 | 65902898 |
| chr12 | 65922769 | 65959529 |
| chr12 | 68784718 | 68828841 |
| chr12 | 75622322 | 75726502 |
| chr12 | 75739128 | 75878420 |
| chr12 | 75917476 | 75987193 |
| chr12 | 76010779 | 76038109 |
| chr12 | 77086159 | 77127109 |
| chr12 | 77937067 | 77950160 |
| chr12 | 88296711 | 88340953 |
| chr12 | 88888565 | 88905892 |
| chr12 | 89051403 | 89083574 |
| chr12 | 89154740 | 89194246 |
| chr12 | 89209019 | 89254766 |
| chr12 | 89369499 | 89394094 |
| chr12 | 89416864 | 89478134 |
| chr12 | 89703925 | 89736120 |
| chr12 | 90250406 | 90284232 |
| chr12 | 92538660 | 92606435 |
| chr12 | 93761626 | 93805851 |
| chr12 | 94168347 | 94177922 |
| chr12 | 96189871 | 96222323 |
| chr12 | 96392539 | 96454807 |

| | | |
|---|---|---|
| chr12 | 105912786 | 105924715 |
| chr12 | 107026313 | 107031363 |
| chr12 | 108716312 | 108735458 |
| chr12 | 108800925 | 108858077 |
| chr12 | 111393521 | 111443871 |
| chr12 | 114656329 | 114692866 |
| chr12 | 115146226 | 115158385 |
| chr12 | 115310918 | 115355587 |
| chr12 | 120221807 | 120247927 |
| chr12 | 124903508 | 124940784 |
| chr12 | 127229550 | 127286601 |
| chr13 | 23652728 | 23680292 |
| chr13 | 33168346 | 33208601 |
| chr13 | 33227854 | 33286637 |
| chr13 | 33646891 | 33650173 |
| chr13 | 37303302 | 37320082 |
| chr13 | 37403403 | 37454272 |
| chr13 | 37473333 | 37483237 |
| chr13 | 41587813 | 41649953 |
| chr13 | 42803364 | 42837343 |
| chr13 | 44220342 | 44318954 |
| chr13 | 50730144 | 50734848 |
| chr13 | 64538076 | 64556655 |
| chr13 | 76004922 | 76016139 |
| chr13 | 77858776 | 77879826 |
| chr13 | 79932689 | 79944492 |
| chr13 | 79984388 | 79999635 |
| chr13 | 80030322 | 80057516 |
| chr13 | 87085137 | 87132143 |
| chr13 | 93845998 | 93868122 |
| chr13 | 94232266 | 94241937 |
| chr13 | 97254357 | 97328802 |
| chr13 | 100874011 | 100902829 |
| chr13 | 101516616 | 101536802 |
| chr13 | 106367061 | 106381218 |
| chr13 | 107096697 | 107104830 |
| chr13 | 107184273 | 107217755 |
| chr13 | 107246444 | 107308317 |
| chr13 | 107346859 | 107378040 |
| chr13 | 107421336 | 107458824 |
| chr13 | 108943774 | 108995081 |
| chr13 | 110300847 | 110361716 |
| chr13 | 110379356 | 110428231 |
| chr13 | 110938587 | 110967170 |
| chr13 | 114098519 | 114133118 |
| chr14 | 22822747 | 22853873 |
| chr14 | 23207625 | 23213163 |
| chr14 | 31926140 | 31957095 |
| chr14 | 32341681 | 32374247 |
| chr14 | 35154937 | 35163637 |
| chr14 | 35362318 | 35420457 |

| | | |
|---|---|---|
| chr14 | 51384687 | 51401891 |
| chr14 | 51439051 | 51475424 |
| chr14 | 51532361 | 51582239 |
| chr14 | 52788573 | 52814692 |
| chr14 | 52894175 | 52920763 |
| chr14 | 54588349 | 54688098 |
| chr14 | 55365492 | 55419601 |
| chr14 | 61460764 | 61478766 |
| chr14 | 61523631 | 61611008 |
| chr14 | 61990573 | 62002837 |
| chr14 | 62015422 | 62020050 |
| chr14 | 68124012 | 68149200 |
| chr14 | 68774999 | 68816930 |
| chr14 | 68931227 | 68988971 |
| chr14 | 69036407 | 69081218 |
| chr14 | 70881390 | 70911276 |
| chr14 | 72634812 | 72702711 |
| chr14 | 73754126 | 73790659 |
| chr14 | 76191304 | 76206341 |
| chr14 | 88971569 | 89009367 |
| chr14 | 91222455 | 91249568 |
| chr14 | 96084939 | 96146891 |
| chr14 | 96159861 | 96218665 |
| chr14 | 96237885 | 96293705 |
| chr14 | 99743828 | 99780640 |
| chr14 | 100978568 | 100998797 |
| chr15 | 32681599 | 32731074 |
| chr15 | 32818438 | 32880648 |
| chr15 | 33079566 | 33127737 |
| chr15 | 34807880 | 34816812 |
| chr15 | 39118794 | 39156293 |
| chr15 | 39198462 | 39317508 |
| chr15 | 39453187 | 39479887 |
| chr15 | 48670600 | 48724969 |
| chr15 | 49490197 | 49507432 |
| chr15 | 51982270 | 52022714 |
| chr15 | 59165137 | 59201332 |
| chr15 | 59292601 | 59307644 |
| chr15 | 60346769 | 60422448 |
| chr15 | 62098675 | 62139298 |
| chr15 | 62886352 | 62897913 |
| chr15 | 63466312 | 63516493 |
| chr15 | 65652951 | 65668334 |
| chr15 | 67064808 | 67152710 |
| chr15 | 68581398 | 68645585 |
| chr15 | 70758472 | 70807407 |
| chr15 | 71042476 | 71094588 |
| chr15 | 74381216 | 74430937 |
| chr15 | 74503752 | 74543215 |
| chr15 | 79942837 | 79979825 |
| chr15 | 85401319 | 85421704 |

| | | |
|---|---|---|
| chr15 | 85583776 | 85626024 |
| chr15 | 88621211 | 88650548 |
| chr15 | 89083355 | 89123384 |
| chr15 | 90399104 | 90425868 |
| chr15 | 91536599 | 91564357 |
| chr15 | 101167504 | 101209327 |
| chr16 | 9046906 | 9111558 |
| chr16 | 15012281 | 15029567 |
| chr16 | 20864542 | 20914089 |
| chr16 | 21757166 | 21771907 |
| chr16 | 21786571 | 21795451 |
| chr16 | 22573306 | 22588078 |
| chr16 | 22602695 | 22611409 |
| chr16 | 27221752 | 27274696 |
| chr16 | 29250862 | 29350088 |
| chr16 | 51853527 | 51902669 |
| chr16 | 56603583 | 56623798 |
| chr16 | 70011239 | 70028469 |
| chr16 | 70802256 | 70808478 |
| chr16 | 82181972 | 82198177 |
| chr16 | 82628672 | 82674781 |
| chr16 | 83527632 | 83554586 |
| chr16 | 86341323 | 86405957 |
| chr16 | 87431473 | 87468943 |
| chr16 | 88223494 | 88275193 |
| chr16 | 89300238 | 89336073 |
| chr16 | 89408596 | 89511274 |
| chr17 | 2156733 | 2198997 |
| chr17 | 2211654 | 2250538 |
| chr17 | 5825247 | 5837369 |
| chr17 | 7833946 | 7845768 |
| chr17 | 8914387 | 8949406 |
| chr17 | 13337885 | 13366483 |
| chr17 | 13538383 | 13564370 |
| chr17 | 13577111 | 13602491 |
| chr17 | 21280030 | 21306168 |
| chr17 | 30241850 | 30277531 |
| chr17 | 40095885 | 40124604 |
| chr17 | 40900895 | 40923980 |
| chr17 | 40947997 | 40960781 |
| chr17 | 47222478 | 47280298 |
| chr17 | 59753060 | 59788491 |
| chr17 | 59804647 | 59859993 |
| chr17 | 60081651 | 60120556 |
| chr17 | 61284109 | 61340397 |
| chr17 | 63273700 | 63294079 |
| chr17 | 64647068 | 64750479 |
| chr17 | 65154196 | 65208687 |
| chr17 | 67412052 | 67444589 |
| chr17 | 67474929 | 67535566 |
| chr17 | 68286688 | 68321691 |

| | | |
|---|---|---|
| chr17 | 68374929 | 68401438 |
| chr17 | 71399400 | 71443818 |
| chr17 | 72386333 | 72433048 |
| chr17 | 78309203 | 78360186 |
| chr17 | 78374058 | 78393073 |
| chr17 | 80850346 | 80861888 |
| chr18 | 3246790 | 3306967 |
| chr18 | 3445763 | 3479557 |
| chr18 | 3578708 | 3635062 |
| chr18 | 9704426 | 9753889 |
| chr18 | 22464400 | 22472391 |
| chr18 | 23562299 | 23595698 |
| chr18 | 29148059 | 29202556 |
| chr18 | 39943506 | 39951234 |
| chr18 | 48719076 | 48735863 |
| chr18 | 48922001 | 48999302 |
| chr18 | 49046883 | 49058334 |
| chr18 | 54732702 | 54745719 |
| chr18 | 55499118 | 55517297 |
| chr18 | 58204937 | 58242674 |
| chr18 | 59872447 | 59918017 |
| chr18 | 63736017 | 63790587 |
| chr18 | 63866875 | 63890548 |
| chr18 | 67755293 | 67785628 |
| chr18 | 68192270 | 68195496 |
| chr18 | 68416747 | 68455786 |
| chr18 | 68548903 | 68564451 |
| chr18 | 68836665 | 68846003 |
| chr18 | 73679867 | 73719564 |
| chr18 | 73923271 | 73926645 |
| chr18 | 73943915 | 73981750 |
| chr19 | 11088165 | 11097892 |
| chr19 | 13615494 | 13641202 |
| chr19 | 13837964 | 13862679 |
| chr19 | 18359774 | 18389068 |
| chr19 | 31212464 | 31234171 |
| chr19 | 37996241 | 38006437 |
| chr19 | 38650119 | 38694318 |
| chr19 | 42108302 | 42130434 |
| chr19 | 42188869 | 42219163 |
| chr19 | 45421819 | 45485883 |
| chr2 | 5625405 | 5637791 |
| chr2 | 5765423 | 5805310 |
| chr2 | 6823028 | 6835493 |
| chr2 | 9166149 | 9252338 |
| chr2 | 10571328 | 10599931 |
| chr2 | 24244977 | 24272140 |
| chr2 | 24702120 | 24743542 |
| chr2 | 33209104 | 33244604 |
| chr2 | 33353189 | 33374576 |
| chr2 | 37373883 | 37396080 |

| | | |
|---|---|---|
| chr2 | 37560692 | 37589484 |
| chr2 | 37623001 | 37689364 |
| chr2 | 39479939 | 39514568 |
| chr2 | 39989814 | 40043078 |
| chr2 | 46305278 | 46337022 |
| chr2 | 54524472 | 54589600 |
| chr2 | 62570872 | 62587759 |
| chr2 | 65354724 | 65383904 |
| chr2 | 66838781 | 66898098 |
| chr2 | 67254632 | 67312510 |
| chr2 | 67499381 | 67527483 |
| chr2 | 67558594 | 67579588 |
| chr2 | 69162020 | 69261517 |
| chr2 | 72569939 | 72578611 |
| chr2 | 87444796 | 87513380 |
| chr2 | 95433638 | 95445426 |
| chr2 | 95631309 | 95643100 |
| chr2 | 99839911 | 99883995 |
| chr2 | 102226214 | 102259998 |
| chr2 | 108571028 | 108596274 |
| chr2 | 109207872 | 109243679 |
| chr2 | 111437283 | 111505724 |
| chr2 | 112782109 | 112842369 |
| chr2 | 112865827 | 112888743 |
| chr2 | 113884614 | 113919118 |
| chr2 | 127000089 | 127035657 |
| chr2 | 142857468 | 142881448 |
| chr2 | 144079097 | 144141583 |
| chr2 | 145591601 | 145603009 |
| chr2 | 145617426 | 145656984 |
| chr2 | 145738603 | 145762565 |
| chr2 | 146309608 | 146325834 |
| chr2 | 149731762 | 149747580 |
| chr2 | 150108710 | 150147831 |
| chr2 | 150471821 | 150691778 |
| chr2 | 157150944 | 157190355 |
| chr2 | 160269345 | 160317963 |
| chr2 | 160375923 | 160434969 |
| chr2 | 162070164 | 162119131 |
| chr2 | 162194210 | 162262590 |
| chr2 | 167132838 | 167145332 |
| chr2 | 173116472 | 173191817 |
| chr2 | 174743588 | 174808436 |
| chr2 | 179453263 | 179468954 |
| chr2 | 180507818 | 180526272 |
| chr2 | 180703614 | 180761496 |
| chr2 | 181774001 | 181815425 |
| chr2 | 181947657 | 181979861 |
| chr2 | 187542102 | 187555304 |
| chr2 | 189253028 | 189307543 |
| chr2 | 191199947 | 191250071 |

| | | |
|---|---|---|
| chr2 | 191611799 | 191696789 |
| chr2 | 195520906 | 195577578 |
| chr2 | 196231820 | 196280887 |
| chr2 | 196658183 | 196663538 |
| chr2 | 200396084 | 200456264 |
| chr2 | 201116166 | 201132685 |
| chr2 | 202686659 | 202696561 |
| chr2 | 203673900 | 203697244 |
| chr2 | 205649749 | 205734969 |
| chr2 | 207220293 | 207261833 |
| chr2 | 207309326 | 207343541 |
| chr2 | 212839662 | 212858685 |
| chr2 | 215651476 | 215666365 |
| chr2 | 215680339 | 215774846 |
| chr2 | 216297205 | 216320585 |
| chr2 | 217047089 | 217107776 |
| chr2 | 217204815 | 217240755 |
| chr2 | 225089778 | 225120375 |
| chr2 | 225988517 | 226004829 |
| chr2 | 226048202 | 226082708 |
| chr2 | 230227107 | 230244969 |
| chr2 | 230748885 | 230799651 |
| chr2 | 234239780 | 234262598 |
| chr2 | 237113162 | 237236623 |
| chr2 | 237253557 | 237316748 |
| chr2 | 237400836 | 237504237 |
| chr2 | 238423185 | 238441711 |
| chr20 | 1376542 | 1436600 |
| chr20 | 4533176 | 4575049 |
| chr20 | 6465085 | 6474668 |
| chr20 | 6492270 | 6535328 |
| chr20 | 7713082 | 7745742 |
| chr20 | 7771583 | 7776074 |
| chr20 | 7824494 | 7892190 |
| chr20 | 10510626 | 10543004 |
| chr20 | 17541542 | 17581597 |
| chr20 | 19810406 | 19833155 |
| chr20 | 23139194 | 23167750 |
| chr20 | 31693149 | 31722301 |
| chr20 | 35302254 | 35329694 |
| chr20 | 38123105 | 38174106 |
| chr20 | 41048371 | 41080832 |
| chr20 | 44486300 | 44526386 |
| chr20 | 44571942 | 44610659 |
| chr20 | 47306552 | 47361644 |
| chr20 | 50303840 | 50349926 |
| chr20 | 50415143 | 50564715 |
| chr20 | 51325634 | 51401661 |
| chr20 | 51433153 | 51489884 |
| chr20 | 53735678 | 53807473 |
| chr20 | 53826070 | 53831533 |

| | | |
|---|---|---|
| chr20 | 53864957 | 53961676 |
| chr21 | 27564705 | 27583666 |
| chr21 | 27644292 | 27652813 |
| chr21 | 28292903 | 28318694 |
| chr21 | 29160541 | 29225923 |
| chr21 | 33360321 | 33385473 |
| chr21 | 34790710 | 34819534 |
| chr21 | 34834579 | 34891858 |
| chr21 | 34965081 | 35005113 |
| chr21 | 35159251 | 35266492 |
| chr21 | 38243687 | 38330183 |
| chr21 | 38804617 | 38860648 |
| chr21 | 38984753 | 39021977 |
| chr21 | 41993419 | 42058308 |
| chr21 | 43492694 | 43523652 |
| chr21 | 46013415 | 46144181 |
| chr22 | 20508728 | 20533963 |
| chr22 | 30177944 | 30263656 |
| chr22 | 30408832 | 30459712 |
| chr22 | 36324561 | 36399854 |
| chr22 | 37302359 | 37330669 |
| chr22 | 38201902 | 38242299 |
| chr22 | 38298161 | 38334442 |
| chr3 | 4410015 | 4428689 |
| chr3 | 4976313 | 5027223 |
| chr3 | 5294610 | 5332216 |
| chr3 | 10169947 | 10223839 |
| chr3 | 11272198 | 11307007 |
| chr3 | 15630098 | 15648874 |
| chr3 | 15767843 | 15808124 |
| chr3 | 16052147 | 16076435 |
| chr3 | 16089548 | 16148880 |
| chr3 | 23648240 | 23670608 |
| chr3 | 24363741 | 24367264 |
| chr3 | 27519527 | 27562597 |
| chr3 | 30329130 | 30357545 |
| chr3 | 30538005 | 30555096 |
| chr3 | 39139988 | 39180208 |
| chr3 | 43708192 | 43771277 |
| chr3 | 45040999 | 45181057 |
| chr3 | 46087449 | 46127717 |
| chr3 | 55142981 | 55206091 |
| chr3 | 58987699 | 59001504 |
| chr3 | 62585430 | 62616061 |
| chr3 | 66397877 | 66443165 |
| chr3 | 67076852 | 67095161 |
| chr3 | 67781731 | 67784771 |
| chr3 | 70430531 | 70435310 |
| chr3 | 71048788 | 71130839 |
| chr3 | 81723931 | 81765630 |
| chr3 | 98777635 | 98825023 |

| | | |
|---|---|---|
| chr3 | 98885389 | 98906813 |
| chr3 | 98953580 | 98987706 |
| chr3 | 99105815 | 99132212 |
| chr3 | 99619823 | 99679378 |
| chr3 | 99858749 | 99903350 |
| chr3 | 100029780 | 100076676 |
| chr3 | 101926354 | 101965178 |
| chr3 | 105996785 | 106031403 |
| chr3 | 108128293 | 108134606 |
| chr3 | 115217417 | 115249124 |
| chr3 | 115766516 | 115807487 |
| chr3 | 115986915 | 116020958 |
| chr3 | 119630022 | 119645196 |
| chr3 | 124832160 | 124862203 |
| chr3 | 127722900 | 127823603 |
| chr3 | 129485522 | 129504973 |
| chr3 | 129649501 | 129661961 |
| chr3 | 132248492 | 132260861 |
| chr3 | 132619727 | 132638712 |
| chr3 | 133270513 | 133304551 |
| chr3 | 134077138 | 134104133 |
| chr3 | 141329053 | 141427072 |
| chr3 | 143123044 | 143179087 |
| chr3 | 143434998 | 143446804 |
| chr3 | 149331179 | 149345434 |
| chr3 | 149361546 | 149402420 |
| chr3 | 149569290 | 149621794 |
| chr3 | 152256454 | 152332969 |
| chr3 | 153273692 | 153294577 |
| chr3 | 155062470 | 155087140 |
| chr3 | 156532746 | 156556201 |
| chr3 | 156664612 | 156733790 |
| chr3 | 156749415 | 156837732 |
| chr3 | 157064610 | 157101163 |
| chr3 | 158701298 | 158750806 |
| chr3 | 170699271 | 170728831 |
| chr3 | 171780186 | 171835911 |
| chr3 | 172124635 | 172187100 |
| chr3 | 172622418 | 172686166 |
| chr3 | 176662562 | 176665661 |
| chr3 | 177960520 | 177995944 |
| chr3 | 183265529 | 183307722 |
| chr3 | 188260809 | 188300245 |
| chr3 | 190000149 | 190078535 |
| chr3 | 190172299 | 190235337 |
| chr3 | 190301500 | 190347815 |
| chr3 | 191325184 | 191340327 |
| chr3 | 194548654 | 194599603 |
| chr3 | 195149843 | 195199461 |
| chr4 | 5709877 | 5787971 |
| chr4 | 7806993 | 7859278 |

| | | |
|---|---|---|
| chr4 | 13890419 | 13933507 |
| chr4 | 23943495 | 23959493 |
| chr4 | 26014722 | 26089796 |
| chr4 | 26170853 | 26199787 |
| chr4 | 28367794 | 28387860 |
| chr4 | 38119349 | 38164636 |
| chr4 | 38940181 | 38996460 |
| chr4 | 40977605 | 41010494 |
| chr4 | 46424178 | 46447536 |
| chr4 | 48322964 | 48344009 |
| chr4 | 54058677 | 54087005 |
| chr4 | 71124663 | 71134944 |
| chr4 | 71286688 | 71305633 |
| chr4 | 73683458 | 73742875 |
| chr4 | 74005021 | 74058435 |
| chr4 | 74096938 | 74118511 |
| chr4 | 74391364 | 74427287 |
| chr4 | 74500274 | 74550676 |
| chr4 | 74883415 | 74888762 |
| chr4 | 76723042 | 76784689 |
| chr4 | 76976585 | 77019183 |
| chr4 | 78746861 | 78787946 |
| chr4 | 79998069 | 80079240 |
| chr4 | 80124497 | 80156481 |
| chr4 | 80230667 | 80272562 |
| chr4 | 85759592 | 85796987 |
| chr4 | 88520563 | 88538026 |
| chr4 | 102786246 | 102830320 |
| chr4 | 113465709 | 113478833 |
| chr4 | 113945584 | 113998827 |
| chr4 | 119093242 | 119115868 |
| chr4 | 121666865 | 121712403 |
| chr4 | 122748179 | 122810972 |
| chr4 | 123823775 | 123839386 |
| chr4 | 126140943 | 126146329 |
| chr4 | 130478714 | 130487714 |
| chr4 | 138536840 | 138541747 |
| chr4 | 140036752 | 140134393 |
| chr4 | 140251790 | 140268985 |
| chr4 | 145277606 | 145309634 |
| chr4 | 147312554 | 147373184 |
| chr4 | 147542433 | 147591269 |
| chr4 | 148748531 | 148795829 |
| chr4 | 156750014 | 156783322 |
| chr4 | 156929541 | 157005530 |
| chr4 | 158156023 | 158162437 |
| chr4 | 158774543 | 158825029 |
| chr4 | 168432312 | 168453560 |
| chr4 | 176421060 | 176446037 |
| chr4 | 176532225 | 176550828 |
| chr4 | 176726609 | 176806326 |

| | | |
|---|---|---|
| chr4 | 176899736 | 176928619 |
| chr4 | 176979571 | 177017840 |
| chr4 | 177124967 | 177129518 |
| chr4 | 177479104 | 177493715 |
| chr4 | 186900658 | 186937175 |
| chr4 | 188399091 | 188452152 |
| chr5 | 365938 | 425108 |
| chr5 | 8825319 | 8852081 |
| chr5 | 9033137 | 9070219 |
| chr5 | 9146250 | 9213384 |
| chr5 | 9431203 | 9469053 |
| chr5 | 14033151 | 14063202 |
| chr5 | 14142170 | 14212032 |
| chr5 | 14399420 | 14478780 |
| chr5 | 17105198 | 17191266 |
| chr5 | 30500600 | 30507411 |
| chr5 | 33772036 | 33819082 |
| chr5 | 33831749 | 33856672 |
| chr5 | 34561150 | 34612141 |
| chr5 | 35115622 | 35139204 |
| chr5 | 37696220 | 37739570 |
| chr5 | 38824565 | 38857351 |
| chr5 | 39092721 | 39122552 |
| chr5 | 39391483 | 39448168 |
| chr5 | 39695521 | 39699257 |
| chr5 | 39749168 | 39784779 |
| chr5 | 40219823 | 40239205 |
| chr5 | 42983867 | 43009246 |
| chr5 | 52982948 | 53072348 |
| chr5 | 53321756 | 53431400 |
| chr5 | 58230872 | 58275404 |
| chr5 | 59742892 | 59762034 |
| chr5 | 61274995 | 61334718 |
| chr5 | 65028180 | 65038164 |
| chr5 | 65101048 | 65128992 |
| chr5 | 65162999 | 65214153 |
| chr5 | 67003137 | 67039962 |
| chr5 | 67298726 | 67316087 |
| chr5 | 75592015 | 75612135 |
| chr5 | 76787284 | 76839103 |
| chr5 | 78484002 | 78552011 |
| chr5 | 78585551 | 78606468 |
| chr5 | 78889191 | 78908907 |
| chr5 | 87112493 | 87156917 |
| chr5 | 92524958 | 92563353 |
| chr5 | 96014085 | 96093397 |
| chr5 | 96682624 | 96716529 |
| chr5 | 96878070 | 96904763 |
| chr5 | 98094205 | 98112564 |
| chr5 | 98294620 | 98316491 |
| chr5 | 98452684 | 98510614 |

| | | |
|---|---|---|
| chr5 | 106375186 | 106404352 |
| chr5 | 109821419 | 109845625 |
| chr5 | 109862454 | 109879373 |
| chr5 | 110621208 | 110629147 |
| chr5 | 110649313 | 110654241 |
| chr5 | 111061069 | 111080057 |
| chr5 | 111983115 | 112013123 |
| chr5 | 113221207 | 113267363 |
| chr5 | 114313226 | 114334274 |
| chr5 | 115413600 | 115467978 |
| chr5 | 120124261 | 120162489 |
| chr5 | 120364408 | 120407348 |
| chr5 | 122074827 | 122105104 |
| chr5 | 122141626 | 122183865 |
| chr5 | 125039553 | 125060212 |
| chr5 | 132064796 | 132111075 |
| chr5 | 132421298 | 132461012 |
| chr5 | 135979149 | 136022701 |
| chr5 | 140292283 | 140339224 |
| chr5 | 142318933 | 142366449 |
| chr5 | 142443812 | 142486721 |
| chr5 | 142998505 | 143019446 |
| chr5 | 143234694 | 143264486 |
| chr5 | 143306112 | 143323359 |
| chr5 | 143535587 | 143597009 |
| chr5 | 145478720 | 145509868 |
| chr5 | 148964783 | 149016639 |
| chr5 | 149130972 | 149229794 |
| chr5 | 149999521 | 150068037 |
| chr5 | 151062437 | 151109263 |
| chr5 | 151658880 | 151701131 |
| chr5 | 151739304 | 151751010 |
| chr5 | 155928702 | 155933533 |
| chr5 | 157540438 | 157612230 |
| chr5 | 159158145 | 159186780 |
| chr5 | 172353398 | 172365202 |
| chr5 | 172756954 | 172956794 |
| chr5 | 173449587 | 173478385 |
| chr6 | 1890353 | 1922290 |
| chr6 | 1948979 | 1974287 |
| chr6 | 4344852 | 4384706 |
| chr6 | 4636683 | 4669413 |
| chr6 | 11745049 | 11774230 |
| chr6 | 12174414 | 12211113 |
| chr6 | 14658533 | 14682933 |
| chr6 | 14695661 | 14770743 |
| chr6 | 16316598 | 16352592 |
| chr6 | 16674416 | 16690777 |
| chr6 | 16704211 | 16769963 |
| chr6 | 17904460 | 17998673 |
| chr6 | 30740978 | 30787312 |

| | | |
|---|---|---|
| chr6 | 36643044 | 36682983 |
| chr6 | 37238104 | 37260501 |
| chr6 | 42190200 | 42227661 |
| chr6 | 44034003 | 44074893 |
| chr6 | 45433522 | 45484527 |
| chr6 | 52491220 | 52551580 |
| chr6 | 56516746 | 56543915 |
| chr6 | 56669361 | 56731452 |
| chr6 | 63789437 | 63809178 |
| chr6 | 64221036 | 64262960 |
| chr6 | 71382898 | 71420818 |
| chr6 | 73694486 | 73736433 |
| chr6 | 74247883 | 74270728 |
| chr6 | 75266274 | 75284936 |
| chr6 | 85400394 | 85418496 |
| chr6 | 85447670 | 85484032 |
| chr6 | 92930971 | 92950316 |
| chr6 | 99834722 | 99837761 |
| chr6 | 105151517 | 105214313 |
| chr6 | 105544962 | 105618613 |
| chr6 | 105726506 | 105820002 |
| chr6 | 106061386 | 106103530 |
| chr6 | 111227018 | 111241162 |
| chr6 | 111551534 | 111606587 |
| chr6 | 111970872 | 112049198 |
| chr6 | 112201785 | 112252589 |
| chr6 | 113628480 | 113646543 |
| chr6 | 116373914 | 116426273 |
| chr6 | 117485232 | 117506286 |
| chr6 | 121502758 | 121529914 |
| chr6 | 127850071 | 127856112 |
| chr6 | 128486961 | 128541165 |
| chr6 | 129488316 | 129501517 |
| chr6 | 129671617 | 129703787 |
| chr6 | 130586869 | 130609696 |
| chr6 | 132682206 | 132692826 |
| chr6 | 133237123 | 133257370 |
| chr6 | 134419404 | 134449578 |
| chr6 | 136582005 | 136614845 |
| chr6 | 137687154 | 137776474 |
| chr6 | 137804588 | 137878339 |
| chr6 | 138943183 | 138987706 |
| chr6 | 139289612 | 139336362 |
| chr6 | 139530709 | 139547017 |
| chr6 | 139966350 | 139985455 |
| chr6 | 140032357 | 140097778 |
| chr6 | 141539032 | 141563182 |
| chr6 | 142819030 | 142906927 |
| chr6 | 150992312 | 151072007 |
| chr6 | 152294771 | 152320670 |
| chr6 | 157755438 | 157813513 |

| | | |
|---|---|---|
| chr6 | 158002915 | 158071580 |
| chr6 | 159679453 | 159732491 |
| chr6 | 161026895 | 161050016 |
| chr7 | 2795843 | 2828375 |
| chr7 | 12016588 | 12068917 |
| chr7 | 17520726 | 17565775 |
| chr7 | 18502661 | 18522606 |
| chr7 | 18848894 | 18869096 |
| chr7 | 20217051 | 20231413 |
| chr7 | 22560645 | 22612749 |
| chr7 | 24776814 | 24789468 |
| chr7 | 36650776 | 36686535 |
| chr7 | 39568291 | 39628355 |
| chr7 | 41026597 | 41054856 |
| chr7 | 41095261 | 41116259 |
| chr7 | 41143488 | 41203479 |
| chr7 | 41218662 | 41256820 |
| chr7 | 41686087 | 41710945 |
| chr7 | 41867525 | 41894451 |
| chr7 | 41909890 | 41943255 |
| chr7 | 43868833 | 43885353 |
| chr7 | 46834308 | 46862093 |
| chr7 | 47449779 | 47484232 |
| chr7 | 50791435 | 50852152 |
| chr7 | 55048544 | 55142289 |
| chr7 | 74061695 | 74100860 |
| chr7 | 76568937 | 76614509 |
| chr7 | 77072594 | 77084544 |
| chr7 | 77407648 | 77464572 |
| chr7 | 78195466 | 78219308 |
| chr7 | 79974084 | 79989720 |
| chr7 | 80486986 | 80501936 |
| chr7 | 81584081 | 81656858 |
| chr7 | 81673926 | 81729884 |
| chr7 | 84171715 | 84266095 |
| chr7 | 84368650 | 84386397 |
| chr7 | 90401363 | 90414232 |
| chr7 | 92621052 | 92646227 |
| chr7 | 92716517 | 92770304 |
| chr7 | 94042826 | 94069781 |
| chr7 | 94129977 | 94162476 |
| chr7 | 101094218 | 101140198 |
| chr7 | 102786810 | 102798769 |
| chr7 | 102859108 | 102898806 |
| chr7 | 104943326 | 104989740 |
| chr7 | 106343419 | 106350038 |
| chr7 | 111423688 | 111431316 |
| chr7 | 115340310 | 115365301 |
| chr7 | 116310235 | 116336174 |
| chr7 | 116500820 | 116589182 |
| chr7 | 117123260 | 117144664 |

| | | |
|---|---|---|
| chr7 | 120062067 | 120067600 |
| chr7 | 130884734 | 130966978 |
| chr7 | 131231652 | 131254982 |
| chr7 | 134593095 | 134633317 |
| chr7 | 134658954 | 134693210 |
| chr7 | 137576966 | 137592568 |
| chr7 | 139647589 | 139693738 |
| chr7 | 141038963 | 141049032 |
| chr7 | 152988080 | 152991241 |
| chr7 | 155188380 | 155215009 |
| chr8 | 1743781 | 1760044 |
| chr8 | 6526544 | 6620677 |
| chr8 | 8269124 | 8314665 |
| chr8 | 8330165 | 8408252 |
| chr8 | 11853072 | 11901945 |
| chr8 | 13342717 | 13377904 |
| chr8 | 13436620 | 13486466 |
| chr8 | 16402603 | 16424144 |
| chr8 | 16695346 | 16764972 |
| chr8 | 17069838 | 17092483 |
| chr8 | 17334412 | 17364043 |
| chr8 | 19118686 | 19143141 |
| chr8 | 19166744 | 19251355 |
| chr8 | 23055876 | 23084083 |
| chr8 | 23296523 | 23367671 |
| chr8 | 23497829 | 23561900 |
| chr8 | 23744439 | 23765899 |
| chr8 | 23781035 | 23810242 |
| chr8 | 24192296 | 24235095 |
| chr8 | 25178998 | 25279555 |
| chr8 | 26607793 | 26643624 |
| chr8 | 29626826 | 29676458 |
| chr8 | 30384671 | 30433527 |
| chr8 | 32317813 | 32363153 |
| chr8 | 38985476 | 39043513 |
| chr8 | 42847653 | 42870300 |
| chr8 | 43250504 | 43256259 |
| chr8 | 43494293 | 43501361 |
| chr8 | 48266602 | 48325060 |
| chr8 | 48397908 | 48434555 |
| chr8 | 48569727 | 48596547 |
| chr8 | 48888202 | 48925809 |
| chr8 | 49794156 | 49819877 |
| chr8 | 54136925 | 54173105 |
| chr8 | 56873747 | 56893278 |
| chr8 | 58191175 | 58198279 |
| chr8 | 58827171 | 58850937 |
| chr8 | 61744784 | 61772388 |
| chr8 | 66112620 | 66139782 |
| chr8 | 66439102 | 66464693 |
| chr8 | 71828097 | 71867495 |

| | | |
|---|---|---|
| chr8 | 76953717 | 76957235 |
| chr8 | 79783614 | 79798444 |
| chr8 | 81210288 | 81235703 |
| chr8 | 81339457 | 81370474 |
| chr8 | 83107697 | 83145254 |
| chr8 | 88101299 | 88125428 |
| chr8 | 89754631 | 89845477 |
| chr8 | 89885625 | 89904759 |
| chr8 | 93836846 | 93847492 |
| chr8 | 93987150 | 94044768 |
| chr8 | 94211898 | 94263349 |
| chr8 | 96904860 | 96929498 |
| chr8 | 100867882 | 100909813 |
| chr8 | 106044893 | 106062757 |
| chr8 | 115426429 | 115466330 |
| chr8 | 117697693 | 117732383 |
| chr8 | 117950197 | 118026421 |
| chr8 | 118878080 | 118915228 |
| chr8 | 119029853 | 119083383 |
| chr8 | 122103524 | 122185612 |
| chr8 | 122256355 | 122270615 |
| chr8 | 122310622 | 122322197 |
| chr8 | 123516085 | 123548428 |
| chr8 | 125507218 | 125570182 |
| chr8 | 126824742 | 126841558 |
| chr8 | 126934016 | 126965387 |
| chr8 | 127204814 | 127254863 |
| chr8 | 127889459 | 127950591 |
| chr8 | 133568529 | 133585467 |
| chr8 | 133883142 | 133936319 |
| chr8 | 135577811 | 135609777 |
| chr8 | 140579341 | 140599186 |
| chr8 | 141127102 | 141161786 |
| chr9 | 3843038 | 3881234 |
| chr9 | 6537454 | 6569483 |
| chr9 | 12747473 | 12825692 |
| chr9 | 15997993 | 16065503 |
| chr9 | 21533433 | 21560934 |
| chr9 | 21586510 | 21620567 |
| chr9 | 21672252 | 21690717 |
| chr9 | 21829402 | 21838837 |
| chr9 | 22138552 | 22172647 |
| chr9 | 27108869 | 27131474 |
| chr9 | 32393604 | 32431676 |
| chr9 | 37929488 | 37977632 |
| chr9 | 37993738 | 38068074 |
| chr9 | 70403379 | 70440700 |
| chr9 | 70553395 | 70610361 |
| chr9 | 72464839 | 72479373 |
| chr9 | 73115459 | 73155779 |
| chr9 | 82016906 | 82036378 |

| | | |
|---|---|---|
| chr9 | 89135861 | 89160691 |
| chr9 | 91037222 | 91086185 |
| chr9 | 91161469 | 91200741 |
| chr9 | 93091992 | 93160342 |
| chr9 | 98355571 | 98380742 |
| chr9 | 99415530 | 99446886 |
| chr9 | 100411793 | 100444815 |
| chr9 | 104862927 | 104881543 |
| chr9 | 105026072 | 105074261 |
| chr9 | 106860746 | 106898132 |
| chr9 | 109765825 | 109821275 |
| chr9 | 109877541 | 109914614 |
| chr9 | 110013580 | 110033875 |
| chr9 | 110047658 | 110074638 |
| chr9 | 110095807 | 110153729 |
| chr9 | 110623081 | 110661234 |
| chr9 | 111944799 | 111969870 |
| chr9 | 111992913 | 112076338 |
| chr9 | 113183248 | 113191167 |
| chr9 | 113619843 | 113659429 |
| chr9 | 114879633 | 114910191 |
| chr9 | 115104994 | 115171583 |
| chr9 | 115189445 | 115215287 |
| chr9 | 115590230 | 115618824 |
| chr9 | 115638176 | 115696031 |
| chr9 | 115937857 | 115957618 |
| chr9 | 116058022 | 116107326 |
| chr9 | 125626014 | 125683738 |
| chr9 | 127517492 | 127603256 |
| chr9 | 131983812 | 132009919 |
| chrX | 13460441 | 13490486 |
| chrX | 29657796 | 29667159 |
| chrX | 45696294 | 45720469 |
| chrX | 45756658 | 45810400 |
